# Next generation Human Neocortical Neurosolver model with biophysically realistic somatic, dendritic, and synaptic dynamics

**DOI:** 10.64898/2026.10.29.762964

**Authors:** Katharina Duecker, Austin Soplata, Nicholas Tolley, Sarah Pugliese, Taylor A. E. Jones, Chetan Kandpal, Darcy A. Waller, Dylan S. Daniels, Stephanie R. Jones

## Abstract

The Human Neocortical Neurosolver (HNN) is an open-source biophysical modeling tool for multiscale interpretation of human electroencephalography (EEG) and magnetoencephalography (MEG) signals. HNN’s foundation is a pre-tuned canonical neocortical circuit model consisting of excitatory pyramidal neurons and inhibitory interneurons in supra-(L2/3) and infragranular (L5) layers. This model was originally developed based on studies in rodent and non-human primate cortices when dynamics of human neurons were poorly characterized. Yet recent advances in recordings from human tissue now provide unprecedented detail to update the model. Here, we present a significantly updated model for use in the HNN software, with electrophysiological properties and synaptic connections tuned to human tissue. Beyond reproducing biologically realistic somatic dynamics, the pyramidal cells simulate dendritic calcium (Ca^2+^) spikes in the L5 extratelencephalic-projecting (ET) neurons, and shorter dendritic spikes in the L2/3 pyramidal neurons. We demonstrate how the updated circuitry and new features in the HNN software can be used to disentangle somatic, synaptic, and dendritic contributions to human E-/MEG signatures. The model is distributed with workflows to simulate event-related responses and oscillations, and tutorials demonstrating how membrane potentials, Ca^2+^ dynamics, and synaptic currents can be recorded and visualized. These enhancements allow extensive model validations based on in vitro and in vivo electrophysiology, voltage and Ca^2+^ imaging, and provide users with novel insights on the mechanisms underlying E-/MEG signals.

## Introduction

Brain research is revolutionized by major advances in invasive recording and stimulation techniques in animal models [1], [2], [3]. However, understanding human perception, cognition, and neuropsychiatric conditions requires research in human participants [4], [5]. In humans, non-invasive recordings with electroencephalography (EEG) and magnetoencephalography (MEG) are widely-used for studying the neural correlates of cognitive processes [6] and for identifying biomarkers [7], [8](Fig. 1a). Like many scientific disciplines, human E-/MEG research is transformed by increased data sharing efforts and the development of sophisticated analysis techniques based on machine learning [9], [10], [11], [12], [13]. While translating discoveries from these macroscale signals to the microcircuit dynamics studied in animal models remains challenging, this translation is critical to harness the full potential of E-/MEG for understanding cognition and for diagnosing and treating neuropathology.

**Fig. 1.**
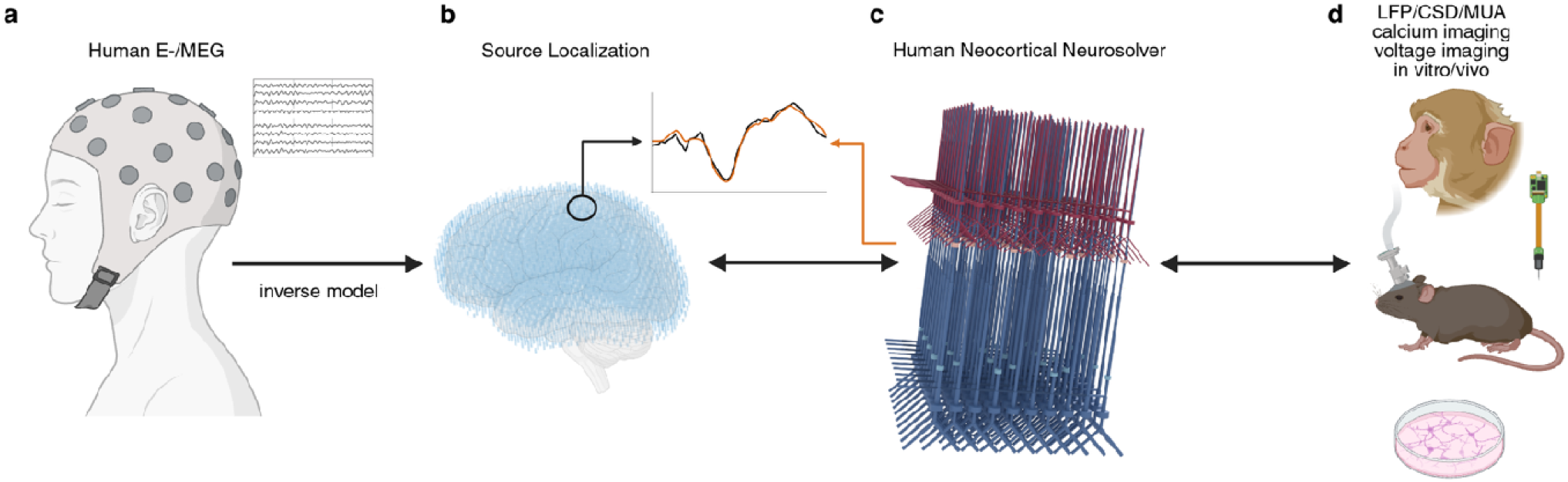
The Human Neocortical Neurosolver (HNN) is a biophysically detailed modeling framework for translation between source-localized human E-/MEG data and mesoscale and microscale dynamics observed in animal models. **a** E-/MEG sensor signals can be projected into **b** source space using inverse modeling techniques, to examine the equivalent current dipole signals within regions of interest. **c** Using HNN, the micro- and mesoscale generators of these dipolar signals can be studied by simulating cell and circuit activity alongside current dipoles that can be directly compared to the source-localized signal in a region of interest. **d** HNN’s predictions can be constrained and validated based on multiscale invasive in vivo and in vitro data. As such, HNN is designed to bridge across scales, from microscale cell and circuit properties to non-invasive human E-/MEG signals. Parts of figure created in biorender.com.

Even though E-/MEG signals are poorly understood in terms of their underlying microcircuit dynamics, it is well understood that their electromagnetic generators can be estimated as equivalent current dipoles, also referred to as primary currents [14], [15], [16](Fig. 1b). These primary currents have been shown to be created by the synchronous postsynaptic currents in the spatially aligned apical dendrites of neocortical pyramidal neurons [14], [16], [17], [18], [19], [20](Box 1). The location, orientation and time course of these primary currents can be estimated with inverse modeling (source localization, Fig. 1b) [14], [21], [22]. While inverse modeling is useful for inferring time and frequency-domain signal dynamics in cortical [23], [24], [25] and occasionally subcortical [26], [27] structures, it does not provide information about the underlying microcircuit-level generators [16].

Computational neural models are powerful tools to address this gap, as they can bridge across scales from human macroscale signals to microcircuit information accessible in animal models (Fig. 1c, d). The Human Neocortical Neurosolver (HNN; hnn.brown.edu) is a computational modeling software designed for interpreting source-localized E-/MEG signals in terms of their multiscale cell and circuit generators (Fig. 1, Box 1) [28], [29]. HNN can be used through a graphical user interface (GUI) or a Python application programming interface (API) that are distributed with tutorials. HNN is constructed with a modular design that is continuously being adapted to allow flexible model expansions based on new insights from experiments or the literature. Box 1 provides a detailed description of the biophysical structure of HNN’s neocortical model and the process for activating the network to develop and test predictions on the multiscale generators of E/MEG signals.

The original model was based on studies from somatosensory cortex [30], [31], but its canonical structure has proven useful for studying neocortical signals from auditory [32], [33], [34], [35], motor [36], [37], and frontal [38], [39] areas, that only required slight adaptations of the original parameters. Despite its success, the model was developed when the biophysics of human neocortical neurons were still poorly characterized, and its cellular and synaptic properties were therefore largely constrained by animal data and general modeling principles.

Here, we build on *in vitro* data from human tissues to present a significantly updated neocortical model that is openly distributed through the HNN Python API and referred to as the *duecker_ET_model*. This model retains the intermediate level of morphological complexity as in the original neocortical model (Box 1) and its biophysics reproduce a wide array of somatic dynamics and postsynaptic potentials (PSPs) observed in human tissue in vitro [40], [41], [42], [43], [44], [45], [46], [47]. Additionally, the L2/3 and L5 pyramidal neurons generate dendritic Ca^2+^ spikes, which were not constrained in previous versions of the model [29], [30], [34], [48].

With corresponding adjustments to the parameters defining the exogenous drive patterns activating the network, the *duecker_ET_model* reproduces the sensory evoked responses and low frequency brain rhythms studied with the original model [29](Box 1). Moreover, simultaneous visualization of membrane potentials, Ca^2+^ dynamics, and synaptic currents, gives users direct access to the microscale dynamics underlying each of these signals. Disentangling these multifaceted contributions reveals that E-/MEG waveforms reflect nuanced cell and network processes. To our knowledge, this is the first biophysical E-/MEG modeling framework to incorporate this breadth of human cell and circuit physiology and dendritic dynamics.

### Box 1: Overview of HNN and process for developing and testing hypotheses on the multiscale cell and circuit origin of human E-/MEG signals

#### HNN simulates the multiscale biophysical generators of E-/MEG

HNN is built around a biophysically detailed model representing a patch of neocortex designed study the cell and circuit generators of source-localized primary currents (i.e. equivalent dipoles)[29], [30]. These primary currents emerge from synchronous longitudinal currents to current in the spatially aligned dendrites in large populations of neocortical pyramidal neurons and are associated with extracellular electric and magnetic fields (Box 1 Fig. 1a) [14], [16], [18], [19], [20]. Locally, the return currents flowing through the extracellular space are associated with the Local Field Potential (LFP)[17]. At longer distances, the volume-conducted fields propagate outside of the head and can be recorded non-invasively at the scalp with EEG, and the associated magnetic field with MEG (Box 1 Fig. 1b,c)[15], [17]. Source localization methods, such as beamformers and minimum norm estimation, invert the signals recorded by EEG and MEG sensors to estimate the primary currents in units of current over distance (nano-Ampere-meters, nAm)[15], [24], [25], [49], [50], [51]. HNN simulates these primary currents from the intracellular current flow in the pyramidal neuron dendrites in the same nAm units, enabling one-to-one comparison between simulated and recorded current source signals (Fig. 1). The dendrites of interneurons are not spatially aligned, rendering direct contributions to E-/MEG signals (and LFP) negligible ([14], [20], [52] but see [17] for counter arguments). Nonetheless, through the inhibitory modulation of pyramidal neuron activity, inhibitory neurons critically contribute to the network dynamics and are therefore included in the model as single-compartment neurons.

#### Neocortical model architecture enables multiscale interpretation and flexible expansion

HNN’s neocortical model has a template architecture consisting of supra- and infragranular layers with multi-compartment pyramidal neurons and single compartment inhibitory interneurons (Box 1 Fig. 1d,e) [29], [30], [31]. The pyramidal neuron models are designed with a deliberately chosen intermediate level of morphological detail [53] to ensure that the simulations are computationally manageable and the outputs interpretable. By default, the architecture contains 100 pyramidal neurons and 35 interneurons per layer (270 neurons by default) but can be scaled for added or reduced complexity (Box 1 Fig. 1d). To account for thousands of synchronous pyramidal neurons known to contribute to E-/MEG signals [15], [20], a *scaling factor* is applied to the simulated equivalent current dipole to match the amplitude of the recorded dipole and provides an estimate of the size of the network that generated the recorded signal [29], [30]. The biophysical details, including ion channel conductance and synaptic weights, are pre-tuned based on empirical data.

#### Thalamocortical and cortico-cortical activation pathways

Simulation experiments in HNN begin by activating HNN’s neocortical column model with simulated trains of action potentials (drives) targeting the cells through excitatory synapses in layer-specific pathways [29], [31], [31](Box 1 Fig. 1f). One pathway, referred to as the ‘proximal drive’ represents input from the lemniscal thalamus targeting the granular layer (not explicitly modeled) which subsequently activates the inhibitory neurons and the basal dendrites of the pyramidal neurons in L2/3 and L5. In HNN simulations of the event-related potential (ERP), the proximal drive to L5 is conventionally weaker and delayed compared to the drive to L2/3, in line with the repeatedly reported sequence of inputs from L4 to L2/3 to L5 during a stimulus-evoked response [31], [54], [55], [56], [57]. However, users can adjust the timing, strengths, and synaptic delays of the proximal drive according to their research questions, for instance, to model direct thalamic input to the infragranular layers [58], [59]. The second pathway, referred to as the ‘distal drive’, represents inputs from non-lemniscal thalamus or higher order cortex and arrives at the distal apical tuft compartment of the pyramidal neurons, and the L2/3 interneuron [60], [61] (Box 1 Fig. 1f). With this construction, in any simulation experiment, users can inspect cell type specific spiking, membrane dynamics including ionic and synaptic currents in somatic and dendritic compartments, Local Field Potentials (LFPs) and Current Source Density (CSD), alongside their E-/MEG signature of interest.

##### Process for developing and testing predictions with HNN

As with any large-scale neural modeling framework, not all parameters can be estimated at once. Most of HNN’s pre-tuned parameters remain fixed when beginning a simulation experiment and only a limited set of target parameters is adjusted to examine the multiscale generation of a particular experimental data set. Therefore, in addition to the pre-tuned neocortical column models, HNN is distributed with pre-tuned sequences of thalamocortical and cortico-cortical drives (proximal and distal drives, see above) constructed to simulate ERPs [31] and low-frequency oscillations [29], [30], [39], [48], [62] in prior studies. This pre-tuned framework is used as the starting point for any simulation experiments. The following outlines the steps and recommended initial set of target parameters (i.e. scaling factor and drive parameters) to adjust when fitting the model to recorded data.

1. Prediction: Starting from a pre-tuned HNN neocortical model and exogenous drive patterns for either ERPs or low frequency oscillations, users first adjust the scaling factor to match the amplitude of the empirical primary current signal. This is followed by adjustments of the parameters describing the strength and timing of the thalamocortical and/or corticocortical inputs. It is recommended that users begin their tuning process manually, guided by HNN’s tutorials and their own hypotheses about the neural generators of their signal of interest. The manually tuned parameters that approximate the general waveform shape can then serve as a starting point for one of the three automated parameter optimization algorithms available in HNN, to find a close fit between the simulated and recorded primary current dipole. Additionally, users can estimate the uniqueness of their parameters using Simulation-Based Inference methods [63](available in Python API). If users find that their parameter adjustments cannot explain the variation in the data, HNN can be used to refine their initial hypotheses or develop new hypotheses through parameter exploration. For instance, users may find that an additional drive is needed [64], or that intra-network connections have to be adjusted to reproduce the waveform shape of the E-/MEG signature [38].
2. Testing: The multiscale signals corresponding to the final waveform shape, including cell voltages, calcium dynamics and LFP/CSD, represent model predictions that can be visualized and quantified. These predictions can be tested through comparison to invasively obtained empirical data from animal models or other human imaging methods under comparable experimental conditions [39].
3. Updating: Due to the modular design of HNN’s core code, new data obtained to test model-derived predictions can be used to further expand and constrain the model [34], [38], [48].

**Box 1 Fig. 1.**
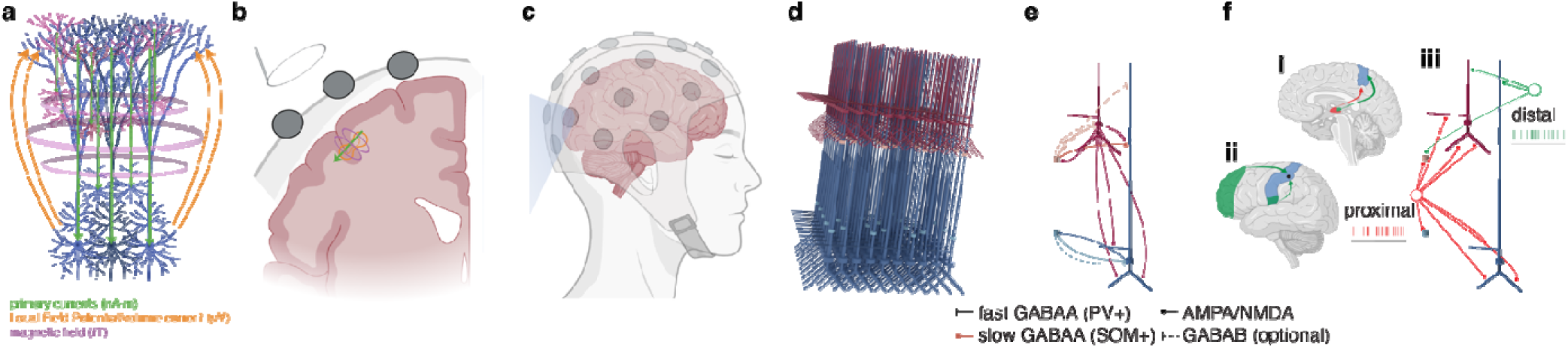
Generators of E-/MEG signals and HNN architecture. **a** Synchronous postsynaptic intracellular currents in the spatially aligned dendrites of thousands of pyramidal neurons (primary currents) and their return currents are associated with electric and magnetic fields. **b, c** These fields can be recorded at longer distances using EEG and MEG. **d** Architecture of HNN models with pyramidal neurons and interneurons in supra- and infragranular layers. HNN was designed to simulate the primary currents in pyramidal neuron dendrites **e** Microcircuit architecture in HNN. Within layers, the single compartment interneuron acts as the PV+ neuron inhibiting the soma of the pyramidal neurons in the same layer through a fast GABAA synapse. The L2/3 interneuron also acts as the SOM+ interneuron for the L5 pyramidal neuron, inhibiting the apical dendrite through a slow GABAA synapse. GABAB synapses can optionally be included but were inactive in the model presented here. **f** The neocortical column model (iii) simulates a small patch of neocortex (blue circle, ii). This patch is activated by a proximal drive reflecting input from sensory thalamus (i, red), and a distal drive conceptualized as connections from non-lemniscal thalamus (i, green), frontal regions, and higher-order sensory regions (ii). Parts of figure created with <u>biorender.com</u>.

## Results

We first describe our updated neocortical column model (named *duecker_ET_model*), including the individual cell models, synaptic dynamics, and connectivity profiles. We show that despite the reduced morphological detail of the pyramidal neurons and interneurons, the model reproduces a myriad of active somatic and dendritic dynamics described in human neurons, along with realistic passive properties. We then outline updates to the network activation parameters needed to reproduce ERPs and low frequency oscillations, consistent with our prior studies.

### Layer 5 pyramidal neuron model reproduces active and passive properties of human L5ET cells

Pyramidal cells in the deep infragranular layer 5, referred to as extratelencephalic-projecting (ET), pyramidal tract (PT), or L5b cells, are well-established generators of complex somatic and dendritic spiking dynamics linked to intracellular Ca2+ [65], [66], [67], [68], [69], [70], [71], [72]. L5ET cells also have the longest apical dendrites of all pyramidal neurons, with a characteristic thick tuft and complex arborizations, both in rodents [59], [73], [74] and in humans [42], [43], [75]. As pyramidal neurons with long apical dendrites have been shown to generate large current dipole moments [20], L5ET a e likely to be major contributors to E-/MEG signals. More recently, their dendritic Ca^2+^ spikes have been proposed to be detectable at the cortical surface [76]. Therefore, all L5 pyramidal neurons were designed to behave as human L5ET cells, allowing us to study how dendritic Ca^2+^ dynamics contribute to E-/MEG signatures of interest (but see Discussion).

The updated L5ET model has the same reduced morphology used in prior HNN models, with 9 compartments as shown in Fig. 2a [29], [31], [53]. Ten active ion channels, adapted from a state-of-the-art model of a L5b rat pyramidal neuron [77], are included in the somatic and dendritic compartments, distributed uniformly (Table 1) and non-uniformly (Supp. Fig. 1b) to reproduce the somatic dynamics observed in human L5ET cells in vitro [43], and dendritic Ca2+ spikes observed in rat neurons [69], [70].

**Fig. 2.**
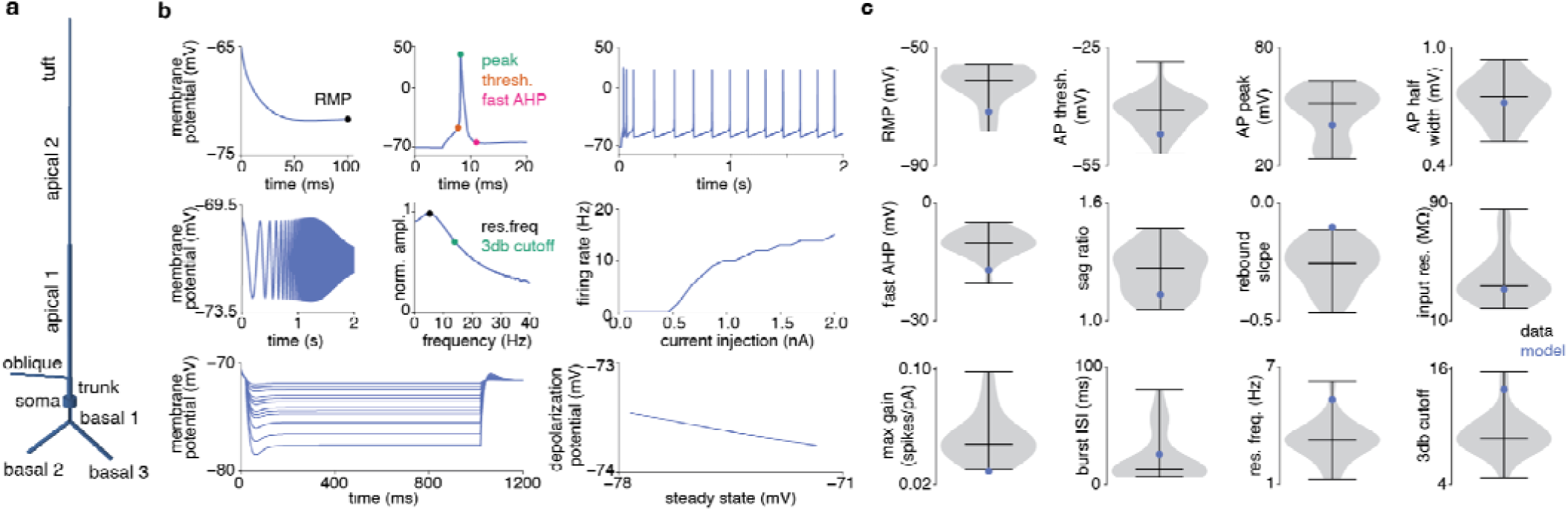
**a** Reduced morphology of the L5ET neuron, preserved from prior HNN models. **b** In vitro experiments from human L5 ET pyramidal cells [43] reproduced in the L5ET cell model. **c** The model’s somatic dynamics (blue) are within a biophysically plausible range for all measures (n=71 human ET neurons, data from [43]). RMP: resting membrane potential, AP: action potential, AHP: afterhyperpolarization, input res.: input resistance, ISI: inter-spike interval, res. freq.: resonance frequency.

**Table 1.** Active conductances in the L5ET model cell. □s in S/cm2; y fraction of free Ca2+, τ decay in ms. Dashes indicate that the ion channel was excluded from the compartment.

|  | soma | basal dendrite | apical dendrite |
|---|---|---|---|
| □.Nat | 1.53 | 0.0102 | 0.01065 |
| □.Kv3.1 | 0.3465 | 0.0001305 | 0.000261 |
| □.Nap | 9.46e-05 | - | - |
| □.SK | 0.6615 | - | 0.0036 |
| □.Ca_HVA | 0.000496 | - | 1.39e-05 |
| □.Ca_LVA | 0.001715 | - | 4.675e-05 |
| □.Ih | 5e-05 | 2.57e-05 | 6e-05 |
| □.Im | 1e-4 | - | 3.375e-05 |
| □.Kp | 0.001115 | - | - |
| □.Kt | 0.0406 | - | - |
| □.pas (1/Rm) | 8.45e-06 | 2.625e-05 | 3.9267e-05 |
| e.pas | -90 | -90 | -85 |
| Ca.τdecay | 460 | 50 | 122 |
| Ca.γ | 0.000501 | 0.0005096 | 0.0005096 |

### Reproduction of somatic dynamics of human L5ET neurons

The human in vitro data were downloaded from the Allen Institute Brain Map (https://brain-map.org/our-research/cell-types-taxonomies/multimodal-characterization-of-individual-neurons#human) and are described in [43]. The maximum conductance values of the ion channels in the model were tuned to reproduce 12 electrophysiological measures recorded at the soma that have been shown to distinguish L5ET cells from intratelencephalic-projecting (IT) neurons [43]. This is demonstrated in the L5ET model simulations mimicking the in vitro experiments conducted in human cells [43](Fig. 2b). The model generates a fast action potential in response to a single pulse, and a burst followed by regular firing in response to a continuous current injection (Fig. 2b top middle and right). Moreover, the active and passive parameters of the cell produce a selective amplification of frequencies at around 5-7 Hz upon injection of a subthreshold chirp stimulus (Fig. 2b, middle panel left). The tuned sodium (Na+) and potassium (K+) conductance ensure gradually increasing firing rates for current injections with increasing amplitudes, and plateau for amplitudes > 1 nA (Fig. 2b middle right). Finally, the density of the I_h_ current reproduces measures from the hyperpolarization experiments conducted in the original study (Fig. 2b, bottom panel). Comparison of the model outputs to each of the electrophysiological measures described in vitro [43](gray violins; n=71 neurons) confirms that the model tuning was successful in that it generates somatic dynamics within a biophysically realistic range (Fig. 2c).

### Reproduction of L5ET dendritic Ca^2+^ spikes

A key goal of this work is to develop a biophysical model for studying the effects of pyramidal neuron dendritic dynamics on E-/MEG signals. Of particular interest are dendritic Ca^2+^ spikes in L5ET cells, which have been linked to the integration of feedforward and feedback inputs [69], [78], perception [79], [80], consciousness [81] and learning [82]. They have also been detected in surface electrodes [76], suggesting that they contribute to the EEG. As detailed experiments of Ca^2+^ spikes in human L5ET neurons are unavailable, we manually tuned the active ion channels in the L5ET cell model based on reports of in vitro experiments from the rat [69], [70] (see Discussion).

First, the maximum conductance values of all ion channels were manually adjusted until the cell generated dendritic Ca^2+^ spikes in response to high-frequency backpropagating action potentials [70](Fig. 3a). In the final model, a sequence of four pulses presented to the soma at 100 Hz leads to an accumulating depolarization at the distal apical dendrite that is associated with a strong influx of Ca^2+^, consistent with the rat L5b experiments (Fig. 3b, middle). Fewer fast action potentials (Fig. 3, top) and current injections presented at an interstimulus interval of 13 ms (Fig. 3b bottom) do not trigger the Ca^2+^ spike, as shown experimentally [70]. As observed in rat L5b cells, the L5ET model cell shows a critical frequency threshold of 95-100 Hz at which Ca^2+^ spikes are generated (cf. Fig. 3c to Fig. 2c, left in [70]). The amount of input current needed to induce a depolarization above 0mV reduces with distance from the soma, again consistent with in vitro experiments (cf. Fig. 3c right Fig. 3f in [83]).

**Fig. 3.**
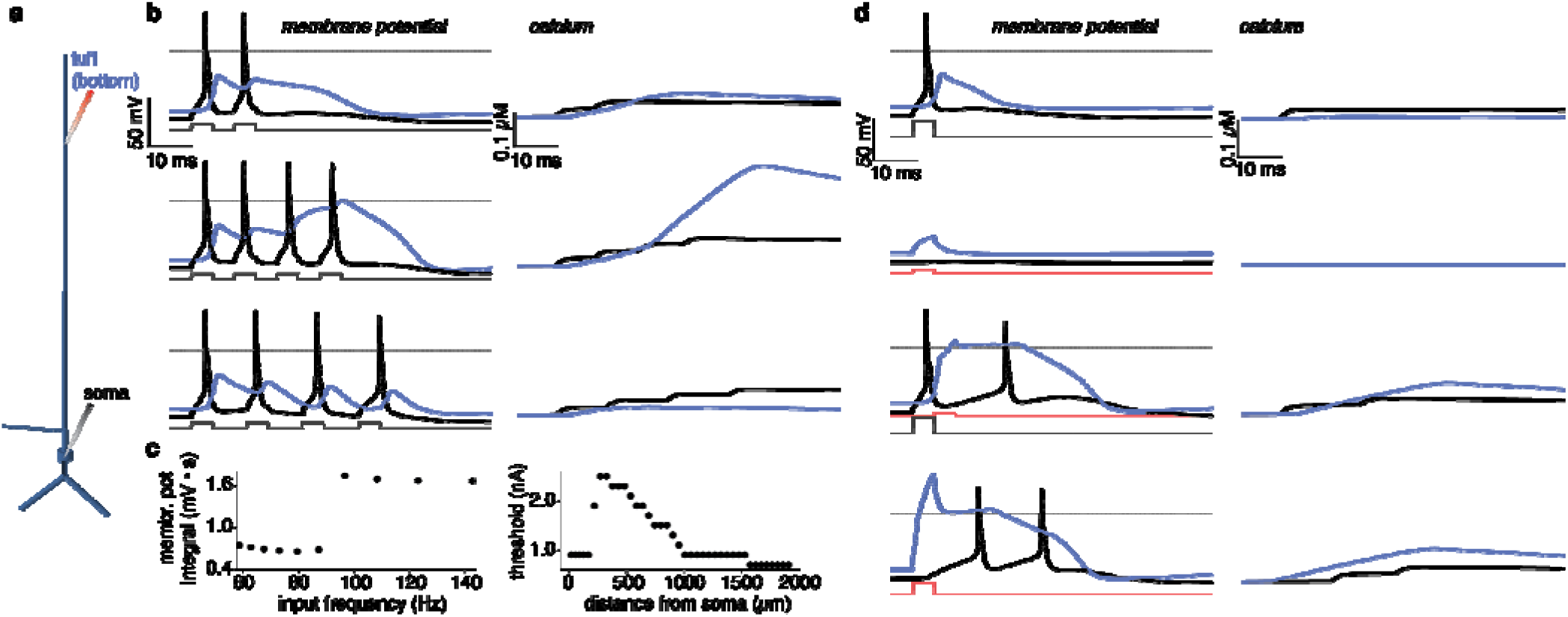
**a** Location of simulated pipettes on L5ET model. Black traces in **b** and **d** denote current injection into soma, red traces represent current injection into the bottom of the tuft, simulating the bifurcation point. **b** Ca^2+^ spikes triggered by high-frequency action potentials at the soma. (top) Two spikes at the soma causing a slight depolarization at the tuft that does not reach the 0mV threshold. (middle) Four action potentials at 100 Hz cause the dendritic membrane potential to reach 0mV, which is associated with a strong influx of Ca2+ into the dendrite. (bottom) Four action potentials with an inter-spike interval of 13 ms (77Hz) do not lead to a Ca^2+^ spike. **c** (left) Membrane potential integral as a function of input frequency demonstrates a critical frequency of >95-100 Hz for inducing a Ca^2+^ spike. (right) The threshold for inducing a depolarization that exceeds 0 mV changes with distance to soma and is lowest in the tuft. **d** Ca^2+^ spikes triggered by coincident somatic and dendritic inputs. (top) A single pulse to the soma triggers an action potential that is attenuated at the tuft. (second from top) a weak pulse to the distal dendrite weakly depolarizes the tuft. (second from bottom) The combination of the two stimuli ignites a Ca^2+^ spike. (bottom) A strong injection to the distal dendrite alone also triggers a Ca^2+^ spike.

Second, all conductance values were tuned to generate dendritic Ca^2+^ spikes upon coincident inputs to the soma and dendrite [69]. To demonstrate this behavior in the final model, we first determined a current injection strength such that a single 5 ms pulse to the soma produces an action potential and a slight depolarization at the bottom of the tuft (Fig. 3 d, top). A weaker input to the apical dendrite creates a small depolarization that does not impact the soma (Fig. 3d, second from top). The combination of these two stimuli ignites a Ca^2+^ spike: a dendritic plateau potential that is associated with an influx of Ca^2+^ into the tuft compartment that causes additional spiking at the soma (Fig. 3d, second from bottom). Additionally, a strong pulse to the distal tuft compartment without somatic input triggers a dendritic Ca2+ spike and somatic bursting (Fig. 3d, bottom), as shown experimentally [69].

### Reproduction of other electrotonic properties

Human L5 pyramidal neurons have been reported to show stronger compartmentalization and signal attenuation between soma and dendrite than their rat counterparts, which is characterized by an exponential increase in input resistance and sag ratio along the apical dendrite [84], [85]. Similar increases in input resistance have been shown for patch-seq characterized human L5ET pyramidal neurons [43]. Indeed, the tuned active and passive properties of the L5ET model cell give rise to an attenuation of signals between soma and dendrite (Supp. Fig. 2a, b) and exponential gradients in input resistance and sag ratio (Supp. Fig. 2c). Of note, the magnitude of the signal attenuation along the dendrite is not as dramatic as shown in previous in vitro experiments [85], which may be due to the reduced morphological detail (see Discussion).

Finally, the dipole moment generated by a single tuned cell is about 0.18 – 0.24 (nA m)/mm2 (normalized to the horizontal reach of the basal dendrites), within range of the dipole moments estimated based on combined human MEG and Electrocorticography (ECoG) recordings (Supp. Fig. 2f) [86].

### L2/3 pyramidal neuron model reproduces active and passive properties of human cells

The L2/3 pyramidal model has the same reduced morphology as in prior HNN models, with 8 compartments as shown in Fig. 4a [29], [30], [31], [53]. Eight active ion channels, adapted from an L2/3 pyramidal mouse model [87] originally taken from [77], Table 2), were distributed uniformly (Table 2) and non-uniformly (Supp. Fig. 1f). The maximum conductance of the channels was adjusted to simulate the dynamic of human L2 and L3 pyramidal cells observed in vitro [88], [89], [90].

**Fig. 4.**
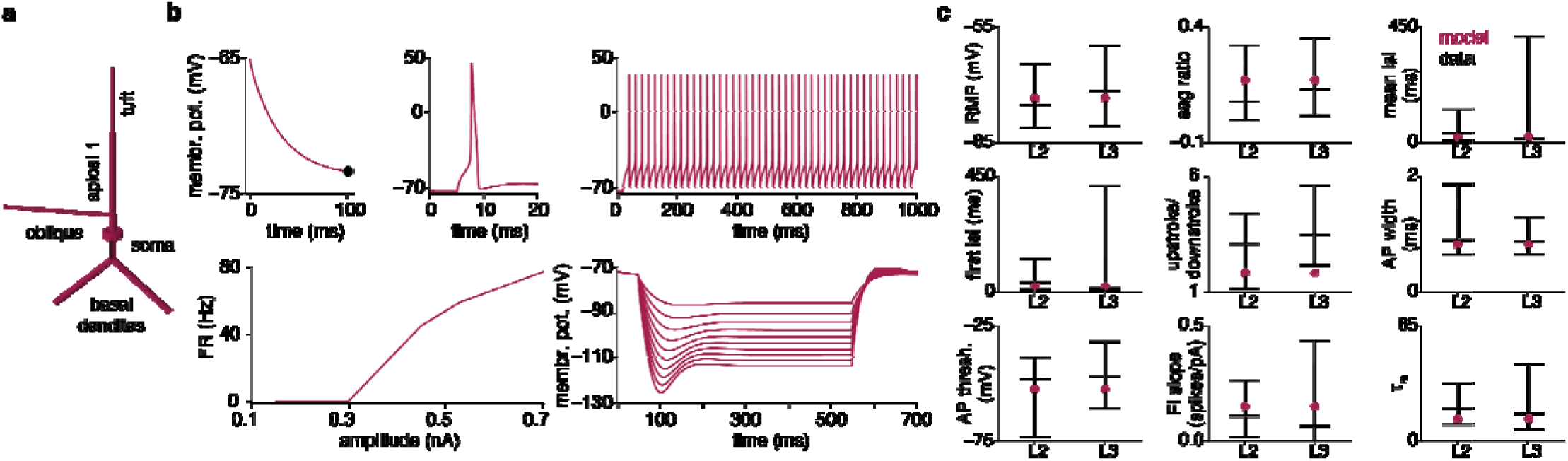
**a** Morphology of the L2/3 pyramidal cell model, consisting of 8 compartments. **b** L2/3 PN model simulations reproducing in vitro experiments from the Allen Cell Types Database. **c** The L2/3 pyramidal neuron model produces dynamics within the biophysically realistic range of L2 and L3 pyramidal neurons. RMP: resting membrane potential, ISI: inter-spike interval, AP: action potential, FI: Firing Rate Input, : membrane time constant.

**Table 2.**
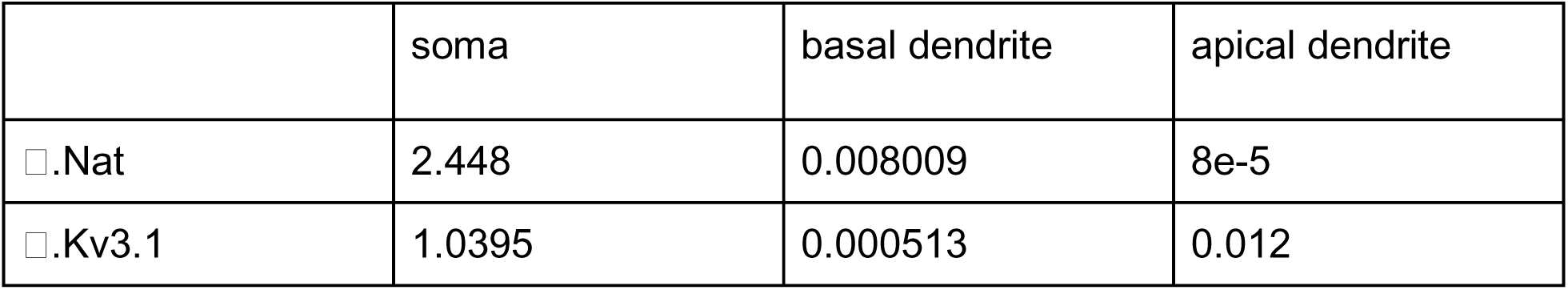
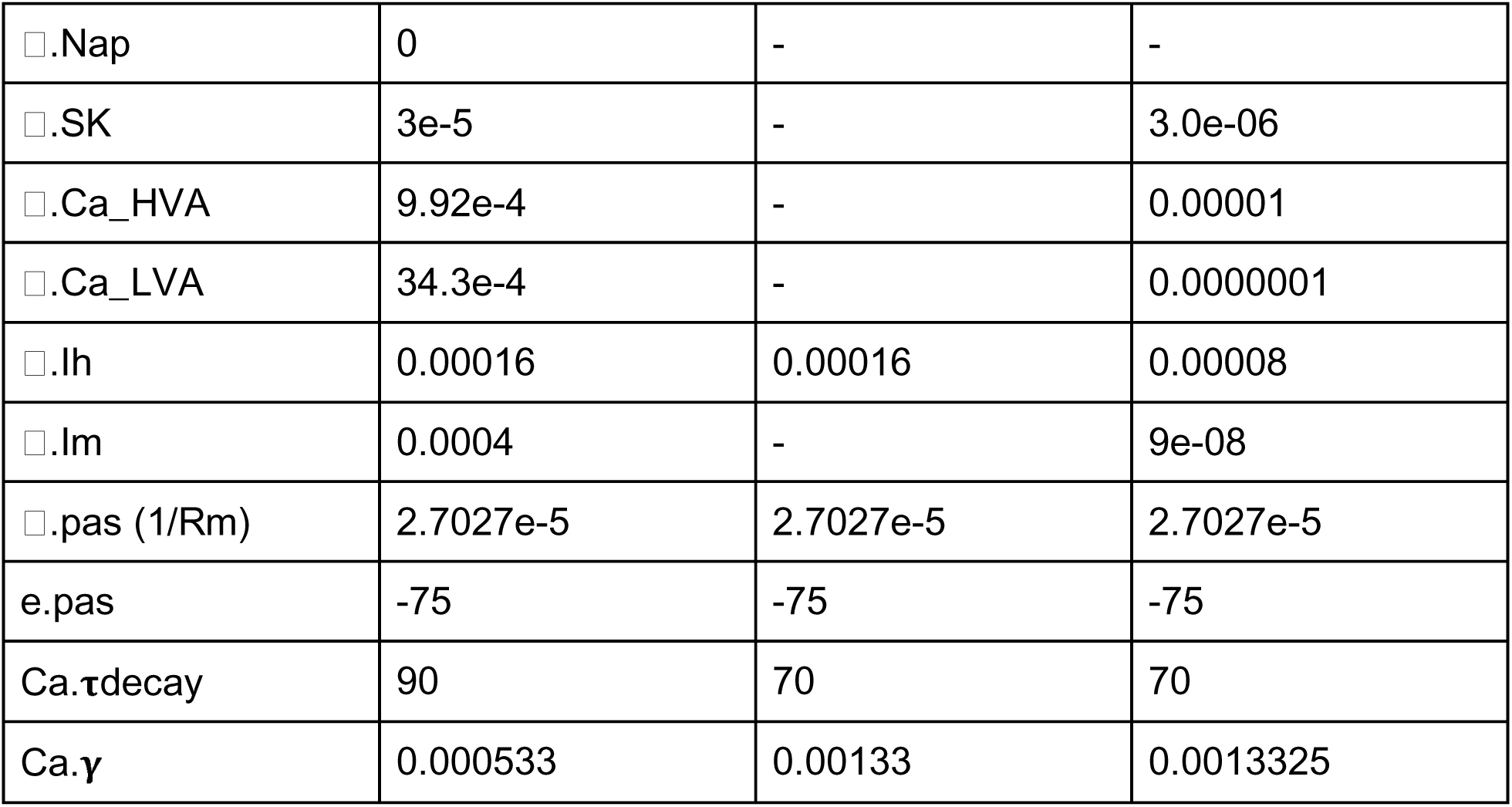
Active conductances in the L2/3 pyramidal cell model. □s in S/cm2; y fraction of free Ca2+, τ decay in ms. Dashes indicate that the ion channel was excluded from the compartment.

|  | soma | basal dendrite | apical dendrite |
|---|---|---|---|
| □.Nat | 2.448 | 0.008009 | 8e-5 |
| □.Kv3.1 | 1.0395 | 0.000513 | 0.012 |
| □.Nap | 0 | - | - |
| □.SK | 3e-5 | - | 3.0e-06 |
| □.Ca_HVA | 9.92e-4 | - | 0.00001 |
| □.Ca_LVA | 34.3e-4 | - | 0.0000001 |
| □.Ih | 0.00016 | 0.00016 | 0.00008 |
| □.Im | 0.0004 | - | 9e-08 |
| □.pas (1/Rm) | 2.7027e-5 | 2.7027e-5 | 2.7027e-5 |
| e.pas | -75 | -75 | -75 |
| Ca.τdecay | 90 | 70 | 70 |
| Ca.γ | 0.000533 | 0.00133 | 0.0013325 |

#### Reproduction of somatic dynamics of human L2/3 pyramidal neurons

All ion channels were tuned to reproduce the somatic dynamics of human L2 (n=19) and L3 (n=179) pyramidal neurons observed in vitro [40], [42], [88](Fig. 4b, c, data downloaded from https://celltypes.brain-map.org/). Additionally, the kinetics of the sodium (Na^+^) channel at the soma were adjusted to produce slower action potentials compared to the L5ET cell (Methods, Fig. 4b). The final model exhibits regular spiking activity triggered by sustained current injection, as reported in human L2 cells [88]. Moreover, the I_h_ channel generates a pronounced voltage sag in response to hyperpolarizing currents [91]. The tuned model produces somatic dynamics within the range of all 9 electrophysiological measures from human neurons, with only the upstroke/downstroke falling marginally below the L3 distribution (Fig. 4c).

#### Reproduction of dendritic Ca^2+^ spikes in human L2/3 pyramidal neurons

The distributions of the I_h_, Ca^2+^, and Ca^2+^-activated potassium (K^+^) channels were tuned to generate comparably brief dendritic spikes upon current injection into the dendrite as observed in vitro ([89], [90] Fig. 5a, Supp. Fig. 1f). In line with the experimental observations, the dendritic spikes generated by the model drive single action potentials at the soma (Fig. 5b) and are associated with a rapid increase in Ca^2+^ at the dendrite (Fig. 5c). Further consistent with in vitro experiments [89], [90] these spikes are not produced by current injection into the soma (Fig. 5d-f). The dendritic spike precedes the somatic action potential by ∼0.5 ms (Fig. 5g; [90]) and has a duration of ∼4.5 ms (Fig. 5g; cf. [89]), matching the temporal signatures of biological L2/3 dendritic spikes. The depolarization caused by the backpropagating action potentials triggered with somatic current injection, on the other hand, have a shorter duration, and lag the action potential at the soma by about 1 ms, again repr ducing previous experimental results (Fig. 5e, g) [89], [90].

**Fig. 5.**
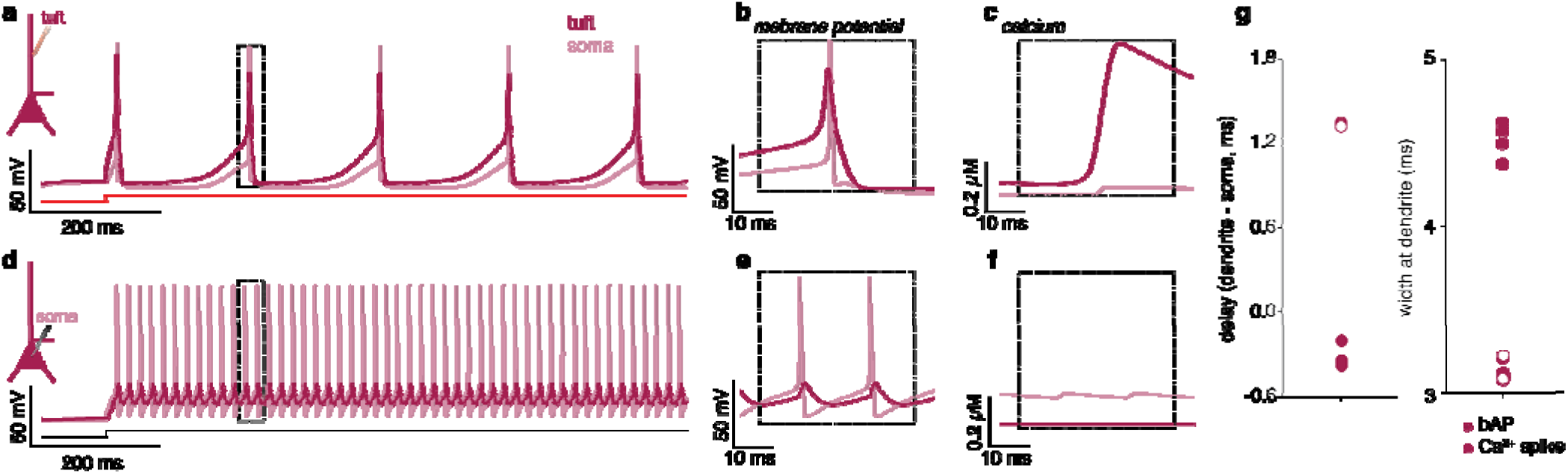
**a** Current injection into the bottom of the tuft compartment of the model cell triggers dendritic spikes. **b** Dendritic spikes lead the spike at the soma and have a shorter duration than Ca^2+^ spikes empirically observed in layer 5 pyramidal cells. **c** Dendritic spikes are associated with strong Ca^2+^ influx into the tuft. **d-f** Continuous current injection into the soma does not trigger these Ca^2+^ spikes. **g** (left) Backpropagating action potentials (e) lead peaks at the dendrite by ∼1.2 ms, while a dendritic Ca2+ spike (b) leads the somatic action potential by ∼0.5 ms. (right) Backpropagating action potentials have a width of about 3 ms when recorded from the tuft, while L2/3 dendritic Ca2+ spikes have a width of about 4.5 ms.

#### Reproduction of additional electrotonic properties of human L2/3 pyramidal neurons

The passive properties of the L2/3 pyramidal model were adjusted to achieve a strong attenuation of depolarizing signals between soma and apical dendrite, as observed experimentally in human L2/3 pyramidal cells ([89], [90] Supp. Fig. 3d,e). Backpropagating action potentials attenuate from soma to dendrite with a length constant of λ = 305 µm (single exponential fit; Supp. Fig. 2d), approximating the 290 µm reported in [89]. The backpropagating action potential upstroke is high near the soma but rapidly reduces to ∼200 mV/ms, the maximum value reported experimentally (cf. Fig. S5A in [89]), and decays exponentially toward zero.

The amplitude of the depolarization induced by current injection into the tuft attenuates with a space constant of about 342 µm, notably larger than the value identified in vitro (154µm, cf. Figure 2C in [90]). This quantitative discrepancy likely reflects the reduced morphological detail of the dendritic tree in our 8-compartment model (see Discussion).

#### Reproduction of equivalent current dipole magnitudes in L2/3 neocortical pyramidal neurons

Lastly, the tuned L2/3 pyramidal neuron model generates a relatively small dipole moment of ∼0.05 nA·mm² (Supp. Fig. 2g), 4-5 times smaller than the dipole generated by the L5 pyramidal neuron, in line with previous *in vitro* work [20] and the original HNN model (Supp. Fig. 3. e, [29]).

### Interneurons reproduce active properties of human inhibitory neuron somatic dynamics

Interneurons are simulated with a single compartment, tuned to in vitro data from human aspiny cells [40], [92]. PV+ and SOM+ cells are simulated with the same ion channel kinetics, and their differences are accounted for by their postsynaptic effects. The same interneuron model was used in L2/3 and L5.

Each interneuron model includes the active ion channels from a previous model of fast-spiking interneurons ([93], [94], Table 3, Methods), together with an I_h_ current that reproduces the sag observed in inhibitory human neurons in vitro [92]. The maximum conductance of each ion channel (Table 3) was tuned to reproduce somatic dynamics observed in human aspiny cells in vitro (n = 79, Fig. 6, data downloaded from data downloaded from https://celltypes.brain-map.org/), focusing on three objectives: (1) short-duration action potentials (< 1 ms, Fig. 6a, b), (2) fast-spiking dynamics in response to sustained current injection, and (3) a steeply increasing firing rate with increasing current injection. Mimicking the in vitro hyperpolarization experiments [92], the I_h_ conductance produces a small sag ratio relative to the pyramidal neurons, together with a fast membrane time constant. The tuned interneuron model replicates all electrophysiological measures within the range of the biological data (Fig. 6a, b, bottom right, respectively).

**Fig. 6.**
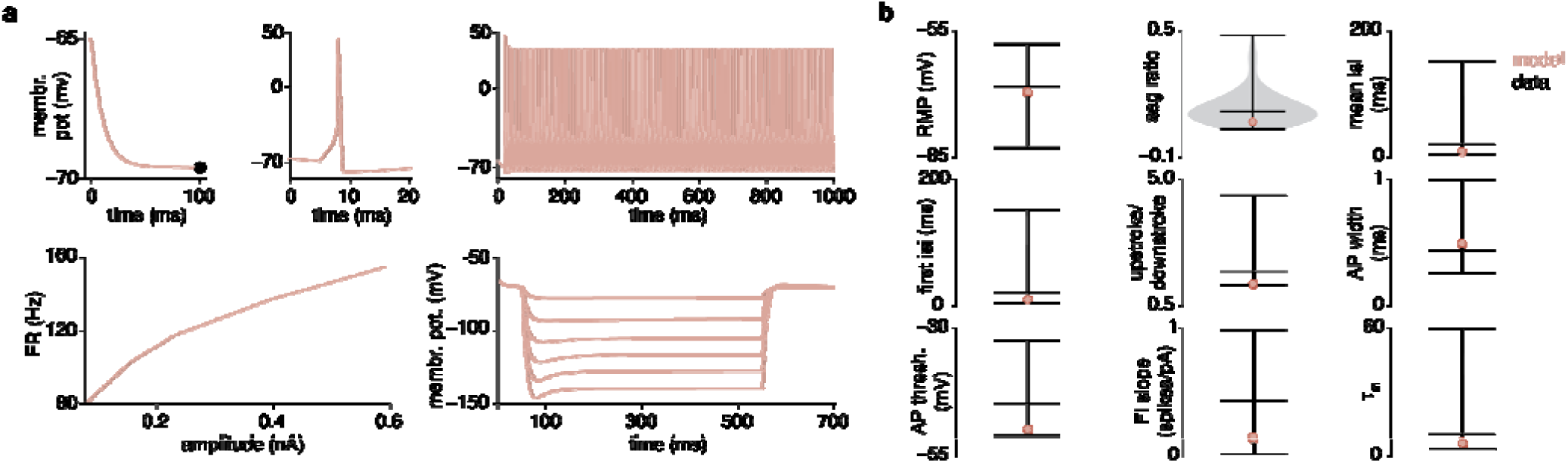
Generic single compartment interneuron model reproduces somatic dynamics of aspiny human cells. **a** In vitro experiments from the cell brain atlas reproduced in the model cell. **b** Comparison of the electrophysiological measures of the model to the human aspiny cell (n=79).

**Table 3.**
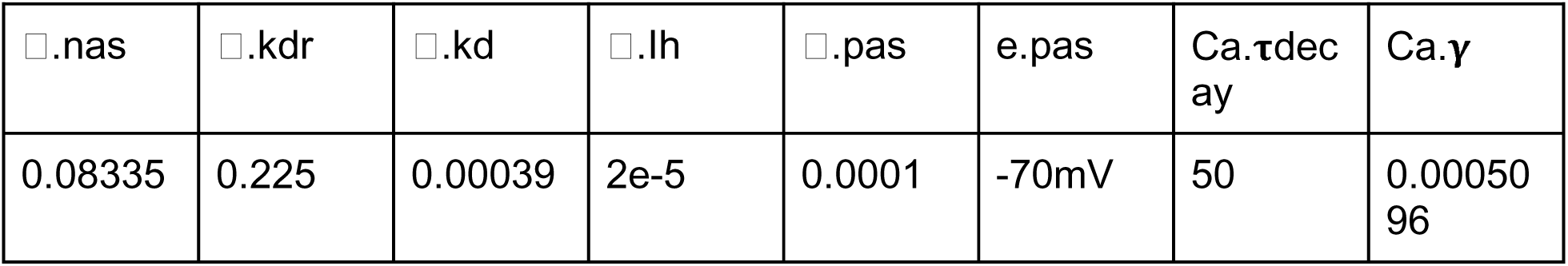
Active maximum conductance values □ and Ca2+ dynamics in the interneuron model. □s in S/cm2; y fraction of free Ca2+,τdecay in ms.

| □.nas | □.kdr | □.kd | □.lh | □.pas | e.pas | Ca.τdecay | Ca.γ |
|---|---|---|---|---|---|---|---|
| 0.08335 | 0.225 | 0.00039 | 2e-5 | 0.0001 | -70mV | 50 | 0.0005096 |

### Biologically realistic synapse models reproduce the balance of excitatory Ca^2+^ currents by dendritic inhibition

The AMPA and peri-somatic GABAA synapses are simulated using the double exponential formalism (i.e., *exp2syn* in NEURON) used for all synapses in the original HNN model [29], [30], while several synaptic connections were updated for further biological realism: (1) the NMDA synapses are modeled with a biologically realistic model incorporating synaptic Ca^2+^ currents and magnesium (Mg^2+^) block [95], [96], [97], [98]; (2) the *exp2syn* GABAA synapse at the L5 pyramidal apical dendrite was updated to match the slower kinetics of SOM+-mediated dendritic inhibition [99]; and (3) the GABAB synapse is modeled with a biologically realistic formalism including second messenger connectivity to a G protein-coupled inwardly-rectifying potassium channel [100], [101].

#### Biologically realistic NMDA synapse couples synaptic input to intracellular Ca^2+^

The updated NMDA synapse is based on the model in Gao et al. [95], [98] that was constructed to reproduce both the fast-rise/slow-decay kinetics of empirically recorded NMDA receptors [102] as well as a synaptic Ca^2+^ current and a voltage-dependent Mg²⁺ block [96], [97] (Fig. 7a). Replacing the original HNN NMDA synapse with this biologically realistic model results in a longer postsynaptic depolarization (Fig. 7b vs c). Critically, the NMDA synapse also produces a measurable increase in postsynaptic Ca2+ induced directly from the synaptic current (Fig. 7d, red), which is not generated by *exp2syn* (Fig. 7d, black). The Mg²⁺ block further produces a voltage-dependent increase in current flow, in contrast with the voltage-dependent decrease in driving force observed in the *exp2syn* model (Fig. 7e).

**Fig. 7.**
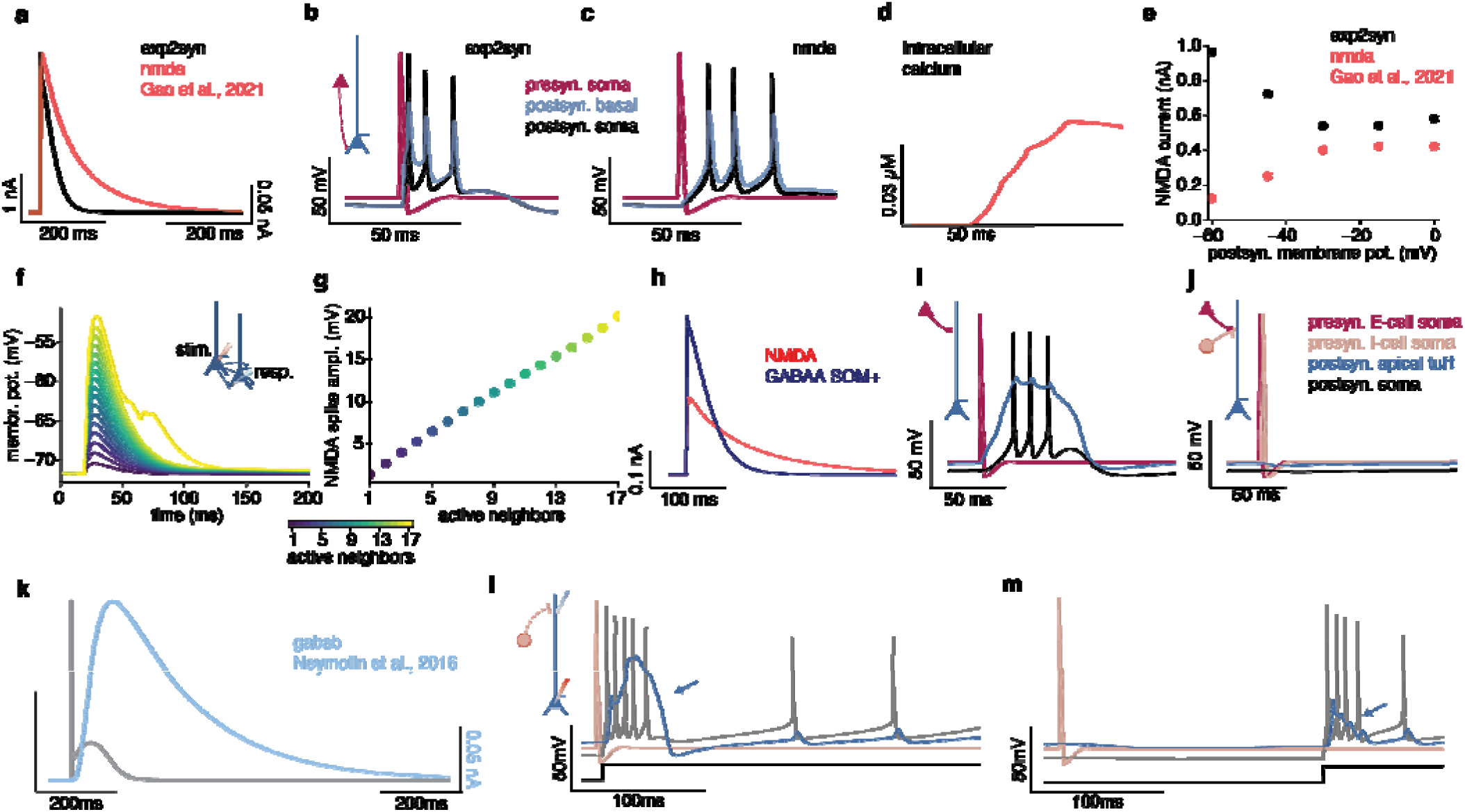
Biologically realistic synaptic dynamics in the duecker_ET_model. **a-e** The NMDA synapse simulates a synaptic Ca^2+^ current and a Mg^2+^ observed in biological NMDA synapses. **a** Comparison of the updated NMDA synap e to the double exponential function used in previous HNN models. **b,c** Postsynaptic potentials in the basal dendrites and resulting spiking dynamics for the different types of synapses. **d** The updated NMDA synapse causes an influx of Ca^2+^ into the compartment where the synapse is placed (red line) which is not modeled by the double exponential function (black line). **e** For the exp2syn formalism, the peak current flowing through the synapse decreases with an increase in membrane voltage at the post-synapse, reflecting a reduction in driving force as the membrane approaches the synapse’s reversal potential. The biophysically realistic NMDA synapse models a Mg^2+^ block, which release as the membrane potential increases, causing an increase in maximum current flow through the synapse with increased depolarization at the post synapse. **f,g** Triggering a single spike in an increasing number of neighboring neurons leads to an increase in the amplitude of the postsynaptic potential at the basal 2 compartment of a reference neuron. **g** The amplitude of the postsynaptic potential increases with an increasing number of active neighboring cells. **h** Inhibition of the L5 dendrite by SOM+ cells is modeled by an exp2syn connection between the L2/3 interneuron and L5ET apical dendrite, whose temporal dynamics oppose the NMDA synapse. **i** Excitation of the L5ET apical dendrite by a neighboring cell causes a Ca^2+^ spike leading to spiking at the soma. **j** This Ca^2+^ spike is suppressed by spiking of the L2/3 interneuron. **k** Optional GABAB synapse with slower rise and decay dynamics than the previously used exp2syn. **l** An inhibitory neuron connected to the apical dendrite through a GABAB synapse fails to suppress the Ca2+ spike when firing shortly before its onset. **m** The Ca2+ spike is suppressed if the inhibitory neuron fires 200 ms before its onset.

NMDA receptors can generate spike-like events in the basal and distal apical tuft dendrites of pyramidal neurons, so-called NMDA spikes [67], [103]. To test whether the NMDA receptors on the L5ET model can reproduce this behavior, we built a network of 100 L5ET cells, recurrently connected at the basal 2, basal 3, and apical oblique compartments through AMPA and NMDA receptors, as in the final *duecker_ET_mode*l (synaptic weights below; Fig. 7f inset). The NMDA spike was quantified based on the membrane potential was recorded at the basal_2 compartment of the center neuron. To increase the input to that center neurons, we triggered single action potentials in an increasing number of neighboring cells. As NMDA spikes have been shown to be independent from voltage-gated Na^+^ and Ca^2+^ channels [67] we recorded the membrane potential in the basal 2 compartment of the reference neuron, which does not contain Ca^2+^ channels and only a low density of Na^+^ channels (Supp. Fig. 1). The model reproduces the qualitative shape of experimentally observed NMDA spikes, but is 50–100 ms shorter in duration [67], [103]. Its amplitude also grows linearly with the incoming input, rather than in the stepwise manner reported experimentally (Fig. 7g) [103].

#### PV+ and SOM+ interneurons distinguished by time-constant and location of postsynaptic GABAA inhibition onto pyramidal neurons

As described above, the model contains single compartment interneurons constructed to represent PV+ and SOM+ interneurons, simulated with the same intrinsic ionic current kinetics (Table 3). The model types are differentiated by the time-constant and location of the postsynaptic connection onto the pyramidal neurons. A fast GABAA synapse at the pyramidal neuron somas (rise,decay 0.5,5 ms), modeled with an *exp2syn* NEURON formalism, represents peri-somatic PV+-mediated inhibition [104], [105], [106]. A slow GABAA synapse at the L5ET apical dendrite represents dendritic SOM+-mediated inhibition, with the rise (1.5ms) and decay time (20ms) specified based on vitro recordings from human tissue [99], [107](Fig. 7h-j).

#### Slow dendritic GABAA inhibition suppresses NMDA-mediated dendritic Ca2+ spikes

GABAA-mediated inhibition of the apical dendrite has been proposed to suppress excitatory inputs and thereby prevent dendritic Ca2+ spikes [99], [107]. We tested whether our model reproduces this behavior using a microcircuit consisting of two Hodgkin-Huxley single-compartment neurons connecting to the apical tuft of the L5ET model. The excitatory point neuron contacts the L5ET neuron through an AMPA and an NMDA synapse, while the inhibitory point neuron contacts the dendrite at the same site through the slow GABAA synapse. The synaptic weights in this experiment were taken from the maximum weights of the final network model (tuned as described below; Fig. 8). Excitatory input alone triggers a Ca^2+^ spike (Fig. 7g, h), which is suppressed by coincident firing of the inhibitory point neuron (Fig. 7i)[108]. This experiment demonstrates how the adjusted time constants of the SOM+-like inhibition, together with the inclusion of a biologically realistic NMDA synapse model, enhance the biophysical realism of the *duecker_ET_model*.

**Fig. 8.**
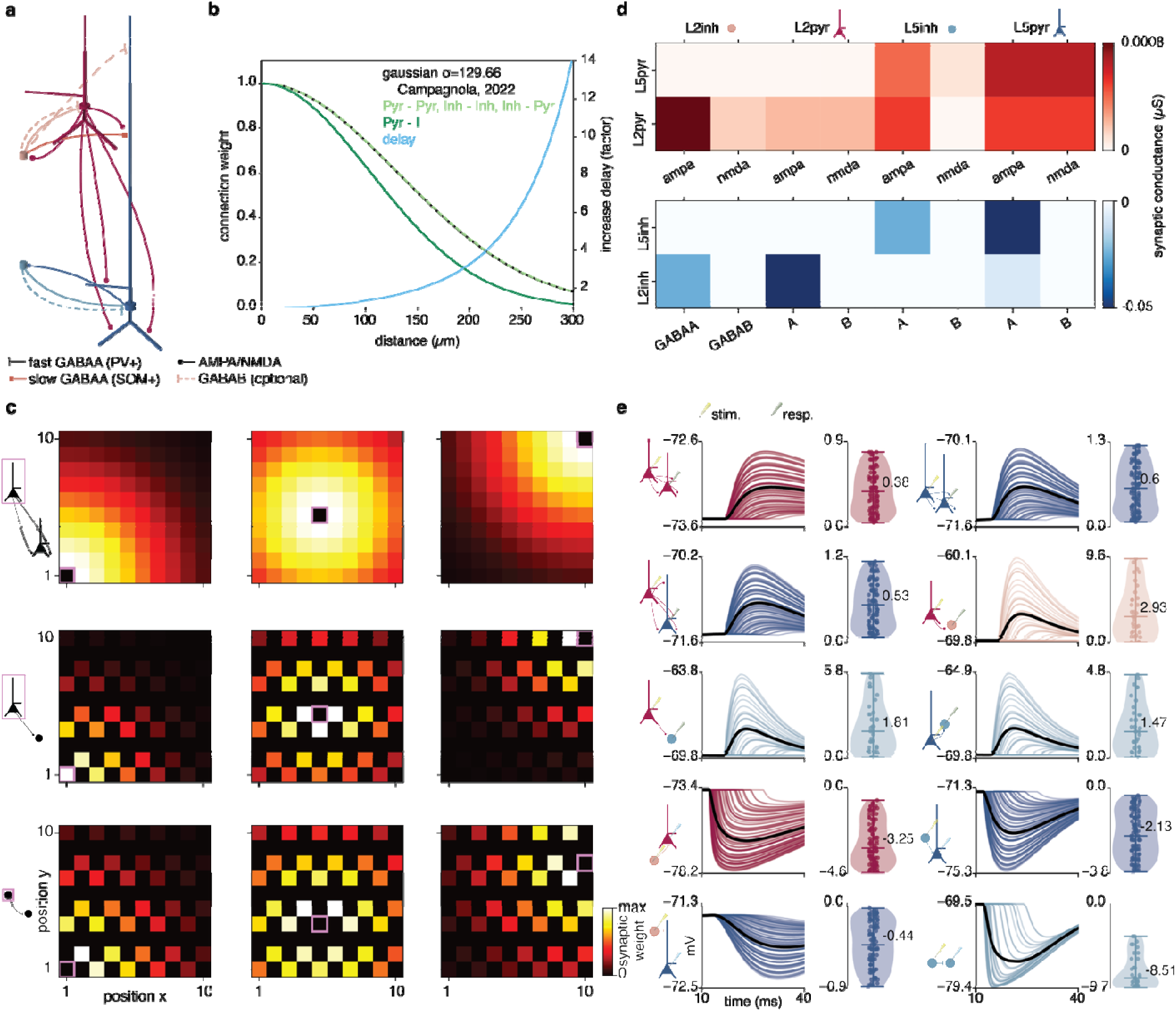
Synaptic architecture and connectivity in the duecker_ET_model. **a** Canonical circuit, consisting of pyramidal neurons and interneurons in supra- and infragranular layers. The pyramidal neurons activate neighboring cells through AMPA and NMDA synapses. Interneurons inhibit the somas of pyramidal neurons in the same layer through a fast GABAA synapse, modeling the effects of fast spiking PV+ interneurons, while the L2/3 interneurons inhibit the apical dendrite of the L5ET pyramidal neuron through a slow GABAA synapse, modeling the effects of SOM+ cells. GABAB at the somas of the pyramidal neurons and the dendrites of the L5 neuron can optionally be added but were not explicitly tuned for this network. **b** Connection strength and synaptic delay as a function of distance between neurons. **c** Heatmaps of the synaptic conductance on the postsynaptic neurons at each location on the grid relative to the location of the pre-synapic neuron in the same layer (pink box). The decay in connection weight demonstrates the effects of the space constant on intralaminar connections between pyramidal neurons (top), pyramidal neurons and interneurons (middle), and interneurons (bottom). For each connection type, an example connection is shown for presynaptic neurons in the bottom left corner (left column), center (middle column) and top right corner (right column). **d** Synaptic weights of the inter- and intralaminar excitatory (top) and inhibitory (bottom) connections. **e** Postsynaptic potentials for all cell type connections. Individual traces show changes in membrane potentials of postsynaptic cells (blue pipette) caused by an action potential of one individual presynaptic cell (red pipette). Violin plots and scatters show the amplitude of the postsynaptic response.

#### GABAB inhibition suppresses Ca^2+^ spikes in a latency-dependent manner

In the fully connected neocortical *duecker_ET_model*, the synaptic weights of the GABAB receptors on all neurons are set to 0 (Fig. 8b), as GABAB receptors are predominantly activated by diffuse release of GABA by inhibitory cells residing in L1 [109], [110], which are not explicitly modeled. However, the synaptic weights of GABAB receptors can optionally be increased to test hypotheses about GABAB inhibition (Fig. 8a). This is different from the original model, which has active GABAB conductances at the soma [29]. Compared to the *exp2syn* formalism used in the original HNN model [29], [30], [31], [48], the updated GABAB synapse has substantially slower rise and decay kinetics and produces a smaller current at equivalent synaptic weights (Fig. 7k).

As a proof-of-concept, we used a simplified circuit to show that the dynamics of the GABAB synapse, when activated at the L5ET apical dendrite, produce realistic dynamics designed to represent inhibition by L1 interneurons [82], [110], [111]. Palmer et al [110] have shown that transcallosal inhibition of L5 apical dendrites, triggered by ipsilateral hindpaw stimulation, was mediated by layer 1 interneurons through GABAB receptors. These inhibitory effects were most effective when the ipsiplateral hindpaw stimulation preceded the contralateral stimulation by 200-400 ms [110], which may be in part attributable to the slower rise kinetics of the GABAB receptor. Likewise, data by Pérez-Garci et al [112] has revealed that the activation of GABAB receptors suppresses the later part of the Ca^2+^ spike in the apical dendrite, when the onset of the inhibition and the Ca^2+^ spike overlap in time. Collectively, these results suggest that inhibition of dendritic Ca^2+^ spikes by GABAB receptor takes several milliseconds to develop and can last hundreds of milliseconds. We reproduce this behavior in a reduced circuit consisting of the L5ET model receiving continuous somatic current injection and an inhibitory point neuron connected to the apical dendrite via the biologically realistic GABAB synapse (Fig. 7l,m). When the current injection to the soma directly follows the action potential of the inhibitory neuron, the Ca^2+^ spike can be observed at the dendrite (Fig. 7l). However, it is suppressed if the current injection lags the spike of the inhibitory cell by 200 ms (Fig. 7m), consistent with [110].

### Local Network Connectivity tuning based on vitro recordings from human tissue

The tuned cells and synapses were combined in the *duecker_ET_model,* preserving the number and ratio of pyramidal and interneuron in each layer of the previous HNN default neocortical column model. We tuned the intra- and interlaminar network connectivity based on studies on human tissue (Fig. 8).

The microcircuit connections between cells of the same and different types are shown in Fig. 8a. Within layers, the inhibitory interneurons act as PV+ cells by inhibiting each other, and the pyramidal neurons at the soma. The L2/3 interneuron also simulates slower, SOM+-like inhibition of the L5ET apical dendrite. Recurrent connections between pyramidal neurons in the same layer activate the basal dendrites and apical obliques through AMPA and NMDA synapses. The L2/3 pyramidal neuron also connects to the L5ET cell at the apical dendrite and the basal dendrites and apical oblique. Pyramidal cells activate interneurons in the same layers through AMPA and (weaker) NMDA receptors.

Consistent with prior HNN network models [29] and experimental work [41], the connection strength between all cells in the model decays with distance between two connected neurons, which is mirrored by an increase in synaptic delay. The standard deviation of the Gaussian distance-dependence was identified from connection probabilities reported in human slice recordings [41](Fig. 8b, c). For pyramidal-to-interneuron connections, the distance-dependent decay is steeper than for pyramidal-to-pyramidal connections to reproduce the increased locality of these connections observed experimentally in rodent tissue [41], [113], [114] (Fig. 8b, c). This profile differs from the original HNN model, which had more local pyramidal connections and wider inhibitory connections (Supp. Fig. 3)[29].

After defining the distance-dependence we manually tuned the maximum synaptic weights to reproduce the distribution of PSPs reported in human tissue (Fig. 8c,e). The PSPs were quantified at the soma of each postsynaptic neuron in response to an action potential in each presynaptic neuron (Fig. 8e), as described in the experimental protocols [44], [45], [46], [47]. Excitatory pyramidal-to-pyramidal connections generate small EPSPs skewed toward zero, matching recordings from human L2/3 [46], [47] Fig. S5 in [41]; also aligning with sparse excitatory connections observed in rodent barrel cortex [115]. Pyramidal-to-interneuron connections within the same layer produce EPSPs of up to 10 mV, consistent with reports of strong excitation onto interneurons [44], [45], as well as observations of strong synaptic connections between excitatory neurons and PV+ interneurons in rodents [116]. SOM+-like inhibitory PSPs at the L5ET apical dendrite from the L2/3 interneuron reach up to 0.9 mV [107]. Intralaminar inhibitory connections, for which human data are currently unavailable, were tuned to prevent excessive spiking activity in the ERP simulation.

The fully connected model, with single cell somatic and dendritic dynamics, synaptic currents, and connectivity tuned as described above, will be referred to as the *duecker_ET_model* in the following.

### Simulating event-related responses and low frequency oscillations in the next-generation HNN

To provide users with a starting point for their own simulations, the *duecker_ET_model*, like its predecessors, is distributed with optimized drive sequences for studying ERPs and low frequency oscillations in the alpha, beta and gamma bands.

#### Reproducing ERPs with a sequence of exogenous drives to the local network

We sought to simulate a tactile evoked response to a brief stimulus source-localized to the primary somatosensory cortex (SI data shown in Fig. 9b [29], [30]. Based on prior studies [31], [55], the SI ERP is simulated by activating the pre-tuned *duecker_ET_model* with a sequence of thalamo-cortical (proximal) and cortico-cortical (distal) drives (Fig. 9a, see Box 1 Fig. 1e). These excitatory drives consist of a pre-defined distribution of spikes that contact the network through AMPA and NMDA synapses. By default, each network cell in the model is activated by one corresponding drive cell, whose spike time within and across trials is drawn from a normal distribution, with a predefined mean and standard deviation. The proximal drive targets all 200 pyramidal neurons and 70 interneurons and consists of 270 cells (Fig. 9a). The distal drive does not target the L5 interneurons and consists of 235 cells [29]. This results in the different network cells being activated at different times (see Supp. Table 1, Fig. 9b), ensuring spike-time variability between cells and trials. The waveform shape of the ERP is reproduced by a sequence of drives consisting of a proximal drive at ∼15 ms [117], representing the first input from thalamus to cortex, followed by a distal drive at ∼59 ms, representing subsequent neocortical-cortical feedback to the local network, followed by a re-emergent thalamic drive at ∼122ms (Supp. Table 1)[29], [31], [55](Fig. 9b, top). The optimized drive parameters (Supp. Table 1, Methods) achieve a close fit to the empirical dipole (RMSE = 10.82, Pearson r=0.99, Fig. 9c, 3 trials).

**Fig. 9.**
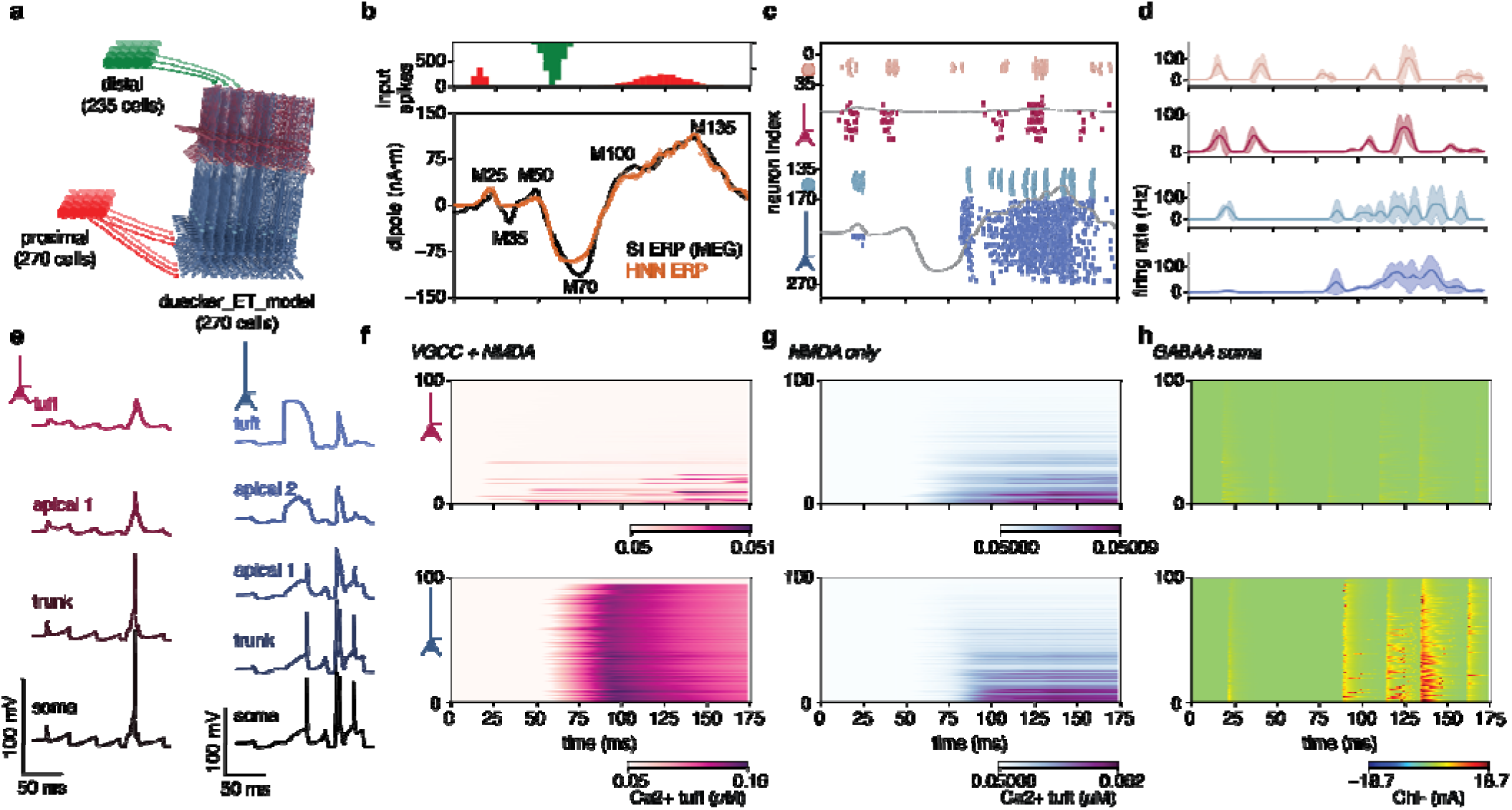
ERP simulations using the duecker_ET_model. **a** Thalamocortical and cortico-cortical inputs to the circuit are modeled by spike trains arriving at the basal dendrites of the pyramidal neurons and somas of the interneurons (the proximal drive) and spikes arriving at the distal apical dendrites and L2/3 interneuron (the distal drive). By default, each network cell is activated by a corresponding drive cell. **b** (top) A sequence of proximal and distal drives is used to simulate the ERP. (bottom) The ERP waveform simulated with the HNN model closely matches the tactile evoked response in SI. **c** Network spiking, overlaid for three trials, and average layer-specific dipoles (dotted lines), show an initial cyclic activity of the L2/3 pyramidal neurons and interneurons, and delayed firing in L5. The distal drive triggers spiking in L5, while the second proximal drive activates predominantly the L5 pyramidal neurons. **d** Stimulus-induced firing rates for each neuron type. Line plots show average over trials shaded area indicates the standard deviation. **e** (left) Membrane potential traces for the L2/3 pyramidal neuron with a median firing rate, showing individual spikes and attenuated backpropagating action potentials. (right) Membrane potential traces for the L5ET neuron with a median firing rate, showing dendritic Ca^2+^ spikes, spiking and bursting. **f** Ca^2+^ concentration at the tuft of the L2/3 (top) and (bottom) L5ET pyramidal neuron resulting from Ca2+ currents through voltage-gated calcium channels (VGCC) and the NMDA synapse. Neurons are ordered based on their distance to the network origin, with neuron 1 being at the center of the network. The L2/3 pyramidal neuron tufts show a relatively small change in dendritic Ca2+ over the course of the trial, while the distal drive triggers an influx of Ca^2+^ into the L5ET dendritic tuft. **g** Ca^2+^ concentration when blocking the VGCCs. This simulation approximates the increase in Ca^2+^ through the NMDA channel alone. **h** Current flow through the GABAA synapse at the soma of the respective pyramidal neurons. Synchronous firing of the L5 interneurons (see **e**) repeatedly leads to strong inhibition of the L5ET somas.

The simulated firing-rate patterns are consistent with recordings from rodent barrel cortex during vibrissae stimulation showing that a strong responses in the L5ET population [79], [80], [118] and sustained firing in L2/3 interneurons over several hundred milliseconds [118]. Furthermore, across cell types, the predicted stimulus-induced firing rates of 50–100 Hz align with stimulus-evoked rates from other sensory areas in non-human primates, including V1 [119] and A1 [120], [121] (Fig. 9e). This is particularly important as previous work using the original HNN model has revealed that the ERP waveform shapes in A1 [34] and frontal regions [38] can be reproduced by adjusting the timing and strength of the sequence of drives used here. This supports the notion that the HNN ERP simulation can be used as a canonical starting point to study evoked responses in other brain areas.

Due to the updated cell physiology and connectivity of the *duecker_ET model*, parameter optimization produced distinct values compared to the previous HNN ERP model, referred to as the *neymotin_2020_model* [29](Supp. Table 1). First, there is a notable difference in the dipole scaling factor. The *duecker_ET_model* produces the empirical ERP amplitude with a factor of 1000, implying that ∼200,000 pyramidal neurons generate the response, while the *neymotin_2020_model* required a scaling factor of 3000 [29], [31]. This difference emerges from two changes in the *duecker_ET_model* compared to the original model: 1) a reduced axial resistivity of the L5ET model from 200Ωcm to 100Ωcm accelerating the evolution of voltage along the dendritic compartments and 2) long-lasting contributions of dendritic Ca^2+^ spikes to the dipole [16]. Supp. Fig. 2f demonstrates that each Ca^2+^ spike produces a sustained downward deflection in the dipole, which is shorter and less pronounced in the previous L5 pyramidal neuron model (Supp. Fig. 3d)[48]. Due to its long duration relative to backpropagating action potentials, this downward deflection is particularly apparent in the smoothed dipole, and sums effectively across cells and time (Supp. Fig. 2f) [16], [48], [64].

Second, the mean arrival time µ and standard deviation σ of the exogenous proximal and distal drives were adjusted. In the *duecker_ET_model*, both the first proximal drive and the distal drive activate the network 5–10 ms earlier than in the original model, while the second proximal drive arrives ∼20 ms later and with greater spike-time variability (Supp. Table 1). These timing shifts follow from the membrane time constants of the network cells, which were tuned explicitly for the L2/3 pyramidal neuron and interneuron (Fig. 5, Fig. 6). While the membrane time constant was not reported for the L5ET neurons [43], we constrained it through the 3-dB cutoff of the membrane’s resistor–capacitor circuit, which the L5ET model reproduces (Fig. 3). As the membrane time constants were not explicitly considered in the development of the *neymotin_2020_model* [29], [31], we argue that the adjusted timing of the exogenous drives result from the increased biophysical realism of the model.

Additionally, all postsynaptic conductance weights are several orders of magnitude smaller, reflecting the reduced axial resistivity of the L5 pyramidal neurons, the adjusted space constants, the higher excitability of the human-tuned interneurons (Supp. Fig. 3c), and the inclusion of NMDA synapses in the input drives (Supp. Table 1).

#### New circuit elements and visualization methods further disentangle somatic, dendritic, and synaptic contributions to the ERP

Recent updates to the HNN software, combined with the biophysical updates of the *duecker_ET_model*, allow a comprehensive inspection of the cellular and synaptic mechanisms underlying each ERP component. The Python API allows users to record and inspect current dipole waveform, layer- and cell-specific firing rates, as well as somatic and dendritic membrane potentials, intracellular Ca^2+^, and synaptic currents in the network cells. These coupled dynamics contribute to the generation of the ERP waveform, as we describe below (Fig. 9).

The net current dipole shown in Fig. 9b represents the summed dipole contributions from the L2/3 and L5 pyramidal neurons, which are shown separately (dotted curves) with the layer-specific spiking activity in Fig. 9c. Due to their length, the L5ET pyramidal cells dominate the dipolar signal. The first positive peak, referred to as M25, emerges from excitatory synaptic currents from the proximal drive and backpropagating action potentials pushing current up the dendrites, as well as inhibition of the distal apical dendrite through the SOM+-like connection of the L2/3 interneurons onto the L5ET pyramidal cells pulling current towards the tuft (Fig. 9c). The negative deflection (M35) arises from peri-somatic inhibition of the L5ET population by the L5 interneurons. This spiking activity in the interneuron population is driven by excitatory input from the L2/3 pyramidal neurons, which also excite the L5ET pyramidal population. This sequence is consistent with optogenetic work showing that L2/3 pyramidal neurons facilitate and sustain stimulus-evoked responses [122]. After being briefly silenced by the L2/3 interneurons, a subset of the L2/3 pyramidal neurons fires again due to sustained excitation through the NMDA receptors (Fig. 7h). The pyramidal cells then trigger spiking in the L2/3 inhibitory population, which pulls the current up the L5ET dendrites, and generates the second positive peak (M50).

The prominent trough, M70, emerges as the distal drive pushes current toward the soma, causing a downward deflection in the dipole [31], [64]. This distal drive also triggers Ca^2+^ spikes in the L5ET population (Fig. 9e, right), causing a strong Ca^2+^ influx into the tuft of L5ET cells (Fig. 9f). To examine the separate contributions of voltage-gated Ca^2+^ channels and NMDA synapses to the dendritic Ca^2+^ concentration, we re-ran the simulation with the maximum conductance of all voltage-gated Ca^2+^ set to zero (Fig. 9g). Under this condition, the distal drive produces only a small Ca^2+^ influx in L5 pyramidal neurons at the network center, showing that the Ca^2+^ influx observed in Fig. 9f is largely driven by the voltage-gated Ca^2+^ channels (VGCC). However, note that this simulation only approximates the portion of Ca^2+^ current through the NMDA receptor, as the Ca^2+^ channels contribute to the depolarization of the membrane potential removing the Mg^2+^ block (Fig. 7d).

Lastly, the large positive components (M100, M135) correspond to upward current flow from the second proximal drive which induces a cascade of rhythmic excitatory and inhibitory cell spiking in both layers. While the somatic inhibition pulls current downward (as reflected in the fluctuations in the single trial simulations, Fig. 9b, faint orange lines), the dominant effect is current flow up the dendrite due to a combination of backpropagating APs in the L5ET population pushing current toward the apical tuft, and inhibition of the apical dendrite from the L2/3 interneurons pulling current upward (Fig. 9c, d).

#### Reproducing the alpha/beta complex of the somatosensory mu rhythm with coincident stochastic 10 Hz bursts of inputs to the proximal basal and distal apical dendrites

Following previous implementations [29], [30], [39], the model was tuned to reproduce several characteristic features of the alpha and beta components of the MEG measured somatosensory mu rhythm source-localized to SI [30]. These features were informed by the observation that the alpha and beta components of the somatosensory mu rhythm emerge as transient high-power events (or “bursts”) in the individual trials (Fig. 10); a phenomenon now observed in many brain areas [36], [37], [39], [123], [124]. While the alpha and beta simulations were tuned based on oscillations in the somatosensory cortex, due to similarities in alpha and beta oscillations in other brain areas, the defined patterns of drive have proven to be a useful starting point for studies of alpha and beta in other brain areas [36], [39].

**Fig. 10.**
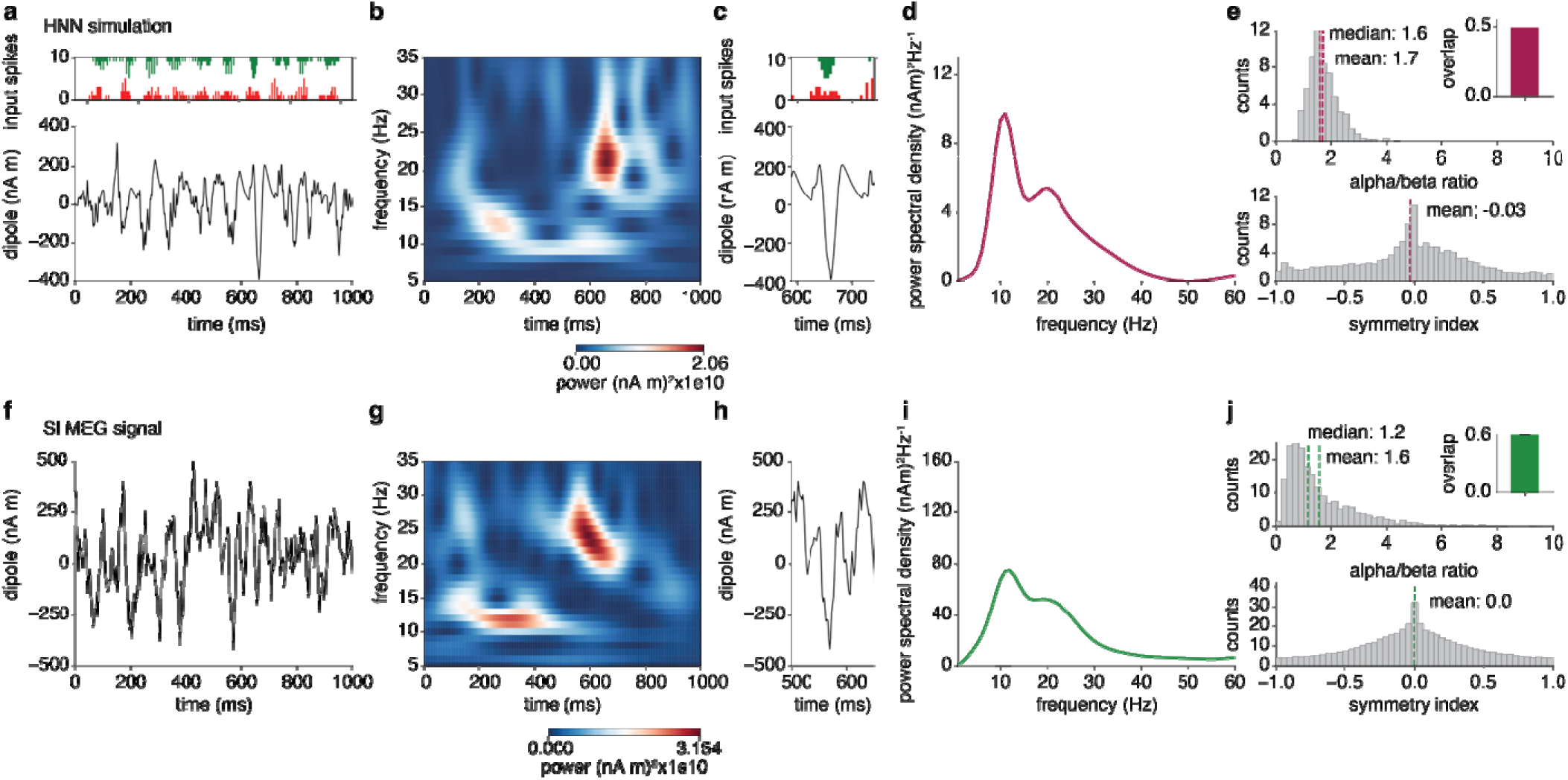
Alpha-beta complex of the SI mu-rhythm simulations using the duecker_ET_model. **a** Bursts of proximal and distal drives (top) and time course of resulting current dipole (bottom) for one example 1-second simulation. **b** Time-frequency representation (TFR) of power of **a**, showing transient alpha and beta events. **c** Zoom in on beta event at about 600 ms, demonstrating that the event is generated by a coincident proximal and distal drive, with a standard deviation of 20 and 15 ms. **d** Power spectrum averaged over all 600 simulated trials shading shows standard deviation over trials. **e** Ratio of alpha and beta power within 100 ms bins over all 600 simulated trials. (inset) Average overlap of the top 1/3 100 ms bin with highest alpha and the top 1/3 bins with the highest beta power. (bottom) Symmetry index of peaks and troughs, following approximately a normal distribution. **f-j** MEG data. **f** Single trial pre-stimulus MEG data, one exemplary subject and trial. **g** TFR of MEG trial in **f**, showing events in the alpha (8-12 Hz) and beta band (15 - 25 Hz). **h** Zoom in on beta event between 500 and 600 ms, showing a similar shape to the one predicted by the simulation. **i** Power spectrum averaged over all trials and participants shading shows standard deviation over participants. **j** Ratio of alpha and beta power, average high alpha/beta overlap, and symmetry index over all trials and participants, exhibiting similar distributions to the simulated data.

As in prior studies, the mu rhythm was simulated by driving the network with approximately coincident 10 Hz bursts of input through proximal and distal projection pathways (Fig. 10). In contrast to the evoked drives, the bursty drive was simulated with 10 drive cells, each connecting to all network cells [29]. Fig. 10a shows the current dipole for one example trial. The histograms in the top panel show the spike times of the proximal and distal drive cells for this trial. These spike times were simulated as periodic population bursts, consisting of two spikes with an inter-spike interval (ISI) of 10 ms. The time of the first spike in the burst was drawn from a Gaussian distribution with mean spike time repeated approximately every 100ms, and standard deviation σ=20 for the proximal drive and σ=15 for the distal drive (Supp. Table 2) [39]. Additionally, on each cycle of the drive, the mean spike times of the distal drive were delayed by 5ms relative to the proximal drive, consistent with previous implementations (Supp. Table 2) [29], [30]. The postsynaptic effects were mediated through AMPA synapses only, whose synaptic weights were selected so that the membrane potential fluctuations in the postsynaptic cells remained subthreshold, modulating membrane potentials without triggering spiking in the network. As in previous implementations, the AMPA conductance of the distal drive was set to be four times stronger than the proximal [30], [39].

The nearly synchronous rhythmic inputs drive currents up and down the pyramidal neuron dendrites to create a primary current dipole signal that closely matches the quantified features of the empirical data, including the event-like nature of alpha and beta activity, characteristic waveform shapes, and power spectral density profiles (compare Fig. 10a-d to f-I, 600 simulated trials). Due to the stochasticity of the input burst spike times, the proximal and distal drives occasionally arrive approximately coincidentally at the basal and apical dendrites. When the coincident distal drive standard deviation is small, the drive cells arrive within a short time window, causing strong postsynaptic conductance to break the upward current flow, and generating a brief, high-amplitude event at about 25 Hz, referred to as a beta event (Fig. 10c)[39]. Alpha events occur on cycles when the proximal and distal drive are further apart. The simulation shows close agreement with several spectro-temporal properties of the empirical data including (1) the ratio of alpha and beta power within epochs of 100 ms (Fig. 10e, j, top), (2) the overlap between high powered alpha and beta events (Fig. 10e, j, top, inset), and (3) the symmetry of the peaks and troughs in the time series (Fig. 10e, j, bottom), quantified as in Jones et al. 2009 [30].

The parameters of the input drives are outlined in Supp. Table 2, juxtaposed with the values chosen for the *neymotin_2020_model* [29], with several notable differences. Again, there is a sizable difference between the dipole scaling factor, with a factor of 20,000 applied to the *duecker_ET_model* output, estimating 4,000,000 pyramidal neurons that contributed to the recorded data, as opposed to the factor of 300,000 applied to the *neymotin_2020_model*, estimating contributions from 60,000,000 pyramidal neurons. As the incoming drives did not generate spiking in the network, this difference can be attributed to the reduced axial resistivity of 100MΩ for the *duecker_ET_model*, compared to 200MΩ in the old model.

#### Reproducing continuous and transient event-like gamma oscillations

Following previous implementations, the model was tuned to reproduce a well-known canonical mechanism of a 30-60Hz gamma rhythm known as Pyramidal-Interneuron Network Gamma (PING) [29], [62]. PING rhythms are generated by fast synaptic interactions of excitatory and inhibitory neurons, where the ∼25ms period is set by the time-constant of decay of strong PV+ mediated GABAA inhibition [125]. In this example, the network activation parameters were tuned to reproduce both sustained and transient gamma oscillations, as in [29]. For illustrative purposes, the interlaminar excitatory connections were all set to zero, and pyramidal-to-pyramidal connections were removed to examine PING mechanisms in each layer separately, focusing in the timing of the GABAA-mediated inhibition, as described previously [29], [62].

To reproduce sustained gamma oscillations, each layer of the network was activated with a stochastic excitatory proximal drive. The drive was simulated with a continuous train of action potentials following a high-frequency Poisson process (base frequency 200Hz, AMPA/NMDA postsynaptic conductances: O_L23_ = 0.0045 S/cm^2^, O_L5ET_ = 0.0055 S/cm^2^) contacting the network neurons through the proximal drive pathway with synaptic delays of 0.1ms and 1ms for the L2/3 and L5ET pyramidal neurons (Fig. 11a, b). The local network connections were tuned to have strong peri-somatic PV+ inhibition (Supp. Table 3) so that the resulting dipole oscillates in the 40-60Hz gamma band (Fig. 11b, d). Layer-specific spiking shows coordinated firing in subsets of L2/3 and L5ET pyramidal neurons that drives highly synchronous interneuron firing in turn inducing a ∼25ms pause in network activity, characteristic of the PING rhythm (Fig. 11c, “weak” PING [62]). The power spectral density shows a narrow peak at 51 Hz in the aggregate (e.g. sum of L2/3 and L5) and L5 dipoles, with a slightly higher peak frequency in L2/3 (Fig. 11e). Inspection of the membrane potentials in an exemplary L2/3 pyramidal cell shows high-frequency somatic spiking, whereas the exemplary L5ET cell fires at lower rates (Fig. 11f, g). Neither cell type produces Ca2+ spikes at the tuft, indicating that the Poisson drive did not trigger L5ET bursting.

**Fig. 11.**
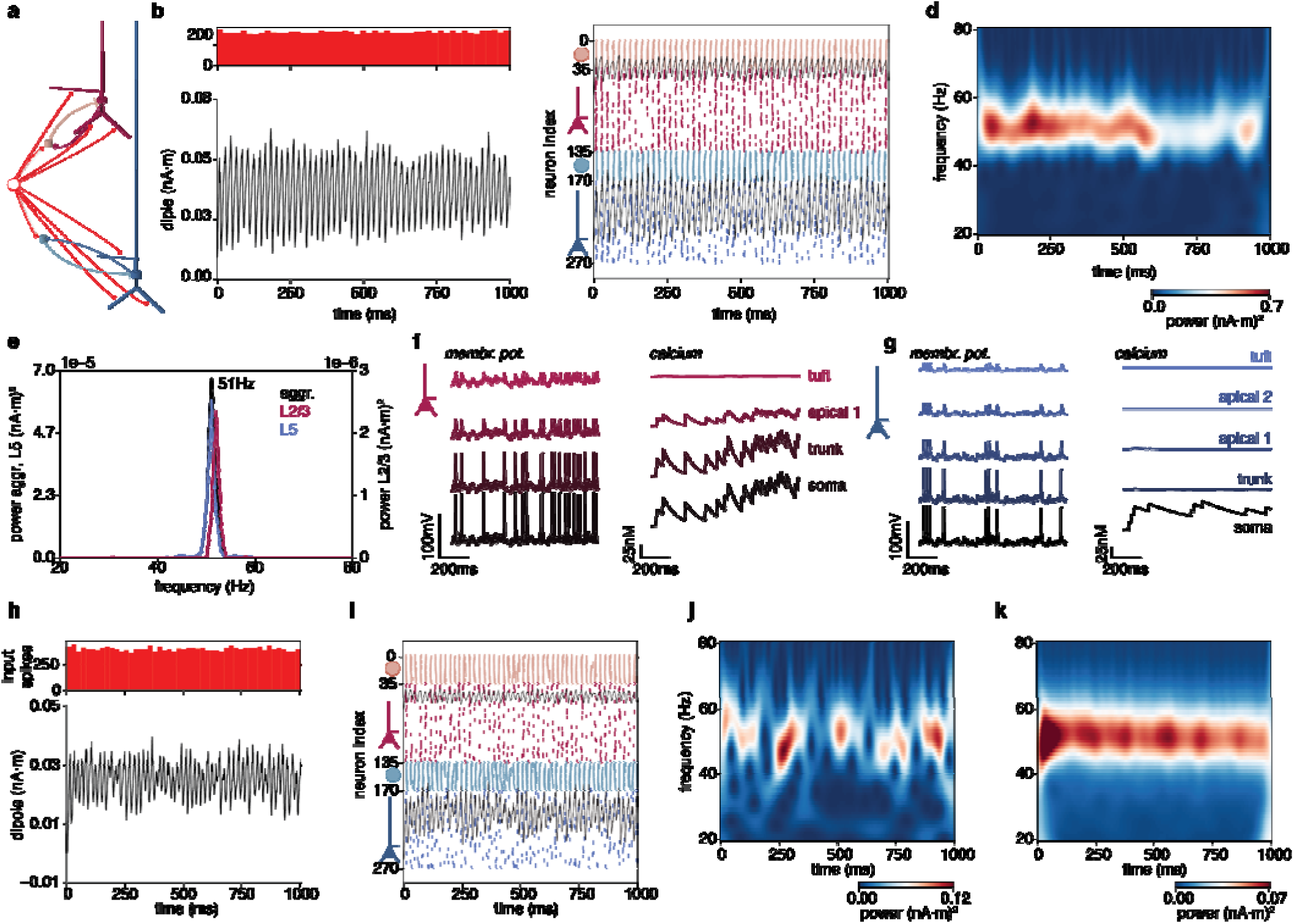
Pyramidal-Interneuron-Network-Gamma (PING) simulations using the duecker_ET model. **a** As in prior studies, for illustrative purposes, all excitatory to excitatory and interlaminar connections were set to zero. The network is activated by a continuous proximal poisson drive with a high base frequency. **b - g** Continuous PING. b (top) Spike times of the high-frequency Poisson dr ve to the network, resulting in approximately continuous oscillations in the dipole (bottom). **c** Layer-specific spike rasters and dipoles (black line plots) show rhythmic firing of the interneuron populations that is preceded by organized firing of the pyramidal neurons **d** Time-frequency of power showing continuous oscillations with a peak frequency of about 51 Hz. **e** Power spectral density the aggregate and L5 dipole (left y axis) and L2/3 dipole, showing that the gamma in L2/3 is a few Hz faster than L5. **f** Membrane potential and Ca^2+^ for the soma and apical compartments of an exemplary L2/3 pyramidal neuron, revealing spiking and Ca^2+^ activity in the compartments proximal to the soma but not in the tuft. **g** Same as in **f**, for one exemplary L5 compartment. The L5 cell shows spiking, but no bursting activity and no dendritic Ca^2+^ spikes. **h - k** Transient PING (i.e., gamma events). **h** (top) Poisson inputs are lower in frequency than for the continuous gamma simulation in **b**. (bottom) the dipole shows a DC shift with intermittent periodic activity. **i** Layer specific dipole and spiking activity shows that interneurons again fire at a gamma frequency, the firing rate of the pyramidal neurons is lower than for the continuous gamma. **j** TFR of power for a single trial, showing the event-like nature of the gamma activity. **k** Average TFR over 100 trials, showing a continuous gamma rhythm.

Similarly to the somatosensory mu rhythm, gamma oscillations in human E-/MEG have been shown to occur in transient events [29], [126]. Reproducing this behavior required the reduction of the inhibitory-to-excitatory connection strength by a factor of three and applying a Poisson drive with a slower base frequency of 100Hz and reduced synaptic weights (Supp. Table 3). The single trial simulation shows a positive DC shift at drive onset followed by an approximately rhythmic signal of varying amplitude (Fig. 11h). Interneuron firing remains rhythmic, but pyramidal neuron firing is notably sparser (Fig. 11i) than in the continuous gamma condition (Fig. 11c). The corresponding Time Frequency Representation (TFR) of power shows transient gamma-band events (Fig. 11j) that appear continuous when averaged over 100 trials (Fig. 11k).

Comparison between the parameters used here and in the *neymotin_2020_model* [29] shows that the inhibitory-to-pyramidal and pyramidal-to-inhibitory weights are of comparable magnitude across the two models. The inhibitory-to-inhibitory connections in the *duecker_ET_model* are an order of magnitude weaker, reflecting the higher excitability of the interneurons (Supp. Fig. 3c). Sustained oscillations required a faster Poisson base rate in the *duecker_ET_model* (200 Hz). This is in part explained by the use of both AMPA and NMDA synapses in the *duecker_ET_model*, while only AMPA connections were used in the *neymotin_2020_model* (140 Hz) [29].

Adjustments made to simulate the transient gamma events differed strongly between models. In the *duecker_ET_model*, the local inhibitory-to-pyramidal weights were reduced and the network was driven with a 100Hz proximal Poisson drive. In the *neymotin_2020_model* [29], the local connections remained fixed to the same value as in all other simulations, and only the proximal Poisson drive parameters were adjusted to include NMDA synapses and a slow 4–5 Hz Poisson drive. These differences in the required drive parameters likely reflect the distinct bursting dynamics of the L5 pyramidal neurons. In the *neymotin_2020_model*, L5 pyramidal neurons burst even in response to brief inputs. The L5ET cells in the *duecker_ET_model* require longer synaptic inputs to generate a burst (Supp. Fig. 3c). The AMPA and NMDA weights of the Poisson drive were of similar magnitude in both models.

## Discussion

We present a new neocortical column model for the HNN software constrained by human cell physiology. The updated model is distributed as *duecker_ET_model*, alongside major software updates that facilitate multiscale interpretations of human E-/MEG signals. Collectively, these updates are referred to as the next generation HNN. The model retains the reduced morphological detail and laminar architecture of the original HNN model [29], [30], [31], with a level of model complexity that is tractable to study the microcircuit generators of E-/MEG signals. The active ion channel kinetics and maximum channel conductances of the cell models were tuned to reproduce somatic dynamics observed in human neurons in vitro [42], [43], [88], [92] as well as dendritic Ca^2+^ spikes [69], [70]. The *duecker_ET_model* also features biologically realistic synaptic kinetics, and the synaptic weights between cells were tuned to reproduce PSPs from human tissue recordings [41], [44], [45], [46], [47], [107]. As a starting point for future studies, we provide pre-tuned thalamocortical and corticocortical activation patterns (drives) to generate somatosensory ERPs (Fig. 9), alpha-beta (Fig. 10) and gamma oscillations (Fig. 11), alongside source code to reproduce all figures in this work.

The comparison between parameters for the *duecker_ET_model* and the original model [29], across ERPs and alpha, beta, and gamma oscillations shows that conserved network dynamics can emerge from distinct parameter regimes in the two models. In each simulation with the *duecker_ET_model*, only parameters regulating the effective strength of the exogenous proximal and distal excitatory drive were tuned to reproduce similar network behaviors when simulating the ERP and alpha-beta rhythm. This underscores how the biophysical detail of the *duecker_ET_model* reshapes the response of the network to driving input but does not change the ability of the network to model activity underlying fundamental E-/MEG signals.

### Incorporating realistic dendritic calcium dynamics

A key goal of this work was to simulate dendritic Ca^2+^ spikes in the pyramidal neurons. These dynamics are interesting in the context of E-/MEG as they occur in the apical dendrites generating the primary currents, volume currents, and magnetic fields [14], [15], [16], [17], [18], [19], [20] (Box 1). Moreover, they have been repeatedly shown to equip pyramidal neurons with rich computational abilities, such as amplifying synaptic inputs in cortex [69], [127], [128] and in hippocampus [129], [130]. Importantly, dendritic Ca^2+^ spikes are known to influence the spiking output of the neuron [69], [131] and have been linked to synaptic plasticity [82], [132]. As such, they shape not just the activity of individual neurons, but the dynamics of the circuits giving rise to meso- and macroscale signals such as LFP and E-/MEG [17]. Additionally, dendritic Ca2+ spikes in L5 pyramidal neurons have been detected in extracellular fields and epidural recordings [76]. With the *duecker_ET_model,* we provide the tools for studying their involvement in generating macroscale E-/MEG signals.

#### Comparison to other modeling tools

Several detailed biophysical models have been developed to study microcircuit dynamics alongside mesoscale field dynamics [14], [18], [52], [133], [134], [135], [136] and macroscale E-/MEG signals e.g. [137], [138], [139], [140], [141], [142], [143]. Many of these models are morphologically complex and computationally demanding, and working with them often requires specialized training in computational neural modeling as well as high-performance computing capabilities. To make biophysical detailed modeling accessible to researchers from different backgrounds, several software tools and tutorials have been developed [141], [144], [145], [146]. HNN adds to this effort with a GUI and a Python API that provide access to models and activation workflows specifically designed to study the neural generators of E-/MEG signals. The reduced morphology of the cells in these models keeps them computationally efficient, allowing simulations on a personalized computer, and their outputs interpretable. Importantly, the simulated primary current dipoles are simulated in the same units as source-localized dipole time courses provided by the user, allowing direct comparison. As such, HNN provides a powerful framework to develop and test hypotheses on how cell and circuit elements contributed to recorded data. Recording and visualization tools give access to the underlying firing rates, membrane potentials, ion concentrations, synaptic currents, and extracellular fields. Unlike tools focused on tuning individual cells, HNN assumes that users will generally work with a provided pre-tuned neocortical model together with pre-tuned driving inputs. HNN’s optimization tools are therefore designed to tune the exogenous drive patterns rather than the cells [29], [90], [147]. This enables researchers to approach biophysical modeling of a wide range of neuroscientific questions without building a new model.

#### Limitations and next steps

The rapidly growing body of literature characterizing the dynamics of single neurons and microcircuits facilitates the development of highly detailed biophysical models. Detailed models are valuable for many questions, such as how cell morphology shapes extracellular fields [14], [16], [17], [19], [20], [134], [140], [148]. However, each added detail increases the number of interacting dynamics, and the behavior of highly complex models can quickly become as difficult to interpret as the neural process they were built to explain [5], [149]. HNN models therefore aim for a level of abstraction that reproduces the phenomena of interest while still providing insight into the underlying mechanisms. To study the generators of the macroscale E-/MEG signals, we therefore made the following simplifying assumptions.

All cells are modeled with reduced morphological detail. This reduces computational demands and keeps model outputs interpretable while still supporting simulations of active dendritic dynamics. However, cell morphology shapes the distribution of transmembrane currents, which in turn shapes the extracellular fields [150], [151] and the E-/MEG signals [140]. While HNN simulates the waveform shape and spectral properties of E-/MEG signal (in contrast to other reduced morphologies [140]), reliably reproducing the LFP and CSD obtained using laminar recordings in rodents and non-human primates is challenging, and was therefore not explored in this publication. This limitation is not unique to HNN, as highly detailed biophysical models have also required dedicated extensions to reproduce the CSD [52], [113]. This work has shown that LFP and CSD do not just depend on the morphology, but also the patterns of synaptic input [52]. The reduced architecture of HNN models is nonetheless useful for predicting the laminar generators of the E-/MEG signatures of interest [32], [39].

The presented model simulates L5ET but not L5IT cells. This abstraction is motivated by a wide interest in dendritic Ca^2+^ dynamics observed in rodent ET cells. Moreover, L5ET neurons have longer apical dendrites than L5IT cells [42]. As the dipole moment of an individual cell depends on the length of the apical dendrite [20] (equation (5)), the individual contribution of an L5ET cell to the E-/MEG signal is likely larger than that of an IT cell. In vivo firing rates of putative ET cells in rodents cells have also been reported to be higher than those of putative IT cells [118], [152], [153], [154]. However, reports of cell-type-resolved activity are currently not available in primates. On the other hand, L5IT cells have been repeatedly reported to be more abundant than ET cells, particularly in humans [42], [43], [75], [155]. As such, their collective contributions to the dipolar currents may be substantial if enough cells are synchronously active. Adding L5IT cells is therefore a natural extension of the model.

The L5ET model shows stronger dendro-somatic signaling than reported for human L5 pyramidal neurons [84], [85]. This attenuation has been argued to result in weaker dendritic spikes in human neurons with little effect on the somatic output [85]. However, transcriptomically labelled human L5 ET cells have been shown to generate prolonged dendritic plateau potentials upon local current injection into the dendrite [43]. To our knowledge, there has so far not been a systematic test of whether high-frequency action potentials or coincident feedforward and feedback inputs elicit Ca^2+^ spikes in transcriptomically labeled human ET cells in a similar way as in rodents [69], [70]. Therefore, we accepted the weaker attenuation to be able to generate dendritic Ca^2+^ spikes in the L5ET model. This decision was motivated by their reported importance in sensory perception [78], [79], [80] and learning [82]. As HNN aims to enable cross-species translation, including Ca^2+^ spikes allows researchers to develop hypotheses about how they could shape network dynamics and E-/MEG signals.

L2 and L3 pyramidal neurons are represented by a single L2/3 cell type in the model. However, recordings from human tissue have shown that the transcriptomic, morphological and electrical properties of L2/3 pyramidal neurons vary with cortical depth [42], [88], [91]. These differences have been attributed, at least in part, to depth-dependent distributions of I_h_ channels [91]. At the same time, however, L2 and L3 pyramidal neurons overlap considerably in morpho-electric feature space (Fig. 1 in [42]). Moreover, it is currently unknown how the identified differences between L2 and L3 pyramidal cells affect firing rates in vivo. We therefore kept the joint L2/3 pyramidal cell type of previous HNN models.

HNN models do not simulate L4 cells but represent thalamic input reaching L2/3 and L5 via L4 with the proximal drive (Box 1). This abstraction was chosen because the model was originally developed to simulate the tactile evoked response in SI [29], [31]. L4 in SI is dominated by spiny stellate cells [156], [157], whose dendrites extend in many directions, causing their transmembrane currents to largely cancel and contribute comparably little to the dipole [17], [18]. We therefore hypothesized that the influence of L4 on the E-/MEG signal is largely indirect, through synaptic input to L2/3 and L5 pyramidal neurons, which is modeled with the proximal drive. Other cortical regions contain L4 IT pyramidal cells, e.g. rodent V1 [156], A1 [154], and M1 which was long considered agranular layer due to the lack of stellate cells [158], cat A1 [159], and several areas in human cortex, including M1, MTG [42], [155] A1, S1, and V1 [75]. However, the direct contributions of these cells to the E-/MEG signal are unknown. Because currents generated in L4 are an important landmark for identifying layers in laminar recordings [55], [56], [57], [160], future HNN models focused on extracellular fields may explicitly model L4 cells rather than abstracting them with a drive.

Finally, all inhibitory cells are modeled with the same generic interneuron model. The different types are represented by the kinetics and locations of their synapses on the postsynaptic neuron. This preserves the architecture and morphology of previous HNN models. While the presented model was not built to explicitly test hypotheses about the involvement of different inhibitory interneuron types to shape the E-/MEG signal, expanding to more diverse interneuron types is a natural next step [143]. Model validation is however complicated by the limited availability of cell-type-resolved in vivo firing rates.

## Conclusion

Our updated neocortical model, termed *duecker_ET_model*, simulates a wide array of physiologically plausible somatic, dendritic and synaptic dynamics, as well as macroscale E-/MEG signatures alongside biophysically realistic firing rates. The model is shared within a major update of the HNN software suite, alongside updated visualization tools that facilitate the multiscale interpretation of E-/MEG signatures, we refer to as the Next Generation HNN. This manuscript and the associated source code provide interested users with a starting point for interpreting their E-/MEG signatures of interest in terms of the underlying cell and circuit generators.

## Methods

### Computational Neural Model

#### Cells

##### Data used to constrain cell tuning

The L5 pyramidal model was tuned to reproduce 12 measures from in vitro recordings at the soma that have been shown to distinguish L5ET neurons from L5 intra-encephalic projecting (IT) cells [43] (https://brain-map.org/our-research/cell-types-taxonomies/multimodal-characterization-of-individual-neurons#human). The L2/3 and interneuron model cells were each tuned to 9 electrophysiological measures from human layer 2 and 3 spiny and aspiny neurons [40], [42], [88], [92].

##### Identifying tuning targets for all cell models

We identified the resting membrane potential as the voltage at the soma after a 100ms simulation, with all cell compartments initialized at -65mV. Action potentials at the soma were triggered using a single 5ms pulse, and the action potential threshold was identified as the membrane potential at which dV/dt exceeded 20mV/s [43]. The lowest voltage following the peak of the action potential, normalized to the threshold, was identified as the fast afterhyperpolarization. For the L2/3 pyramidal neurons and interneurons, we estimated the upstroke/downstroke ratio by dividing the slope of the increase in membrane potential between action potential threshold and action potential peak by the slope of the decrease between the peak and fast afterhyperpolarization.

The sag ratio and rebound slopes were calculated based on hyperpolarizing current injections into the soma using amplitudes from the Allen Cell Types database [40]. For the L5ET cell, the sag ratio was computed as R_max / R_ss, where R_max and R_ss are linear fits of input current against peak and steady-state voltage [43]. The sag ratio of the L2/3 pyramidal neuron and interneuron was defined as the ratio of the difference between sag amplitude and steady state, normalized by sag amplitude (following https://github.com/AllenInstitute/ipfx). The rebound slope was calculated as a linear fit between rebound peak and steady state [43]. For the L2/3 pyramidal neurons and interneurons, we determined the membrane time constant from these experiments based on a single exponential fit.

As information on the membrane time constant of the L5ET cell was not available from the openly shared data, we quantified the temporal properties of the membrane using a subthreshold chirp stimulus at frequencies between 0.2 and 40 Hz. The impedance spectrum was calculated as the ratio of the Fourier transforms of the voltage response and the chirp stimulus (Fig. 2b, center left and center). The resonance frequency was then identified as the frequency with the highest peak in the impedance spectrum, and the 3db cutoff as the frequency at which the impedance profile attenuated to sqrt(0.5*Zmax) [43].

We characterized the firing pattern of each cell using a simulated 2s continuous current injection. For the L2/3 pyramidal neuron and interneuron we extracted the inter-spike interval (ISI) between the first two spikes (first ISI) and the average ISI (mean) during the stimulus (Fig. 4b, Fig. 6a). The burst ISI for the L5ET cells was calculated as the mean spike interval within the first 80ms of the 2s stimulus (Fig. 2b).

##### L5ET passive properties

We set the membrane capacitance to the biophysically realistic value of cm=1µF/cm2 [85], [161]. While previous work suggests that the membrane capacitance in biophysical models should be increased to 2µF/cm2 to account for dendritic spines [77] we found that cm = 1µF/cm2 was required to simulate Ca^2+^ spikes and largely reproduced the electrotonic properties of human L5 neurons [85](Supp Fig. 2, but see Discussion). The axial resistance (Ra) of the model was set to Ra = 100Ωcm matching biophysically identified values [77], [162]. Original reports recommended increasing Ra to 200Ωcm for reduced morphology cells, however, these simulations only considered passive dendritic compartments [53], [163]. For this model, we found that a lower Ra was needed to achieve somato-dendritic coupling that generates dendritic Ca^2+^ spikes (Fig. 5).

##### L5ET ion channels

Active ion channels were adapted from a model of a morphologically detailed rat L5b pyramidal neuron [77], which were assembled from several primary sources. As the default temperature for simulations in HNN is 37°C, a Q10 of 3.7910 was applied to channels originally characterized at lower temperatures: the fast inactivating Na^+^ channel (Nat)[164], persistent Na^+^ channel (Nap) [165], slow inactivating K^+^ current (Kp) [166], fast inactivating K^+^ current (Kt) [166], muscarinic K^+^ current (Im) [167], and low-voltage activated Ca2+ current [168], [169]. Four channels did not require a temperature correction: The high-voltage activated Ca2+ channel [170](Ca2+_HVA) was characterized at 36°C and the non-specific cation current [171](I_h_) at 35°C. The fast-non activating K+ channel [172](Kv3.1) governs the fast afterhyperpolarization which we explicitly tuned for each cell (see Fig. 33 and Supp. Fig. 3). The time constant of the small conductance Ca^2+^-activated K^+^ (SK-E2) current was treated as instantaneous [77], [173].

Evolutionary algorithms are standard for tuning active maximum ion channel conductance, but scale poorly with objective count [174], [175], [176], [177]. Given our 12 somatic objectives plus dendritic constraints, we tuned manually, guided by the literature. As a starting point we halved all active maximum conductance values used in [77], to reflect the reduced channel densities and increased compartmentalization reported for human L5 pyramidal neurons [84], [85]. We then adjusted all values until all somatic measures fell within the biophysical range of human neurons [43](Fig. 3b).

Three ion channels were distributed non-uniformly along the apical dendrite (Supp. Fig. 1). I_h_ increases exponentially along the apical dendrite following an experimentally identified distribution [77], [171]. Ca^2+^ HVA and LVA increase linearly to permit Ca^2+^ accumulation in the apical dendrite [72]. In line with the L5b model [77], we added a Ca^2+^ “hotzone” to the bottom of the apical tuft compartment where the HVA and LVA densities were increased 4.5- and 2.25-fold. This hotzone was critical for generating dendritic Ca^2+^ spikes [77], [83], [178](Supp. Fig. 1b).

The original L5b model [77] did not include active ion channels in the basal dendrites. Because the L5 pyramidal neurons in HNN receive thalamocortical and neocortical inputs at their basal compartments, we added Nat, Kv3.1, I_h_, and a passive leak, with maximal conductance tuned to attenuate backpropagating action potentials to ∼65–70 mV at the basal tip, consistent with in vitro recordings from rodents [179], [180](Supp. Fig. 1c). In the interest of reducing the number of free parameters in the tuning process and computational demand of the model, and due to the lack of guidance on basal dendrite tuning in the literature, we refrained from adding other ion channels to the basal dendrites.

##### L2/3 pyramidal passive properties

The axial resistivity was set to Ra = 200Ωcm to reproduce the strong attenuation of depolarizing and hyperpolarizing currents between dendrite and soma observed in the literature [89], [90](Supp Fig. 3d,e). The membrane capacitance was raised to cm=1.5µF/cm2 to match the membrane time constant of human L2/3 pyramidal neurons (Supp Fig. 2b).

##### L2/3 pyramidal ion channels

Following a previous implementation of a rodent L2/3 [87] we included all ion channels from the L5b cell [77], except for Kt and Kp (Table 2). To widen the action potential compared to the L5ET cell, we shifted the kinetics of the Nat channel to 32°C. Starting from the conductance values used in the previous publications [77], [87], we manually tuned the soma until all 9 electrophysiological measures fell within the human data range [42], [88].

A previous model of the human L2/3 pyramidal neuron simulated the dendritic Ca2+ spike but did not include the Na+ and K+ channels underlying somatic action potentials [89]. Other models of L2/3 pyramidal neurons were not explicitly tuned to generate dendritic Ca2+ spikes [143]. We therefore drew on our experience with the L5ET cell and distributed I_h_, Ca^2+^ HVA and LVA non-uniformly between soma and tuft (Supp. Fig. 1f). As Ca^2+^-activated K^+^ channels have also been suggested to be non-uniformly distributed in L5 pyramidal neurons [181], [182], [183], we chose a similar profile for the SK-E2 channel which shapes spike duration (Supp Fig. 1f). While Ca^2+^-activated K^+^ channels have been proposed to vary along L5 pyramidal dendrites [181], [182], [183], the specific distribution of the SK-E2 channel in L2/3 pyramidal neurons has not been characterized experimentally. The tuned model predicts that the Ca^2+^ and K^+^ channel distributions producing these fast dendritic spikes follow the non-uniform patterns shown in Supp. Fig. 1f. To enable the faster dendritic spikes observed in L2/3 [89], we reduced the Ca^2+^ decay time constant and increased the fraction of free Ca^2+^ relative to the L5ET model (Tables 1, 2).

##### Interneuron active and passive properties

Compared to the original HNN model we reduced the interneuron size to 400 µm² and set Ra = 200Ωcm and cm = 1µF/cm². The electric properties of these point neurons were tuned to in vitro recordings from human aspiny cells [40]. Channel equations were adapted from a previous model of fast-spiking interneurons [93], [94], including a Na+ current (nas), delayed rectifier K+ current (kdr), D-type K+ current (kd), and passive leak (pas; Table 3). We added an I_h_ current to reproduce the sag observed in human PV+ and SOM+ interneurons [92], [184]. Although the interneuron model lacks active Ca^2+^ channels, it receives NMDA-mediated Ca^2+^ influx through excitatory synapses (see below). We therefore tracked intracellular Ca^2+^ with decay parameters τ_decay = 50 ms and γ = 0.0005096 (Table 3).

#### Synapses and Network Connectivity

Each layer in the model consists of 100 pyramidal neurons and 35 interneurons, aligning with estimated density of interneurons of about 25-40% relative to pyramidal neurons in rodents [113], [185], [186], [187] and non-human primates [188]. However, current estimates in humans suggest a higher density of inhibitory neurons [75], [189].

##### Synaptic dynamics

AMPA, fast and slow GABAA receptors were simulated using a double exponential formalism (AMPA =0.5,5 ms, rise, decay; fast GABAA 0.5, 5ms; slow GABAA 1.5: 20ms)[99]. For NMDA synapses, we adopted a biophysically explicit model [95], [98], which includes an Mg^2+^ block and an explicit Ca^2+^ current. The biophysically detailed GABAB synapse models a second-order G-protein transduction and K^+^ channel opening [101], and can be optionally included.

##### Intra- and inter-laminar connectivity

The strength of each pairwise connection within and across layers decreases with distance along the horizontal plane [41], whereby the synaptic conductance decreases and synaptic delays increase following a Gaussian distribution. The synaptic conductance *s* and synaptic delay *d* for connections between two cells *i* and *j* is calculated as:

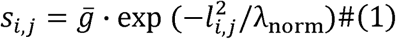

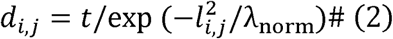

with □ being the maximum synaptic conductance, and *t* the minimum synaptic delay defined for connections between the two cell types, *l_i,j_* being the planar distance between the cell and λ*norm =* λ *· inplane_distance* being the space constant controlling the rate at which synaptic strength and delays change with distance. This abstraction preserves the functional effect of distance-dependent connectivity, while ensuring that connections are sufficiently strong to modulate spiking activity in the network.

After identifying the distance-dependent connectivity profile, we tuned the maximum synaptic conductance between different cell types based on electrophysiological recordings from human tissue (Fig. 8b). Reproducing experimental protocols, we iteratively injected a 5 ms pulse into the soma of one cell at a time and recorded the membrane potential at the soma of all postsynaptic neurons (Fig. 8e). We ensured that the strength of the connectivity would result in postsynaptic potentials within a plausible range of those identified in vitro. Furthermore, we verified at the circuit level that excitatory connections were strong enough to propagate activity between neighboring cells, while inhibitory connections were sufficient to prevent runaway excitation. For intralayer connections from the pyramidal neurons to interneurons, we set the conductance of the NMDA receptors to 18% (L2) and 20% (L5) of the AMPA conductance, to account for the lower density of NMDA receptors compared to AMPA reported for PV+ interneurons [190], [191].

#### Network simulations

##### Exogenous inputs to the neocortical column

All exogenous inputs to the HNN model are excitatory [31], [192], [193], [194] and reflect thalamocortical and neocortical inputs to the network (Box1). The HNN software provides three options for defining the spike times of these drives. *Evoked drives*, recommended for simulations of event-related responses (ERPs), draw spike times from a Gaussian distribution with a user-defined mean and standard deviation. *Bursty drives* stochastically generate rhythmic spike trains at a user-defined frequency. *Poisson drives* sample input spike times from a Poisson distribution with a user-defined base frequency [28], [29]. By default, the evoked drive uses a space constant of λ = 3, strongly driving cells at the center of the network, while the bursty and Poisson drive use a space constant of λ = 100, driving all cells in the network equally. These settings were used, respectively, for the ERP, alpha-beta, and gamma simulations.

##### Calculation of the primary current dipole

The equivalent current dipole is calculated from the intracellular axial currents flowing along the apical dendrites of pyramidal neurons [14], [16], [31]. For each segment *s* in each apical compartment of each pyramidal cell, the dipole moment is computed as:

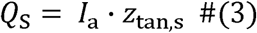

with the intracellular axial current *I*_a_ given by

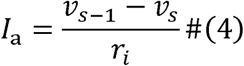

With v_s_ being the membrane potential of segment *s*, *v_s-1_* the membrane potential of the adjacent segment, and *r_i_* the axial resistance between them. *z_tan_*is calculated as

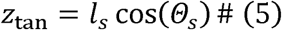

where *l*_s_ is the segment length and θ_s_ is the angle between the segment s and the apical trunk.

The dipole moment of a single pyramidal cell is the sum of its segment dipoles, and the total dipole simulated by the model is the sum over all pyramidal neurons.

##### Baseline correction of the dipole

The non-uniform density of I_h_ channels along the pyramidal neurons produces a gradient in the resting membrane potential across the apical compartments, as observed in vitro [171]. This yields a non-zero dipole even in the absence of thalamocortical input. To correct for it, we identified a single exponential fit to each layer-specific dipole for a simulation without any inputs. This fit is automatically subtracted from the simulation when using the *duecker_ET_model* in the Python API.

##### Optimization to the MEG suprathreshold tactile evoked response

Hand-tuned parameters served as initial conditions for optimization using Covariance Matrix Adaptation Evolution Strategy (CMA-ES)[195], an evolutionary algorithm recently added to the HNN-core API [147]. CMA-ES iteratively samples parameter sets from a Gaussian distribution, evaluates a user-defined loss, and updates the distribution’s mean, covariance, and step size toward successful solutions [195]. Here, we used a population size of 50 parameter sets that were evaluated over 50 iterations.

In HNN, the goodness of fit is typically quantified using the root mean squared error (RMSE) or the Pearson correlation coefficient (r) [147]. As RMSE quantifies the mean distance between simulated and experimental dipole, dipoles with complex waveforms including closely spaced smaller peaks are sometimes not well reproduced.Correlation, on the other hand, captures waveform shape but is insensitive to amplitude. To optimize for both waveform shape and amplitude, we developed a new measure:

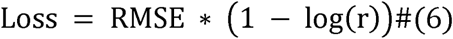

with r clipped to [1 e - 1 0, 1 ]. When the predicted and target waveforms have the same shape (r = 1), the loss reduces to the RMSE. For solutions that lower the average distance to the MEG dipole but differ in shape, the penalty increases as the waveform diverges from the target.

Further penalties were added for parameter combinations that did not lead to spiking in the L2/3 and the L5ET pyramidal neurons in response to the first proximal drive, as well as simulations with more spiking in the L5ET compared to the L2/3 pyramidal population in the first 40ms of the simulation.

The alpha-beta and gamma simulations were hand-tuned to reproduce the properties of interest.

### MEG data analysis

#### ERP

The suprathreshold ERP, used as the target for the ERP simulations, was calculated as the average tactile evoked response over seven participants [31] and can be downloaded from the hnn-data repository (https://github.com/jonescompneurolab/hnn-data).

#### Alpha-beta rhythm

The alpha-beta components of the somatosensory mu rhythm were compared to several measures obtained from pre-stimulus MEG data (https://github.com/jonescompneurolab/SpectralEvents) as described in [30]. The time frequency representation of power (TFR) was calculated using a normalized Morlet wavelet of the form

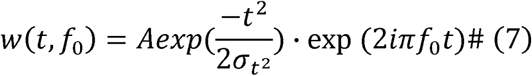

for each frequency of interest f_0_ with 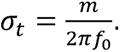

The normalization factor 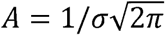 ensures that the sum of the magnitude of the wavelet coefficients over all frequencies is one and emphasizes higher frequencies to more effectively visualize the time course of the 15-29 Hz beta rhythm [30].

We quantified the distribution of alpha and beta components of the mu rhythm by dividing the trial-averaged TFRs into 100 ms windows and averaging power within the alpha (7–14 Hz) and beta (15–29 Hz) bands in each window. To assess whether high alpha and high beta power tended to co-occur, we identified the windows with highest third of alpha and the highest third of beta power and computed the proportion of windows falling into both sets (Fig. 8e, inset). The alpha-beta ratio was computed per window as alpha power divided by beta power (Fig. 8e).

To test whether the mu rhythm waveform is symmetric around zero, we first bandpass-filtered the single-trial data between 7 and 29 Hz using a second-order Butterworth filter, then identified peaks and troughs of the filtered signal.

The symmetry index was computed as

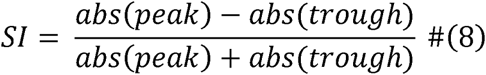

[30] and compared between the empirical MEG data and the HNN simulation.

## Code availability

Code to reproduce all figures in this manuscript is shared at https://github.com/jonescompneurolab/hnn-tuning.

## Funding

This work was supported by NIH R01EB022889, U24NS129945, and P50MH109429. Computational resources were supported by the NIH S10 instrumentation grant S10OD036341 (High-Performance Compute Cluster for Brain Science) via the Center for Computation and Visualization (CCV) at Brown University.

## Declaration of Interests

S.R.J. and N.T. hold equity in PyraSim Neuroscience, maintain director and officer roles within the company, and are listed as inventors on patent applications owned by Brown University which describe translational use cases of the Human Neocortical Neurosolver software. HNN as a software is distributed with an open-source license, and its use is not restricted by company activities or patent applications.

## Declaration of generative AI and AI-assisted technologies in the manuscript preparation process

During the preparation of this work, the author(s) used Claude Opus 5.5 for code review and text editing. The author(s) reviewed and edited the output as needed and take full responsibility for the content of the published article.

## Supplementary Figures

**Supp. Fig. 1.**
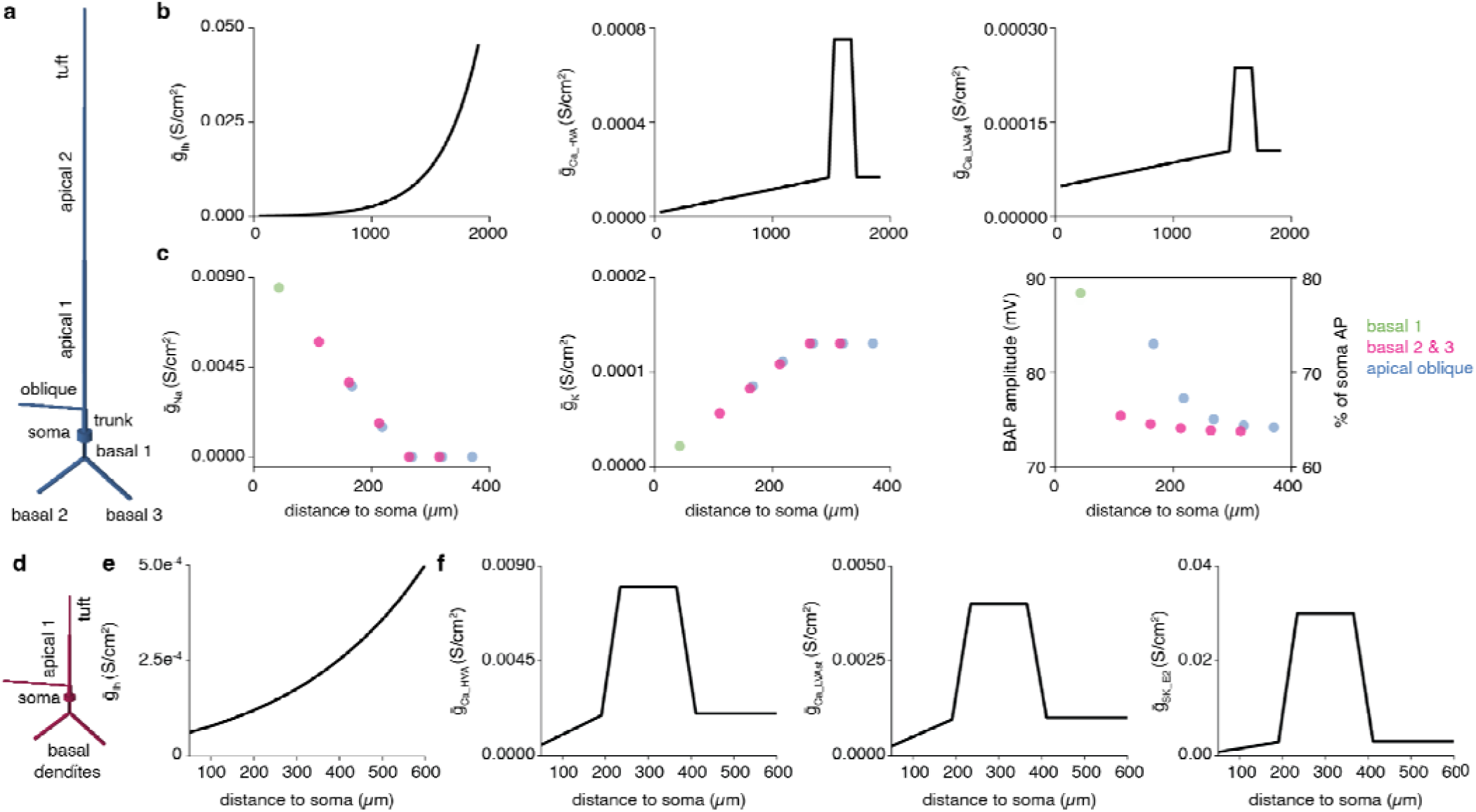
**a** Morphology of L5 extratelencephalic-projecting (ET) pyramidal cell model. **b** (left) Conductance of HCN/Ih channel increases exponentially along apical dendrite (ref ref). (middle, right) The conductance of high- and low-voltage activated Ca2+ channels increases linearly along apical dendrite (see Yuste), with a “hotzone” in the apical tuft. **c** Gradient in N+ (left) and K+ (middle) channel density in the basal dendrites. The non-uniform distribution causes an attenuation of backpropagating action potentials from soma to the tip of the basal dendrite. **d** Morphology of L2/3 model. e (left) Exponential distribution of the HCN channels. **f** Non-uniform distribution of the high- and low-voltage activated Ca2+ channels and Ca2+-activated potassium channels (SK-E2) was inspired by the distribution in the L5 pyramidal cell and tuned to produce dendritic Ca2+ spikes in the L2/3 pyramidal model.

**Supp. Fig. 2.**
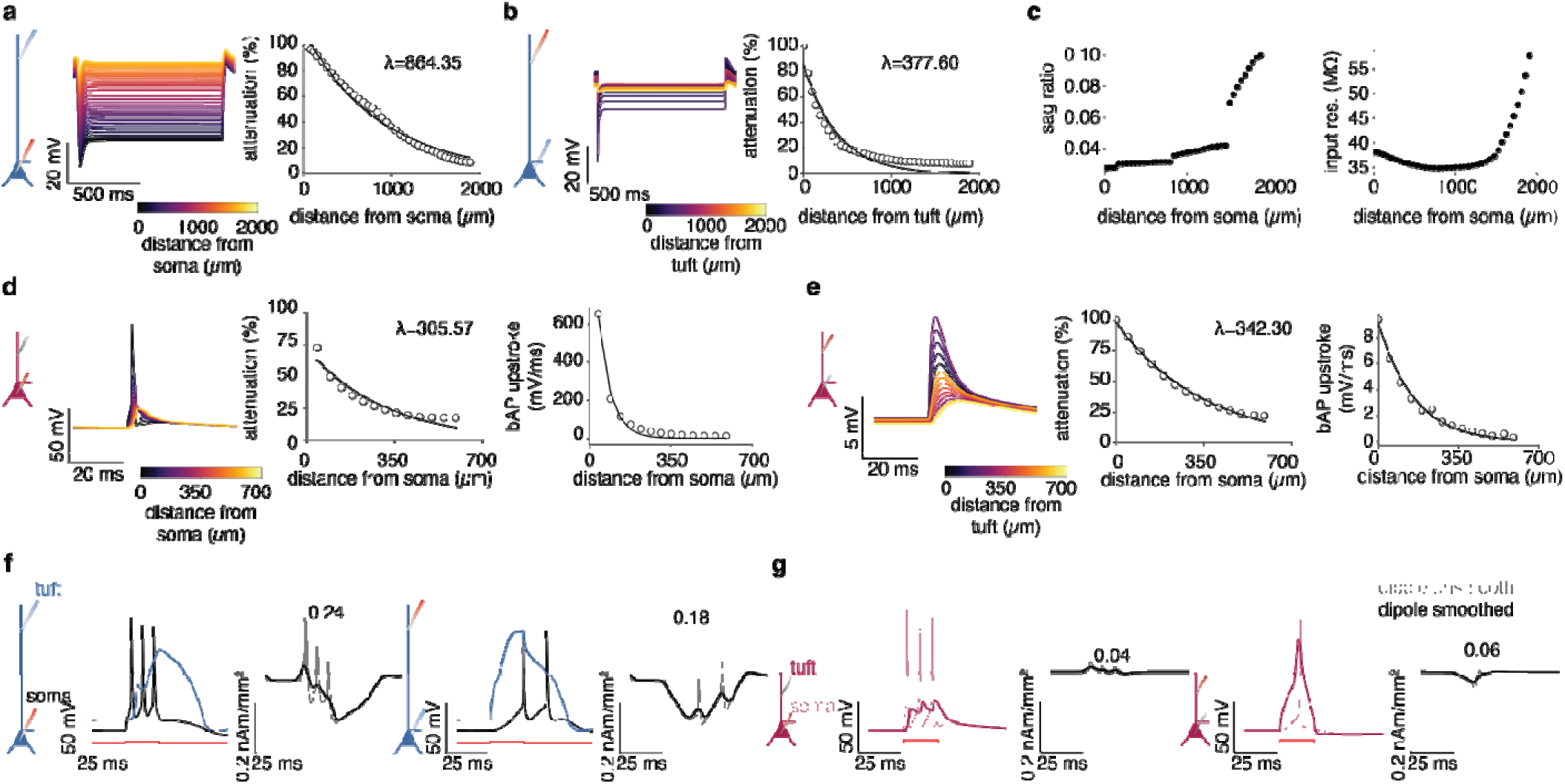
**a, b** Attenuation of hyperpolarization signals along the L5 pyramidal dendrite, length constant λ identified with a single exponential fit. **c** The pyramidal neuron model shows an exponential increase in sag ratio (left) and input resistance (right) with distance from soma. **d** Attenuation of somatic action potentials along the dendrite of the L2/3 neuron. **e** Attenuation of dendritic inputs from tuft to soma. **f** (left) A 20 ms current injection at the soma of a single L5ET neuron triggers high-frequency bursting at the soma and a Ca2+ spike at the tuft. The associated dipole has a moment of about 0.24 nA/mm2. (right) Current injection into the tuft of the pyramidal neuron leads to a Ca2+ spike and burst at the soma, again generating a dipole with a moment of about 0.18 nA/mm2. **g** Experiments in f reproduced for a single L2/3 pyramidal neuron. (left) Current injection into the soma leads to action potentials but does not trigger Ca2+ spikes. The associated dipole demonstrates that L2/3 pyramidal neurons generate smaller dipole moments due to their smaller size (about 0.02-0.04 nA/mm2). (right) Current injection into the dendrite of the L2/3 pyramidal cell generates a Ca2+ spike which similarly to the L5ET neuron is associated with a downward dipole, whereby the moment is again smaller than for the L5ET neuron.

**Supp. Fig. 3.**
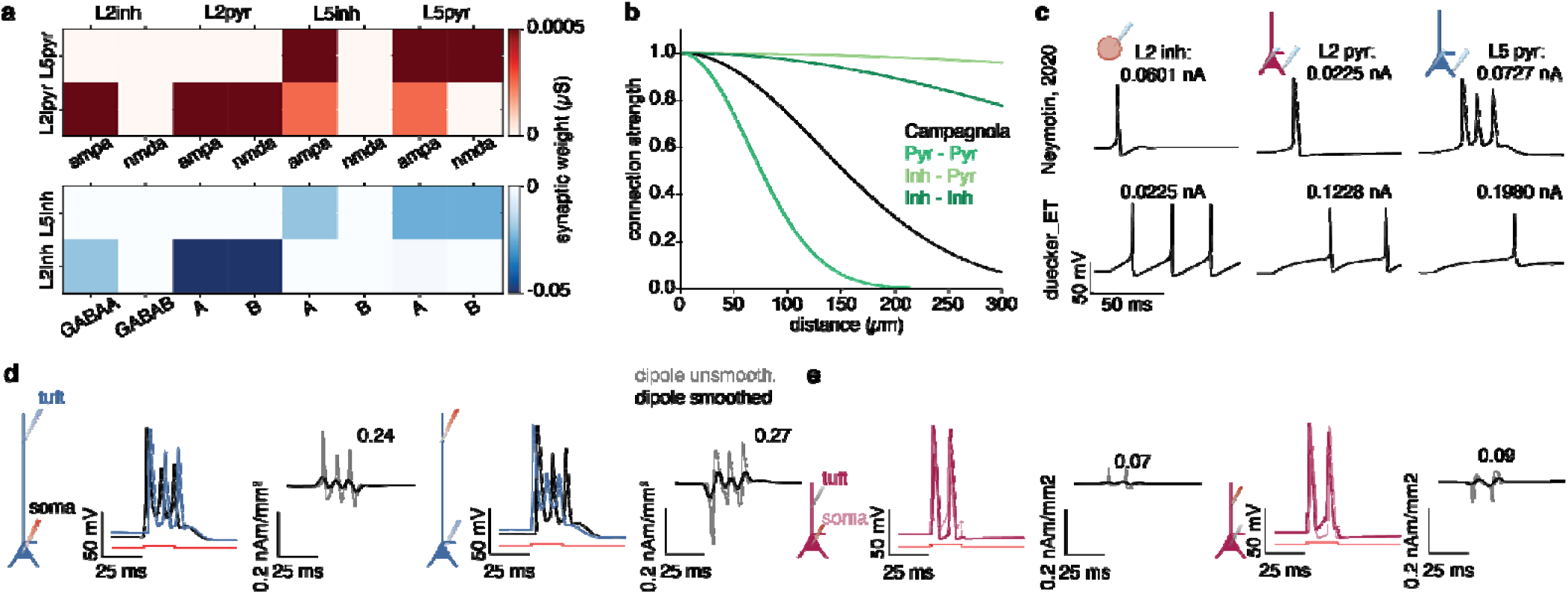
Connectivity, excitability, and dipoles in original HNN model. **a** Comparison to Supp. Fig. 2f-g shows that the maximum excitatory synaptic weights are higher than in the duecker_ET_model, while inhibitory synapses are weaker. **b** The space constant of the pyramidal - pyramidal connections is lower, while inhibitory connections are wider, effectively connecting each interneuron in the circuit to each pyramidal cell. **c** The excitability of each cell type in the Neymotin 2020 model and the duecker_ET_model. The neurons in the Neymotin model have a similar excitability, while the interneurons in the duecker_ET_model are several-fold more excitable. **d** Membrane potentials at the tuft and soma, showing bursting behavior but not Ca2+ spike. As a result, the amplitude of the smoothed dipole is lower than for the duecker_ET_model. **e** Same es d, for the L2/3 pyramidal neuron. The model does not show an a tenuation of signals between dendrites and soma.

## Supplementary Tables

**Supp. Table 1.** Comparison of parameters of the exogenous proximal and distal drives sequence used to generate ERP as published in Neymotin et al., 2020 (white background) and in duecker_ET_model (gray background). # defines the number of spikes each drive cell generates per drive.

|  | μ |  | σ |  | # |  | AMPA (S/cm2) |  | NMDA (S/cm2) |  | delay | Scale factor |  | λ |  | RMSE |  |
|---|---|---|---|---|---|---|---|---|---|---|---|---|---|---|---|---|---|
| p1 | 19.64 | 15.59 | 2.52 | 2.79 | 1 | 1 | L2i: 0.2<br>L2p:0.25<br>L5i: 0.41<br>L5p:0.01 | 0.008<br>0.019<br>7e-15<br>0.014 | - | 0.012<br>0.029<br>1e-15<br>0.005 | 0.1<br>0.1<br>1.0<br>1.0 | 3000 | 1000 | 3 | 3 | 15.5 | 10.82 |
| d | 65.69 | 59.19 | 3.81 | 4.45 | 1 | 4 | L2i: 0.06<br>L2p:1.15<br>L5p:1 | 3e-6<br>1e-4<br>1.36 | - | 1e-12<br>8e-5<br>0.016 | 0.1<br>0.1<br>0.1 |  |  |  |  |  |  |
| p2 | 90.53 | 122.49 | 10.38 | 14.42 | 1 | 3 | L2i: 6e-5<br>L2p:48.2<br>L5i: 0.02<br>L5p:49.9 | 1e-15<br>0.015<br>2e-7<br>0.3 | - | 1e-15<br>2e-5<br>7e-8<br>0.001 | 0.1<br>0.1<br>1.0<br>1. |  |  |  |  |  |  |

**Supp. Table 2.** Comparison parameters of exogenous subthreshold bursty proximal and distal inputs to generate the alpha/beta complex of the somatosensory mu rhythm, as published in Neymotin et al., 2020 (white background) and in duecker_ET_model (gray background).

| Model | prox_burst_std | dist_burst_std | prox_dist_delay | prox_weigts_ampa | dist_weights_ampa | prox_delay | dist_delay | scale_factor |
|---|---|---|---|---|---|---|---|---|
| Neymotin, 2020 | 20 | 20 | 0 (but see synaptic delays) | 5e-5 | 5e-5x4 | L2bsk: 0.1<br>L2pyr: 0.1<br>L5bsk: 1<br>Lpyr: 1 | L2bsk: 5<br>L2pyr: 5<br>Lpyr: 5 | 300000 |
| duecker_ET_model | 20 | 15 | 5 | 2.5e-4 | 2.5e-4x4 | L2bsk: 0.1<br>L2pyr: 0.1<br>L5bsk: 0.1<br>Lpyr: 0.1 | L2bsk: 0.1<br>L2pyr: 0.1<br>Lpyr: 0.1 | 2000 |

**Supp. Table 3.** Input drives and connectivity sustained vs bursty gamma in Neymotin 2020 model and duecker_ET_model (gray).

| PING | Model | Inh - Pyr |  | Inh - Inh |  | Pyr - Inh |  | Poiss Drive freq | Poisson weights AMPA |  | Poisson weights NMDA |  |
|  |  | L2/3 | L5 | L2/3 | L5 | L2/3 | L5 |  | L2/3 | L5 | L2/3 | L5 |
|---|---|---|---|---|---|---|---|---|---|---|---|---|
| sustained | ney | 7e-3 | 8e-2 | 1e-2 | 7.5e-3 | 1.2e-3 | 9.14e-4 | 140 | 8e-4 | 7.5e-3 | 0 | 0 |
|  | due | 1.5e-2 | 1.5e-2 | 5e-4 | 5e-4 | 2e-3 | 2e-3 | 200 | 4.5e-3 | 5.5e-3 | 4.5e-3 | 5.5e-3 |
| bursty | ney | 7e-3 | 8e-2 | 1e-2 | 7.5e-3 | 1.2e-3 | 9.14e-4 | 4 (L2/3) 5 (L5) | 8e-4 | 1e-4 | 7.5e-3 | 1e-3 |
|  | due | 5e-3 | 5e-3 | 5e-4 | 5e-4 | 2e-3 | 2e-3 | 100 | 3.5e-3 | 5.5e-3 | 2.5e-3 | 3.5e-3 |

## Notes

### Summary of Updates

Box Fig 1 was updated for aesthetics and a more plausible representation of dipolar signals.

## References

[1] S. D. Antic, R. M. Empson, and T. Knöpfel, ‘Voltage imaging to understand connections and functions of neuronal circuits’, J. Neurophysiol., vol. 116, no. 1, pp. 135–152, Jul. 2016, doi: 10.1152/jn.00226.2016.

[2] E. S. Boyden, F. Zhang, E. Bamberg, G. Nagel, and K. Deisseroth, ‘Millisecond-timescale, genetically targeted optical control of neural activity’, Nat. Neurosci., vol. 8, no. 9, pp. 1263–1268, Sep. 2005, doi: 10.1038/nn1525.

[3] M. Robbins, C. N. Christensen, C. F. Kaminski, and M. Zlatic, ‘Calcium imaging analysis – how far have we come?’, F1000Research, vol. 10, p. 258, Aug. 2021, doi: 10.12688/f1000research.51755.2.

[4] D. Badre, M. J. Frank, and C. I. Moore, ‘Interactionist Neuroscience’, Neuron, vol. 88, no. 5, pp. 855–860, Dec. 2015, doi: 10.1016/j.neuron.2015.10.021.

[5] N. C. Rust, ‘Elusive cures: why neuroscience hasn’t solved brain disorders—and how we can change that’, in Elusive Cures, Princeton University Press, 2025.

[6] F. Lopes da Silva, ‘EEG and MEG: Relevance to Neuroscience’, Neuron, vol. 80, no. 5, pp. 1112–1128, Dec. 2013, doi: 10.1016/j.neuron.2013.10.017.

[7] F. Maestú, P. Cuesta, O. Hasan, A. Fernandéz, M. Funke, and P. E. Schulz, ‘The Importance of the Validation of M/EEG With Current Biomarkers in Alzheimer’s Disease’, Front. Hum. Neurosci., vol. 13, Feb. 2019, doi: 10.3389/fnhum.2019.00017.

[8] V. Sakkalis, ‘Applied Strategies Towards EEG/MEG Biomarker Identification in Clinical and Cognitive Research’, Biomark. Med., vol. 5, no. 1, pp. 93–105, Feb. 2011, doi: 10.2217/bmm.10.121.

[9] R. Csaky, M. W. J. van Es, O. P. Jones, and M. Woolrich, ‘Interpretable many-class decoding for MEG’, NeuroImage, vol. 282, p. 120396, Nov. 2023, doi: 10.1016/j.neuroimage.2023.120396.

[10] S. d’Ascoli et al., ‘Towards decoding individual words from non-invasive brain recordings’, Nat. Commun., vol. 16, no. 1, p. 10521, Nov. 2025, doi: 10.1038/s41467-025-65499-0.

[11] J. Lévy et al., ‘Noninvasive decoding of typed sentences from human brain activity’, Nat. Neurosci., vol. 29, no. 10, pp. 2438–2444, Oct. 2026, doi: 10.1038/s41593-026-02303-2.

[12] J. Lévy, H. Banville, J. Rapin, J.-R. King, T. Moreau, and S. d’Ascoli, ‘DANCE: Detect and Classify Events in EEG’, May 11, 2026, arXiv: arXiv:2605.10688. doi: 10.48550/arXiv.2605.10688.

[13] F. Magnabosco and O. Hauk, ‘Decoding Semantics from Dynamic Brain Activation Patterns: From Trials to Task in EEG/MEG Source Space’, eNeuro, vol. 11, no. 3, Mar. 2024, doi: 10.1523/ENEURO.0277-23.2023.

[14] G. Halnes, T. V. Ness, S. Næss, E. Hagen, K. H. Pettersen, and G. T. Einevoll, Electric Brain Signals: Foundations and Applications of Biophysical Modeling. Cambridge: Cambridge University Press, 2024. doi: 10.1017/9781009039826.

[15] M. Hämäläinen, R. Hari, R. J. Ilmoniemi, J. Knuutila, and O. V. Lounasmaa, ‘Magnetoencephalography---theory, instrumentation, and applications to noninvasive studies of the working human brain’, Rev. Mod. Phys., vol. 65, no. 2, pp. 413–497, Apr. 1993, doi: 10.1103/RevModPhys.65.413.

[16] S. Næss et al., ‘Biophysically detailed forward modeling of the neural origin of EEG and MEG signals’, NeuroImage, vol. 225, p. 117467, Jan. 2021, doi: 10.1016/j.neuroimage.2020.117467.

[17] G. Buzsáki, C. A. Anastassiou, and C. Koch, ‘The origin of extracellular fields and currents — EEG, ECoG, LFP and spikes’, Nat. Rev. Neurosci., vol. 13, no. 6, pp. 407–420, Jun. 2012, doi: 10.1038/nrn3241.

[18] H. Lindén, K. H. Pettersen, and G. T. Einevoll, ‘Intrinsic dendritic filtering gives low-pass power spectra of local field potentials’, J. Comput. Neurosci., vol. 29, no. 3, pp. 423–444, Dec. 2010, doi: 10.1007/s10827-010-0245-4.

[19] S. Murakami, T. Zhang, A. Hirose, and Y. C. Okada, ‘Physiological origins of evoked magnetic fields and extracellular field potentials produced by guinea-pig CA3 hippocampal slices’, J. Physiol., vol. 544, no. 1, pp. 237–251, 2002, doi: 10.1113/jphysiol.2002.027094.

[20] S. Murakami and Y. Okada, ‘Contributions of principal neocortical neurons to magnetoencephalography and electroencephalography signals’, J. Physiol., vol. 575, no. 3, pp. 925–936, 2006, doi: 10.1113/jphysiol.2006.105379.

[21] R. Grech et al., ‘Review on solving the inverse problem in EEG source analysis’, J. NeuroEngineering Rehabil., vol. 5, no. 1, p. 25, Nov. 2008, doi: 10.1186/1743-0003-5-25.

[22] H. T. A. Moore and D. Pinal, ‘From Scalp to Cortex: A Practical Guide to EEG Source Localization and Overview of Its Methods, Challenges, and Applications’, in *Research Methods for Biological Psychology*, S. Galdo-Álvarez, Ed., New York, NY: Springer US, 2026, pp. 143–168. doi: 10.1007/978-1-0716-5340-1_5.

[23] S. Baillet, ‘Forward and Inverse Problems of MEG/EEG’, in *Encyclopedia of Computational Neuroscience*, D. Jaeger and R. Jung, Eds, New York, NY: Springer New York, 2013, pp. 1–8. doi: 10.1007/978-1-4614-7320-6_529-1.

[24] J. Gross, J. Kujala, M. Hämäläinen, L. Timmermann, A. Schnitzler, and R. Salmelin, ‘Dynamic imaging of coherent sources: Studying neural interactions in the human brain’, Proc. Natl. Acad. Sci., vol. 98, no. 2, pp. 694–699, Jan. 2001, doi: 10.1073/pnas.98.2.694.

[25] B. Veen, J. Joseph, and K. Hecox, ‘Localization of intra-cerebral sources of electrical activity via linearly constrained minimum variance spatial filtering’, 1992 IEEE 6th SP Workshop Stat. Signal Array Process. SSAP 1992 - Conf. Proc., vol. 44, pp. 526–529, 1992, doi: 10.1109/SSAP.1992.246899.

[26] Y. Attal and D. Schwartz, ‘Assessment of subcortical source localization using deep brain activity imaging model with minimum norm operators: a MEG study’, PloS One, vol. 8, no. 3, p. e59856, 2013, doi: 10.1371/journal.pone.0059856.

[27] B. Zoefel, O. Abbasi, J. Gross, and S. A. Kotz, ‘Entrainment echoes in the cerebellum’, Proc. Natl. Acad. Sci., vol. 121, no. 34, p. e2411167121, Aug. 2024, doi: 10.1073/pnas.2411167121.

[28] M. Jas et al., ‘HNN-core: A Python software for cellular and circuit-level interpretation of human MEG/EEG’, J. Open Source Softw., vol. 8, no. 92, p. 5848, 2023, doi: 10.21105/joss.05848.

[29] S. A. Neymotin et al., ‘Human Neocortical Neurosolver (HNN), a new software tool for interpreting the cellular and network origin of human MEG/EEG data’, eLife, vol. 9, p. e51214, Jan. 2020, doi: 10.7554/eLife.51214.

[30] S. R. Jones, D. L. Pritchett, M. A. Sikora, S. M. Stufflebeam, M. Hämäläinen, and C. I. Moore, ‘Quantitative Analysis and Biophysically Realistic Neural Modeling of the MEG Mu Rhythm: Rhythmogenesis and Modulation of Sensory-Evoked Responses’, J. Neurophysiol., vol. 102, no. 6, pp. 3554–3572, Dec. 2009, doi: 10.1152/jn.00535.2009.

[31] S. R. Jones, D. L. Pritchett, S. M. Stufflebeam, M. Hämäläinen, and C. I. Moore, ‘Neural Correlates of Tactile Detection: A Combined Magnetoencephalography and Biophysically Based Computational Modeling Study’, J. Neurosci., vol. 27, no. 40, pp. 10751–10764, Oct. 2007, doi: 10.1523/JNEUROSCI.0482-07.2007.

[32] C. Fernandez Pujol, J. Bruce, R. V. Thorpe, S. R. Jones, and A. R. Dykstra, ‘Neurophysiology of mismatch negativity generation: a biophysical modeling study’, Nov. 09, 2025, bioRxiv: The Preprint Server for Biology: 2025.11.08.687373. doi: 10.1101/2025.11.08.687373.

[33] C. Fernandez Pujol, E. G. Blundon, and A. R. Dykstra, ‘Laminar specificity of the auditory perceptual awareness negativity: A biophysical modeling study’, PLOS Comput. Biol., vol. 19, no. 6, p. e1011003, Jun. 2023, doi: 10.1371/journal.pcbi.1011003.

[34] C. Kohl, T. Parviainen, and S. R. Jones, ‘Neural Mechanisms Underlying Human Auditory Evoked Responses Revealed By Human Neocortical Neurosolver’, Brain Topogr., vol. 35, no. 1, pp. 19–35, Jan. 2022, doi: 10.1007/s10548-021-00838-0.

[35] K. Lankinen, J. Ahveninen, M. Jas, T. Raij, and S. P. Ahlfors, ‘Neuronal Modeling of Cross-Sensory Visual Evoked Magnetoencephalography Responses in the Auditory Cortex’, J. Neurosci., vol. 44, no. 17, Apr. 2024, doi: 10.1523/JNEUROSCI.1119-23.2024.

[36] J. J. Bonaiuto, S. Little, S. A. Neymotin, S. R. Jones, G. R. Barnes, and S. Bestmann, ‘Laminar dynamics of high amplitude beta bursts in human motor cortex’, Neuroimage, vol. 242, p. 118479, Nov. 2021, doi: 10.1016/j.neuroimage.2021.118479.

[37] M. J. Szul et al., ‘Diverse beta burst waveform motifs characterize movement-related cortical dynamics’, Prog. Neurobiol., vol. 228, p. 102490, Sep. 2023, doi: 10.1016/j.pneurobio.2023.102490.

[38] D. A. Diesburg, J. R. Wessel, and S. R. Jones, ‘Biophysical Modeling of Frontocentral ERP Generation Links Circuit-Level Mechanisms of Action-Stopping to a Behavioral Race Model’, J. Neurosci. Off. J. Soc. Neurosci., vol. 44, no. 20, p. e2016232024, May 2024, doi: 10.1523/JNEUROSCI.2016-23.2024.

[39] M. A. Sherman et al., ‘Neural mechanisms of transient neocortical beta rhythms: Converging evidence from humans, computational modeling, monkeys, and mice’, Proc. Natl. Acad. Sci., vol. 113, no. 33, pp. E4885–E4894, Aug. 2016, doi: 10.1073/pnas.1604135113.

[40] Allen Institute for Brain Science, ‘Allen Cell Types Database -- Human Morphology-Electrophysiology’. Allen Institute for Brain Science, 2015. [Online]. Available: https://celltypes.brain-map.org/data

[41] L. Campagnola et al., ‘Local connectivity and synaptic dynamics in mouse and human neocortex’, Science, vol. 375, no. 6585, p. eabj5861, Mar. 2022, doi: 10.1126/science.abj5861.

[42] R. Dalley, et al., ‘Human Neocortical Glutamatergic Neurons Revealed Through Multimodal Profiling’, Jan. 16, 2026, bioRxiv: The Preprint Server for Biology: 2026.01.08.698257. doi: 10.64898/2026.01.08.698257.

[43] B. E. Kalmbach et al., ‘Signature morpho-electric, transcriptomic, and dendritic properties of human layer 5 neocortical pyramidal neurons’, Neuron, vol. 109, no. 18, pp. 2914–2927.e5, Sep. 2021, doi: 10.1016/j.neuron.2021.08.030.

[44] G. Komlósi, G. Molnár, M. Rózsa, S. Oláh, P. Barzó, and G. Tamás, ‘Fluoxetine (Prozac) and Serotonin Act on Excitatory Synaptic Transmission to Suppress Single Layer 2/3 Pyramidal Neuron-Triggered Cell Assemblies in the Human Prefrontal Cortex’, J. Neurosci., vol. 32, no. 46, pp. 16369–16378, Nov. 2012, doi: 10.1523/JNEUROSCI.2618-12.2012.

[45] G. Molnár et al., ‘Complex Events Initiated by Individual Spikes in the Human Cerebral Cortex’, PLOS Biol., vol. 6, no. 9, p. e222, Sep. 2008, doi: 10.1371/journal.pbio.0060222.

[46] S. C. Seeman et al., ‘Sparse recurrent excitatory connectivity in the microcircuit of the adult mouse and human cortex’, eLife, vol. 7, p. e37349, Sep. 2018, doi: 10.7554/eLife.37349.

[47] V. Szegedi et al., ‘Plasticity in Single Axon Glutamatergic Connection to GABAergic Interneurons Regulates Complex Events in the Human Neocortex’, PLOS Biol., vol. 14, no. 11, p. e2000237, Nov. 2016, doi: 10.1371/journal.pbio.2000237.

[48] R. G. Law et al., ‘Thalamocortical Mechanisms Regulating the Relationship between Transient Beta Events and Human Tactile Perception’, Cereb. Cortex, vol. 32, no. 4, pp. 668–688, Feb. 2022, doi: 10.1093/cercor/bhab221.

[49] S. Baillet, J. C. Mosher, and R. M. Leahy, ‘Electromagnetic brain mapping’, IEEE Signal Process. Mag., vol. 18, no. 6, pp. 14–30, Nov. 2001, doi: 10.1109/79.962275.

[50] O. Hauk, ‘Keep it simple: a case for using classical minimum norm estimation in the analysis of EEG and MEG data’, NeuroImage, vol. 21, no. 4, pp. 1612–1621, Apr. 2004, doi: 10.1016/j.neuroimage.2003.12.018.

[51] P. L. Nunez, M. D. Nunez, and R. Srinivasan, ‘Multi-Scale Neural Sources of EEG: Genuine, Equivalent, and Representative. A Tutorial Review’, Brain Topogr., vol. 32, no. 2, pp. 193–214, Mar. 2019, doi: 10.1007/s10548-019-00701-3.

[52] A. E. Rimehaug et al., ‘Uncovering circuit mechanisms of current sinks and sources with biophysical simulations of primary visual cortex’, eLife, vol. 12, p. e87169, Jul. 2023, doi: 10.7554/eLife.87169.

[53] P. C. Bush and T. J. Sejnowski, ‘Reduced compartmental models of neocortical pyramidal cells’, J. Neurosci. Methods, vol. 46, no. 2, pp. 159–166, Feb. 1993, doi: 10.1016/0165-0270(93)90151-G.

[54] D. J. Felleman and D. C. Van Essen, ‘Distributed hierarchical processing in the primate cerebral cortex’, Cereb. Cortex N. Y. N 1991, vol. 1, no. 1, pp. 1–47, Feb. 1991, doi: 10.1093/cercor/1.1.1-a.

[55] A. T. Kulics and L. J. Cauller, ‘Cerebral cortical somatosensory evoked responses, multiple unit activity and current source-densities: their interrelationships and significance to somatic sensation as revealed by stimulation of the awake monkey’s hand’, Exp. Brain Res., vol. 62, no. 1, pp. 46–60, Mar. 1986, doi: 10.1007/BF00237402.

[56] N. N. Peterson, C. E. Schroeder, and J. C. Arezzo, ‘Neural generators of early cortical somatosensory evoked potentials in the awake monkey’, Electroencephalogr. Clin. Neurophysiol. Potentials Sect., vol. 96, no. 3, pp. 248–260, May 1995, doi: 10.1016/0168-5597(95)00006-E.

[57] C. E. Schroeder, A. D. Mehta, and S. J. Givre, ‘A spatiotemporal profile of visual system activation revealed by current source density analysis in the awake macaque.’, Cereb. Cortex, vol. 8, no. 7, pp. 575–592, Oct. 1998, doi: 10.1093/cercor/8.7.575.

[58] C. M. Constantinople and R. M. Bruno, ‘Deep Cortical Layers Are Activated Directly by Thalamus’, Science, vol. 340, no. 6140, pp. 1591–1594, Jun. 2013, doi: 10.1126/science.1236425.

[59] J. A. Harris et al., ‘Hierarchical organization of cortical and thalamic connectivity’, Nature, vol. 575, no. 7781, pp. 195–202, Nov. 2019, doi: 10.1038/s41586-019-1716-z.

[60] N. Kim, S. Bahn, J. H. Choi, J. S. Kim, and J.-C. Rah, ‘Synapses from the Motor Cortex and a High-Order Thalamic Nucleus are Spatially Clustered in Proximity to Each Other in the Distal Tuft Dendrites of Mouse Somatosensory Cortex’, Cereb. Cortex, vol. 32, no. 4, pp. 737–754, Feb. 2022, doi: 10.1093/cercor/bhab236.

[61] G. Minamisawa, S. E. Kwon, M. Chevée, S. P. Brown, and D. H. O’Connor, ‘A Non-canonical Feedback Circuit for Rapid Interactions between Somatosensory Cortices’, Cell Rep., vol. 23, no. 9, pp. 2718–2731.e6, May 2018, doi: 10.1016/j.celrep.2018.04.115.

[62] S. Lee and S. R. Jones, ‘Distinguishing mechanisms of gamma frequency oscillations in human current source signals using a computational model of a laminar neocortical network’, Front. Hum. Neurosci., vol. 7, 2013, doi: 10.3389/fnhum.2013.00869.

[63] N. Tolley, P. L. C. Rodrigues, A. Gramfort, and S. R. Jones, ‘Methods and considerations for estimating parameters in biophysically detailed neural models with simulation based inference’, PLOS Comput. Biol., vol. 20, no. 2, p. e1011108, Feb. 2024, doi: 10.1371/journal.pcbi.1011108.

[64] R. V. Thorpe, C. J. Black, D. A. Borton, L. Hu, C. Y. Saab, and S. R. Jones, ‘Distinct neocortical mechanisms underlie human SI responses to median nerve and laser-evoked peripheral activation’, Imaging Neurosci., vol. 2, pp. 1–29, Feb. 2024, doi: 10.1162/imag_a_00095.

[65] F. Helmchen, K. Svoboda, W. Denk, and D. W. Tank, ‘In vivo dendritic calcium dynamics in deep-layer cortical pyramidal neurons’, Nat. Neurosci., vol. 2, no. 11, pp. 989–996, Nov. 1999, doi: 10.1038/14788.

[66] H. G. Kim and B. W. Connors, ‘Apical dendrites of the neocortex: correlation between sodium- and calcium-dependent spiking and pyramidal cell morphology’, J. Neurosci., vol. 13, no. 12, pp. 5301–5311, Dec. 1993, doi: 10.1523/JNEUROSCI.13-12-05301.1993.

[67] M. E. Larkum, T. Nevian, M. Sandler, A. Polsky, and J. Schiller, ‘Synaptic Integration in Tuft Dendrites of Layer 5 Pyramidal Neurons: A New Unifying Principle’, Science, vol. 325, no. 5941, pp. 756–760, 2009.

[68] M. E. Larkum, J. J. Zhu, and B. Sakmann, ‘Dendritic mechanisms underlying the coupling of the dendritic with the axonal action potential initiation zone of adult rat layer 5 pyramidal neurons’, J. Physiol., vol. 533, no. 2, pp. 447–466, 2001, doi: 10.1111/j.1469-7793.2001.0447a.x.

[69] M. E. Larkum, J. J. Zhu, and B. Sakmann, ‘A new cellular mechanism for coupling inputs arriving at different cortical layers’, Nature, vol. 398, no. 6725, pp. 338–341, Mar. 1999, doi: 10.1038/18686.

[70] M. E. Larkum, K. M. M. Kaiser, and B. Sakmann, ‘Calcium electrogenesis in distal apical dendrites of layer 5 pyramidal cells at a critical frequency of back-propagating action potentials’, Proc. Natl. Acad. Sci., vol. 96, no. 25, pp. 14600–14604, Dec. 1999, doi: 10.1073/pnas.96.25.14600.

[71] S. Ramaswamy and H. Markram, ‘Anatomy and physiology of the thick-tufted layer 5 pyramidal neuron’, Front. Cell. Neurosci., vol. 9, Jun. 2015, doi: 10.3389/fncel.2015.00233.

[72] R. Yuste, M. J. Gutnick, D. Saar, K. R. Delaney, and D. W. Tank, ‘Ca2+ accumulations in dendrites of neocortical pyramidal neurons: An apical band and evidence for two functional compartments’, Neuron, vol. 13, no. 1, pp. 23–43, Jul. 1994, doi: 10.1016/0896-6273(94)90457-X.

[73] A. Baker et al., ‘Specialized Subpopulations of Deep-Layer Pyramidal Neurons in the Neocortex: Bridging Cellular Properties to Functional Consequences’, J. Neurosci., vol. 38, no. 24, pp. 5441–5455, Jun. 2018, doi: 10.1523/JNEUROSCI.0150-18.2018.

[74] A. M. Hattox and S. B. Nelson, ‘Layer V Neurons in Mouse Cortex Projecting to Different Targets Have Distinct Physiological Properties’, J. Neurophysiol., vol. 98, no. 6, pp. 3330–3340, Dec. 2007, doi: 10.1152/jn.00397.2007.

[75] N. L. Jorstad et al., ‘Transcriptomic cytoarchitecture reveals principles of human neocortex organization’, Science, vol. 382, no. 6667, p. eadf6812, Oct. 2023, doi: 10.1126/science.adf6812.

[76] M. Suzuki and M. E. Larkum, ‘Dendritic calcium spikes are clearly detectable at the cortical surface’, Nat. Commun., vol. 8, no. 1, p. 276, Aug. 2017, doi: 10.1038/s41467-017-00282-4.

[77] E. Hay, S. Hill, F. Schürmann, H. Markram, and I. Segev, ‘Models of Neocortical Layer 5b Pyramidal Cells Capturing a Wide Range of Dendritic and Perisomatic Active Properties’, PLOS Comput. Biol., vol. 7, no. 7, p. e1002107, Jul. 2011, doi: 10.1371/journal.pcbi.1002107.

[78] M. E. Larkum, ‘A cellular mechanism for cortical associations: an organizing principle for the cerebral cortex’, Trends Neurosci., vol. 36, no. 3, pp. 141–151, Mar. 2013, doi: 10.1016/j.tins.2012.11.006.

[79] N. Takahashi, C. Ebner, J. Sigl-Glöckner, S. Moberg, S. Nierwetberg, and M. E. Larkum, ‘Active dendritic currents gate descending cortical outputs in perception’, Nat. Neurosci., vol. 23, no. 10, pp. 1277–1285, Oct. 2020, doi: 10.1038/s41593-020-0677-8.

[80] N. Takahashi, T. G. Oertner, P. Hegemann, and M. E. Larkum, ‘Active cortical dendrites modulate perception’, Science, vol. 354, no. 6319, pp. 1587–1590, Dec. 2016, doi: 10.1126/science.aah6066.

[81] M. Suzuki and M. E. Larkum, ‘General Anesthesia Decouples Cortical Pyramidal Neurons’, Cell, vol. 180, no. 4, pp. 666–676.e13, Feb. 2020, doi: 10.1016/j.cell.2020.01.024.

[82] E. Maristany de las Casas et al., ‘Tuft dendrites in frontal motor cortex enable flexible learning’, Science, vol. 392, no. 6798, p. eadx4358, May 2026, doi: 10.1126/science.adx4358.

[83] M. E. Larkum and J. J. Zhu, ‘Signaling of layer 1 and whisker-evoked Ca2+ and Na+ action potentials in distal and terminal dendrites of rat neocortical pyramidal neurons in vitro and in vivo’, J. Neurosci. Off. J. Soc. Neurosci., vol. 22, no. 16, pp. 6991–7005, Aug. 2002, doi: 10.1523/JNEUROSCI.22-16-06991.2002.

[84] L. Beaulieu-Laroche et al., ‘Allometric rules for mammalian cortical layer 5 neuron biophysics’, Nature, vol. 600, no. 7888, pp. 274–278, Dec. 2021, doi: 10.1038/s41586-021-04072-3.

[85] L. Beaulieu-Laroche et al., ‘Enhanced Dendritic Compartmentalization in Human Cortical Neurons’, Cell, vol. 175, no. 3, pp. 643–651.e14, Oct. 2018, doi: 10.1016/j.cell.2018.08.045.

[86] S. Murakami and Y. Okada, ‘Invariance in current dipole moment density across brain structures and species: Physiological constraint for neuroimaging’, NeuroImage, vol. 111, pp. 49–58, May 2015, doi: 10.1016/j.neuroimage.2015.02.003.

[87] D. M. Iascone et al., ‘Whole-Neuron Synaptic Mapping Reveals Spatially Precise Excitatory/Inhibitory Balance Limiting Dendritic and Somatic Spiking’, Neuron, vol. 106, no. 4, pp. 566–578.e8, May 2020, doi: 10.1016/j.neuron.2020.02.015.

[88] J. Berg et al., ‘Human neocortical expansion involves glutamatergic neuron diversification’, Nature, vol. 598, no. 7879, pp. 151–158, Oct. 2021, doi: 10.1038/s41586-021-03813-8.

[89] A. Gidon et al., ‘Dendritic action potentials and computation in human layer 2/3 cortical neurons’, Science, vol. 367, no. 6473, pp. 83–87, Jan. 2020, doi: 10.1126/science.aax6239.

[90] H. M. Gooch et al., ‘High-fidelity dendritic sodium spike generation in human layer 2/3 neocortical pyramidal neurons’, Cell Rep., vol. 41, no. 3, p. 111500, Oct. 2022, doi: 10.1016/j.celrep.2022.111500.

[91] B. E. Kalmbach et al., ‘h-Channels Contribute to Divergent Intrinsic Membrane Properties of Supragranular Pyramidal Neurons in Human versus Mouse Cerebral Cortex’, Neuron, vol. 100, no. 5, pp. 1194–1208.e5, Dec. 2018, doi: 10.1016/j.neuron.2018.10.012.

[92] B. R. Lee et al., ‘Signature morphoelectric properties of diverse GABAergic interneurons in the human neocortex’, Science, vol. 382, no. 6667, p. eadf6484, Oct. 2023, doi: 10.1126/science.adf6484.

[93] A. Cumberland, N. Hale, A. Azhan, C. P. Gilchrist, G. Chincarini, and M. Tolcos, ‘Excitatory and inhibitory neuron imbalance in the intrauterine growth restricted fetal guinea pig brain: Relevance to the developmental origins of schizophrenia and autism’, Dev. Neurobiol., vol. 83, no. 1–2, pp. 40–53, 2023, doi: 10.1002/dneu.22907.

[94] D. Golomb, K. Donner, L. Shacham, D. Shlosberg, Y. Amitai, and D. Hansel, ‘Mechanisms of firing patterns in fast-spiking cortical interneurons’, PLoS Comput. Biol., vol. 3, no. 8, p. e156, Aug. 2007, doi: 10.1371/journal.pcbi.0030156.

[95] P. P. Gao et al., ‘Local glutamate-mediated dendritic plateau potentials change the state of the cortical pyramidal neuron’, J. Neurophysiol., vol. 125, no. 1, pp. 23–42, Jan. 2021, doi: 10.1152/jn.00734.2019.

[96] C. E. Jahr and C. F. Stevens, ‘A quantitative description of NMDA receptor-channel kinetic behavior’, J. Neurosci., vol. 10, no. 6, pp. 1830–1837, Jun. 1990, doi: 10.1523/JNEUROSCI.10-06-01830.1990.

[97] C. E. Jahr and C. F. Stevens, ‘Voltage dependence of NMDA-activated macroscopic conductances predicted by single-channel kinetics’, J. Neurosci. Off. J. Soc. Neurosci., vol. 10, no. 9, pp. 3178–3182, Sep. 1990, doi: 10.1523/JNEUROSCI.10-09-03178.1990.

[98] A. Papoutsi, G. Kastellakis, and P. Poirazi, ‘Basal tree complexity shapes functional pathways in the prefrontal cortex’, J. Neurophysiol., vol. 118, no. 4, pp. 1970–1983, Oct. 2017, doi: 10.1152/jn.00099.2017.

[99] J. M. Schulz, F. Knoflach, M.-C. Hernandez, and J. Bischofberger, ‘Dendrite-targeting interneurons control synaptic NMDA-receptor activation via nonlinear α5-GABAA receptors’, Nat. Commun., vol. 9, no. 1, p. 3576, Sep. 2018, doi: 10.1038/s41467-018-06004-8.

[100] A. Destexhe, T. Bal, D. A. McCormick, and T. J. Sejnowski, ‘Ionic mechanisms underlying synchronized oscillations and propagating waves in a model of ferret thalamic slices’, J. Neurophysiol., vol. 76, no. 3, pp. 2049–2070, Sep. 1996, doi: 10.1152/jn.1996.76.3.2049.

[101] S. A. Neymotin, S. Dura-Bernal, P. Lakatos, T. D. Sanger, and W. W. Lytton, ‘Multitarget Multiscale Simulation for Pharmacological Treatment of Dystonia in Motor Cortex’, Front. Pharmacol., vol. 7, p. 157, Jun. 2016, doi: 10.3389/fphar.2016.00157.

[102] A. Destexhe, Z. F. Mainen, T. J. Sejnowski, and others, ‘Kinetic models of synaptic transmission’, Methods Neuronal Model., vol. 2, pp. 1–25, 1998.

[103] J. Schiller, G. Major, H. J. Koester, and Y. Schiller, ‘NMDA spikes in basal dendrites of cortical pyramidal neurons’, Nature, vol. 404, no. 6775, pp. 285–289, Mar. 2000, doi: 10.1038/35005094.

[104] H. Hu, J. Gan, and P. Jonas, ‘Fast-spiking, parvalbumin+ GABAergic interneurons: From cellular design to microcircuit function’, Science, vol. 345, no. 6196, p. 1255263, Aug. 2014, doi: 10.1126/science.1255263.

[105] H. Markram et al., ‘Reconstruction and Simulation of Neocortical Microcircuitry’, Cell, vol. 163, no. 2, pp. 456–492, Oct. 2015, doi: 10.1016/j.cell.2015.09.029.

[106] H. Markram, M. Toledo-Rodriguez, Y. Wang, A. Gupta, G. Silberberg, and C. Wu, ‘Interneurons of the neocortical inhibitory system’, Nat. Rev. Neurosci., vol. 5, no. 10, pp. 793–807, Oct. 2004, doi: 10.1038/nrn1519.

[107] J. Obermayer et al., ‘Lateral inhibition by Martinotti interneurons is facilitated by cholinergic inputs in human and mouse neocortex’, Nat. Commun., vol. 9, no. 1, 2018, doi: 10.1038/s41467-018-06628-w.

[108] K. A. Wilmes, H. Sprekeler, and S. Schreiber, ‘Inhibition as a Binary Switch for Excitatory Plasticity in Pyramidal Neurons’, PLOS Comput. Biol., vol. 12, no. 3, p. e1004768, Mar. 2016, doi: 10.1371/journal.pcbi.1004768.

[109] S. Oláh et al., ‘Regulation of cortical microcircuits by unitary GABA-mediated volume transmission’, Nature, vol. 461, no. 7268, pp. 1278–1281, Oct. 2009, doi: 10.1038/nature08503.

[110] L. M. Palmer, J. M. Schulz, S. C. Murphy, D. Ledergerber, M. Murayama, and M. E. Larkum, ‘The Cellular Basis of GABAB-Mediated Interhemispheric Inhibition’, Science, vol. 335, no. 6071, pp. 989–993, Feb. 2012, doi: 10.1126/science.1217276.

[111] B. Schuman, S. Dellal, A. Prönneke, R. Machold, and B. Rudy, ‘Neocortical Layer 1: An Elegant Solution to Top-Down and Bottom-Up Integration’, Annu. Rev. Neurosci., vol. 44, pp. 221–252, 2021, doi: 10.1146/annurev-neuro-100520-012117.

[112] E. Pérez-Garci, M. E. Larkum, and T. Nevian, ‘Inhibition of dendritic Ca2+ spikes by GABAB receptors in cortical pyramidal neurons is mediated by a direct Gi/o-βγ-subunit interaction with Cav1 channels’, J. Physiol., vol. 591, no. 7, pp. 1599–1612, 2013, doi: 10.1113/jphysiol.2012.245464.

[113] Y. N. Billeh et al., ‘Systematic Integration of Structural and Functional Data into Multi-scale Models of Mouse Primary Visual Cortex’, Neuron, vol. 106, no. 3, pp. 388–403.e18, May 2020, doi: 10.1016/j.neuron.2020.01.040.

[114] A. L. Bodor et al., ‘The synaptic architecture of layer 5 thick tufted excitatory neurons in mouse visual cortex’, Nat. Neurosci., vol. 28, no. 8, pp. 1704–1715, Aug. 2025, doi: 10.1038/s41593-025-02004-2.

[115] S. Lefort, C. Tomm, J.-C. F. Sarria, and C. C. H. Petersen, ‘The Excitatory Neuronal Network of the C2 Barrel Column in Mouse Primary Somatosensory Cortex’, Neuron, vol. 61, no. 2, pp. 301–316, Jan. 2009, doi: 10.1016/j.neuron.2008.12.020.

[116] A. Pala and C. C. H. Petersen, ‘In Vivo Measurement of Cell-Type-Specific Synaptic Connectivity and Synaptic Transmission in Layer 2/3 Mouse Barrel Cortex’, Neuron, vol. 85, no. 1, pp. 68–75, Jan. 2015, doi: 10.1016/j.neuron.2014.11.025.

[117] I. Ferezou, S. Bolea, and C. C. H. Petersen, ‘Visualizing the Cortical Representation of Whisker Touch: Voltage-Sensitive Dye Imaging in Freely Moving Mice’, Neuron, vol. 50, no. 4, pp. 617–629, May 2006, doi: 10.1016/j.neuron.2006.03.043.

[118] C. P. J. de Kock et al., ‘High-frequency burst spiking in layer 5 thick-tufted pyramids of rat primary somatosensory cortex encodes exploratory touch’, *Commun*. Biol., vol. 4, no. 1, pp. 1–14, Jun. 2021, doi: 10.1038/s42003-021-02241-8.

[119] J. Freeman, C. M. Ziemba, D. J. Heeger, E. P. Simoncelli, and J. A. Movshon, ‘A functional and perceptual signature of the second visual area in primates’, Nat. Neurosci., vol. 16, no. 7, pp. 974–981, Jul. 2013, doi: 10.1038/nn.3402.

[120] M. Niwa, J. S. Johnson, K. N. O’Connor, and M. L. Sutter, ‘Activity related to perceptual judgment and action in primary auditory cortex’, J. Neurosci. Off. J. Soc. Neurosci., vol. 32, no. 9, pp. 3193–3210, Feb. 2012, doi: 10.1523/JNEUROSCI.0767-11.2012.

[121] X. Wang, ‘Cortical Coding of Auditory Features’, Annu. Rev. Neurosci., vol. 41, pp. 527–552, Jul. 2018, doi: 10.1146/annurev-neuro-072116-031302.

[122] M. Quiquempoix, S. L. Fayad, K. Boutourlinsky, N. Leresche, R. C. Lambert, and T. Bessaih, ‘Layer 2/3 Pyramidal Neurons Control the Gain of Cortical Output’, Cell Rep., vol. 24, no. 11, pp. 2799–2807.e4, Sep. 2018, doi: 10.1016/j.celrep.2018.08.038.

[123] S. R. Jones, ‘When brain rhythms aren’t “rhythmic”: implication for their mechanisms and meaning’, Curr. Opin. Neurobiol., vol. 40, pp. 72–80, Oct. 2016, doi: 10.1016/j.conb.2016.06.010.

[124] L. Power and T. Bardouille, ‘Age-related trends in the cortical sources of transient beta bursts during a sensorimotor task and rest’, NeuroImage, vol. 245, p. 118670, Dec. 2021, doi: 10.1016/j.neuroimage.2021.118670.

[125] J. A. Cardin et al., ‘Driving fast-spiking cells induces gamma rhythm and controls sensory responses’, Nature, vol. 459, pp. 663–667, 2009, doi: 10.1038/nature0800.

[126] D. Pantazis, M. Fang, S. Qin, Y. Mohsenzadeh, Q. Li, and R. M. Cichy, ‘Decoding the orientation of contrast edges from MEG evoked and induced responses’, NeuroImage, vol. 180, pp. 267–279, Oct. 2018, doi: 10.1016/j.neuroimage.2017.07.022.

[127] M. E. Larkum, ‘Are Dendrites Conceptually Useful?’, Neuroscience, vol. 489, pp. 4–14, May 2022, doi: 10.1016/j.neuroscience.2022.03.008.

[128] S. R. Williams and G. J. Stuart, ‘Dependence of EPSP Efficacy on Synapse Location in Neocortical Pyramidal Neurons’, Science, vol. 295, no. 5561, pp. 1907–1910, Mar. 2002, doi: 10.1126/science.1067903.

[129] S. Tasciotti et al., ‘Clustered inputs engage dendritic nonlinearities and calcium signaling to support efficient place-field formation in CA1 pyramidal neurons’, Cell Rep., vol. 45, no. 8, Aug. 2026, doi: 10.1016/j.celrep.2026.117793.

[130] R. K. Wong, D. A. Prince, and A. I. Basbaum, ‘Intradendritic recordings from hippocampal neurons’, Proc. Natl. Acad. Sci. U. S. A., vol. 76, no. 2, pp. 986–990, Feb. 1979, doi: 10.1073/pnas.76.2.986.

[131] N. L. Golding, H. Y. Jung, T. Mickus, and N. Spruston, ‘Dendritic calcium spike initiation and repolarization are controlled by distinct potassium channel subtypes in CA1 pyramidal neurons’, J. Neurosci. Off. J. Soc. Neurosci., vol. 19, no. 20, Oct. 1999, doi: 10.1523/JNEUROSCI.19-20-08789.1999.

[132] J. Cichon and W.-B. Gan, ‘Branch-specific dendritic Ca2+ spikes cause persistent synaptic plasticity’, Nature, vol. 520, no. 7546, pp. 180–185, Apr. 2015, doi: 10.1038/nature14251.

[133] J. B. Isbister et al., ‘Modeling and simulation of neocortical micro- and mesocircuitry (Part II, Physiology and experimentation)’, eLife, vol. 13, p. RP99693, Jan. 2026, doi: 10.7554/eLife.99693.

[134] H. Lindén et al., ‘Modeling the Spatial Reach of the LFP’, Neuron, vol. 72, no. 5, pp. 859–872, Dec. 2011, doi: 10.1016/j.neuron.2011.11.006.

[135] M. W. Reimann et al., ‘Modeling and Simulation of Neocortical Micro- and Mesocircuitry. Part I: Anatomy’, eLife, vol. 13, Aug. 2024, doi: 10.7554/eLife.99688.1.

[136] A. E. Rimehaug, A. M. Dale, A. Arkhipov, and G. T. Einevoll, ‘Uncovering population contributions to the extracellular potential in the mouse visual system using Laminar Population Analysis’, PLOS Comput. Biol., vol. 20, no. 12, p. e1011830, Dec. 2024, doi: 10.1371/journal.pcbi.1011830.

[137] S. Dura-Bernal et al., ‘Multiscale model of primary motor cortex circuits predicts in vivo cell-type-specific, behavioral state-dependent dynamics’, Cell Rep., vol. 42, no. 6, p. 112574, Jun. 2023, doi: 10.1016/j.celrep.2023.112574.

[138] S. Dura-Bernal et al., ‘Data-driven multiscale model of macaque auditory thalamocortical circuits reproduces in vivo dynamics’, Cell Rep., vol. 42, no. 11, p. 113378, Nov. 2023, doi: 10.1016/j.celrep.2023.113378.

[139] A. Guet-McCreight et al., ‘In-silico testing of new pharmacology for restoring inhibition and human cortical function in depression’, *Commun*. Biol., vol. 7, no. 1, p. 225, Feb. 2024, doi: 10.1038/s42003-024-05907-1.

[140] A. Guet-McCreight and E. Hay, ‘Preserving spiking and EEG in detailed models of human cortical microcircuits with morphological reductions’, Sep. 24, 2026, bioRxiv. doi: 10.64898/2026.09.18.752744.

[141] E. Hagen, S. Næss, T. V. Ness, and G. T. Einevoll, ‘Multimodal Modeling of Neural Network Activity: Computing LFP, ECoG, EEG, and MEG Signals With LFPy 2.0’, *Front*. Neuroinformatics, vol. 12, Dec. 2018, doi: 10.3389/fninf.2018.00092.

[142] J. Tharayil, J. B. Isbister, E. Neufeld, and M. Reimann, ‘Computational modeling reveals biological mechanisms underlying the whisker flick EEG’, iScience, vol. 28, no. 11, p. 113793, Nov. 2025, doi: 10.1016/j.isci.2025.113793.

[143] H. K. Yao et al., ‘Reduced inhibition in depression impairs stimulus processing in human cortical microcircuits’, Cell Rep., vol. 38, no. 2, Jan. 2022, doi: 10.1016/j.celrep.2021.110232.

[144] K. Dai et al., ‘Brain Modeling ToolKit: An open source software suite for multiscale modeling of brain circuits’, PLOS Comput. Biol., vol. 16, no. 11, p. e1008386, Nov. 2020, doi: 10.1371/journal.pcbi.1008386.

[145] S. Dura-Bernal et al., ‘NetPyNE, a tool for data-driven multiscale modeling of brain circuits’, eLife, vol. 8, p. e44494, Apr. 2019, doi: 10.7554/eLife.44494.

[146] M. L. Hines and N. T. Carnevale, ‘The NEURON simulation environment’, Neural Comput., vol. 9, no. 6, pp. 1179–1209, Aug. 1997, doi: 10.1162/neco.1997.9.6.1179.

[147] N. Tolley et al., ‘A Protocol For Uncovering Neural Mechanisms Of Neurotherapeutic Effects On Electroencephalography Using The Human Neocortical Neurosolver’, J. Vis. Exp. JoVE, no. 231, p. e70618, May 2026, doi: 10.3791/70618.

[148] K. H. Pettersen and G. T. Einevoll, ‘Amplitude variability and extracellular low-pass filtering of neuronal spikes’, Biophys. J., vol. 94, no. 3, pp. 784–802, Feb. 2008, doi: 10.1529/biophysj.107.111179.

[149] A. Rosenblueth and N. Wiener, ‘The Role of Models in Science’, Philos. Sci., vol. 12, no. 4, pp. 316–321, 1945.

[150] T. V. Ness, M. W. H. Remme, and G. T. Einevoll, ‘h-Type Membrane Current Shapes the Local Field Potential from Populations of Pyramidal Neurons’, J. Neurosci., vol. 38, no. 26, pp. 6011–6024, Jun. 2018, doi: 10.1523/JNEUROSCI.3278-17.2018.

[151] T. V. Ness, M. W. H. Remme, and G. T. Einevoll, ‘Active subthreshold dendritic conductances shape the local field potential’, J. Physiol., vol. 594, no. 13, pp. 3809–3825, Jul. 2016, doi: 10.1113/JP272022.

[152] I. N. Beloozerova, M. G. Sirota, H. A. Swadlow, G. N. Orlovsky, L. B. Popova, and T. G. Deliagina, ‘Activity of Different Classes of Neurons of the Motor Cortex during Postural Corrections’, J. Neurosci., vol. 23, no. 21, pp. 7844–7853, Aug. 2003, doi: 10.1523/JNEUROSCI.23-21-07844.2003.

[153] C. P. J. de Kock, R. M. Bruno, H. Spors, and B. Sakmann, ‘Layer- and cell-type-specific suprathreshold stimulus representation in rat primary somatosensory cortex’, J. Physiol., vol. 581, no. Pt 1, pp. 139–154, May 2007, doi: 10.1113/jphysiol.2006.124321.

[154] S. Sakata and K. D. Harris, ‘Laminar Structure of Spontaneous and Sensory-Evoked Population Activity in Auditory Cortex’, Neuron, vol. 64, no. 3, pp. 404–418, Nov. 2009, doi: 10.1016/j.neuron.2009.09.020.

[155] T. E. Bakken et al., ‘Comparative cellular analysis of motor cortex in human, marmoset and mouse’, Nature, vol. 598, no. 7879, pp. 111–119, Oct. 2021, doi: 10.1038/s41586-021-03465-8.

[156] F. Scala et al., ‘Layer 4 of mouse neocortex differs in cell types and circuit organization between sensory areas’, Nat. Commun., vol. 10, no. 1, p. 4174, Sep. 2019, doi: 10.1038/s41467-019-12058-z.

[157] J. F. Staiger, I. Flagmeyer, D. Schubert, K. Zilles, R. Kötter, and H. J. Luhmann, ‘Functional diversity of layer IV spiny neurons in rat somatosensory cortex: quantitative morphology of electrophysiologically characterized and biocytin labeled cells’, Cereb. Cortex, vol. 14, no. 6, pp. 690–701, Jun. 2004, doi: 10.1093/cercor/bhh029.

[158] N. Yamawaki, K. Borges, B. A. Suter, K. D. Harris, and G. M. G. Shepherd, ‘A genuine layer 4 in motor cortex with prototypical synaptic circuit connectivity’, eLife, vol. 3, p. e05422, 2014, doi: 10.7554/eLife.05422.

[159] P. H. Smith and L. C. Populin, ‘Fundamental differences between the thalamocortical recipient layers of the cat auditory and visual cortices’, J. Comp. Neurol., vol. 436, no. 4, pp. 508–519, 2001, doi: 10.1002/cne.1084.

[160] S. Haegens et al., ‘Laminar Profile and Physiology of the α Rhythm in Primary Visual, Auditory, and Somatosensory Regions of Neocortex’, J. Neurosci., vol. 35, no. 42, pp. 14341–14352, Oct. 2015, doi: 10.1523/JNEUROSCI.0600-15.2015.

[161] L. J. Gentet, G. J. Stuart, and J. D. Clements, ‘Direct measurement of specific membrane capacitance in neurons’, Biophys. J., vol. 79, no. 1, pp. 314–320, Jul. 2000, doi: 10.1016/S0006-3495(00)76293-X.

[162] G. Stuart and N. Spruston, ‘Determinants of voltage attenuation in neocortical pyramidal neuron dendrites’, J. Neurosci. Off. J. Soc. Neurosci., vol. 18, no. 10, pp. 3501–3510, May 1998, doi: 10.1523/JNEUROSCI.18-10-03501.1998.

[163] O. Amsalem et al., ‘An efficient analytical reduction of detailed nonlinear neuron models’, Nat. Commun., vol. 11, p. 288, Jan. 2020, doi: 10.1038/s41467-019-13932-6.

[164] C. M. Colbert and E. Pan, ‘Ion channel properties underlying axonal action potential initiation in pyramidal neurons’, Nat. Neurosci., vol. 5, no. 6, pp. 533–538, Jun. 2002, doi: 10.1038/nn0602-857.

[165] J. Magistretti and A. Alonso, ‘Biophysical properties and slow voltage-dependent inactivation of a sustained sodium current in entorhinal cortex layer-II principal neurons: a whole-cell and single-channel study’, J. Gen. Physiol., vol. 114, no. 4, pp. 491–509, Oct. 1999, doi: 10.1085/jgp.114.4.491.

[166] A. Korngreen and B. Sakmann, ‘Voltage-gated K+ channels in layer 5 neocortical pyramidal neurones from young rats: subtypes and gradients’, J. Physiol., vol. 525, no. Pt 3, pp. 621–639, Jun. 2000, doi: 10.1111/j.1469-7793.2000.00621.x.

[167] P. R. Adams, D. A. Brown, and A. Constanti, ‘M-currents and other potassium currents in bullfrog sympathetic neurones’, J. Physiol., vol. 330, pp. 537–572, Sep. 1982, doi: 10.1113/jphysiol.1982.sp014357.

[168] R. B. Avery and D. Johnston, ‘Multiple Channel Types Contribute to the Low-Voltage-Activated Calcium Current in Hippocampal CA3 Pyramidal Neurons’, J. Neurosci., vol. 16, no. 18, pp. 5567–5582, Sep. 1996, doi: 10.1523/JNEUROSCI.16-18-05567.1996.

[169] A. D. Randall and R. W. Tsien, ‘Contrasting biophysical and pharmacological properties of T-type and R-type calcium channels’, Neuropharmacology, vol. 36, no. 7, pp. 879–893, Jul. 1997, doi: 10.1016/s0028-3908(97)00086-5.

[170] I. Reuveni, A. Friedman, Y. Amitai, and M. J. Gutnick, ‘Stepwise repolarization from Ca2+ plateaus in neocortical pyramidal cells: evidence for nonhomogeneous distribution of HVA Ca2+ channels in dendrites’, J. Neurosci., vol. 13, no. 11, pp. 4609–4621, Nov. 1993, doi: 10.1523/JNEUROSCI.13-11-04609.1993.

[171] M. H. P. Kole, S. Hallermann, and G. J. Stuart, ‘Single Ih Channels in Pyramidal Neuron Dendrites: Properties, Distribution, and Impact on Action Potential Output’, J. Neurosci., vol. 26, no. 6, pp. 1677–1687, Feb. 2006, doi: 10.1523/JNEUROSCI.3664-05.2006.

[172] J. Rettig et al., ‘Characterization of a Shaw-related potassium channel family in rat brain.’, EMBO J., vol. 11, no. 7, pp. 2473–2486, Jul. 1992, doi: 10.1002/j.1460-2075.1992.tb05312.x.

[173] M. Köhler et al., ‘Small-Conductance, Calcium-Activated Potassium Channels from Mammalian Brain’, Science, vol. 273, no. 5282, pp. 1709–1714, Sep. 1996, doi: 10.1126/science.273.5282.1709.

[174] H. Ishibuchi, N. Tsukamoto, and Y. Nojima, ‘Evolutionary many-objective optimization: A short review’, in 2008 IEEE Congress on Evolutionary Computation (IEEE World Congress on Computational Intelligence), Jun. 2008, pp. 2419–2426. doi: 10.1109/CEC.2008.4631121.

[175] V. Khare, X. Yao, and K. Deb, ‘Performance Scaling of Multi-objective Evolutionary Algorithms’, in Evolutionary Multi-Criterion Optimization, C. M. Fonseca, P. J. Fleming, E. Zitzler, L. Thiele, and K. Deb, Eds, Berlin, Heidelberg: Springer, 2003, pp. 376–390. doi: 10.1007/3-540-36970-8_27.

[176] B. Li, J. Li, K. Tang, and X. Yao, ‘Many-Objective Evolutionary Algorithms: A Survey’, ACM Comput. Surv. CSUR, vol. 48, no. 1, p. 13:1–13:35, Sep. 2015, doi: 10.1145/2792984.

[177] R. C. Purshouse and P. J. Fleming, ‘On the Evolutionary Optimization of Many Conflicting Objectives’, IEEE Trans. Evol. Comput., vol. 11, no. 6, pp. 770–784, Dec. 2007, doi: 10.1109/TEVC.2007.910138.

[178] A. S. Shai, C. A. Anastassiou, M. E. Larkum, and C. Koch, ‘Physiology of Layer 5 Pyramidal Neurons in Mouse Primary Visual Cortex: Coincidence Detection through Bursting’, PLOS Comput. Biol., vol. 11, no. 3, p. e1004090, Mar. 2015, doi: 10.1371/journal.pcbi.1004090.

[179] C. D. Acker and S. D. Antic, ‘Quantitative Assessment of the Distributions of Membrane Conductances Involved in Action Potential Backpropagation Along Basal Dendrites’, J. Neurophysiol., vol. 101, no. 3, pp. 1524–1541, Mar. 2009, doi: 10.1152/jn.00651.2007.

[180] T. Nevian, M. E. Larkum, A. Polsky, and J. Schiller, ‘Properties of basal dendrites of layer 5 pyramidal neurons: a direct patch-clamp recording study’, Nat. Neurosci., vol. 10, no. 2, pp. 206–214, Feb. 2007, doi: 10.1038/nn1826.

[181] S. Rudolph and M. S. Thanawala, ‘Location matters: somatic and dendritic SK channels answer to distinct calcium signals’, J. Neurophysiol., vol. 114, no. 1, pp. 1–5, Jul. 2015, doi: 10.1152/jn.00181.2014.

[182] C. A. Sailer, W. A. Kaufmann, J. Marksteiner, and H.-G. Knaus, ‘Comparative immunohistochemical distribution of three small-conductance Ca2+-activated potassium channel subunits, SK1, SK2, and SK3 in mouse brain’, Mol. Cell. Neurosci., vol. 26, no. 3, pp. 458–469, Jul. 2004, doi: 10.1016/j.mcn.2004.03.002.

[183] J. L. Maciaszek, H. Soh, R. S. Walikonis, A. V. Tzingounis, and G. Lykotrafitis, ‘Topography of Native SK Channels Revealed by Force Nanoscopy in Living Neurons’, J. Neurosci., vol. 32, no. 33, pp. 11435–11440, Aug. 2012, doi: 10.1523/JNEUROSCI.1785-12.2012.

[184] V. Szegedi et al., ‘HCN channels at the cell soma ensure the rapid electrical reactivity of fast-spiking interneurons in human neocortex’, PLOS Biol., vol. 21, no. 2, p. e3002001, Feb. 2023, doi: 10.1371/journal.pbio.3002001.

[185] E. M. Callaway et al., ‘A multimodal cell census and atlas of the mammalian primary motor cortex’, Nature, vol. 598, no. 7879, pp. 86–102, Oct. 2021, doi: 10.1038/s41586-021-03950-0.

[186] J. DeFelipe, ‘The Evolution of the Brain, the Human Nature of Cortical Circuits, and Intellectual Creativity’, Front. Neuroanat., vol. 5, May 2011, doi: 10.3389/fnana.2011.00029.

[187] S. H. Hendry, H. D. Schwark, E. G. Jones, and J. Yan, ‘Numbers and proportions of GABA-immunoreactive neurons in different areas of monkey cerebral cortex’, J. Neurosci. Off. J. Soc. Neurosci., vol. 7, no. 5, pp. 1503–1519, May 1987, doi: 10.1523/JNEUROSCI.07-05-01503.1987.

[188] P. L. Gabbott and S. J. Bacon, ‘Local circuit neurons in the medial prefrontal cortex (areas 24a,b,c, 25 and 32) in the monkey: II. Quantitative areal and laminar distributions’, J. Comp. Neurol., vol. 364, no. 4, pp. 609–636, Jan. 1996, doi: 10.1002/(SICI)1096-9861(19960122)364:4<609::AID-CNE2>3.0.CO;2-7.

[189] M. F. Glasser and D. C. V. Essen, ‘Mapping Human Cortical Areas In Vivo Based on Myelin Content as Revealed by T1- and T2-Weighted MRI’, J. Neurosci., vol. 31, no. 32, pp. 11597–11616, Aug. 2011, doi: 10.1523/JNEUROSCI.2180-11.2011.

[190] D. S. Koh, J. R. Geiger, P. Jonas, and B. Sakmann, ‘Ca(2+)-permeable AMPA and NMDA receptor channels in basket cells of rat hippocampal dentate gyrus.’, J. Physiol., vol. 485, no. 2, pp. 383–402, 1995, doi: 10.1113/jphysiol.1995.sp020737.

[191] B. Kriener, H. Hu, and K. Vervaeke, ‘Parvalbumin interneuron dendrites enhance gamma oscillations’, Cell Rep., vol. 39, no. 11, p. 110948, Jun. 2022, doi: 10.1016/j.celrep.2022.110948.

[192] L. J. Cauller, B. Clancy, and B. W. Connors, ‘Backward cortical projections to primary somatosensory cortex in rats extend long horizontal axons in layer I’, J. Comp. Neurol., vol. 390, no. 2, pp. 297–310, 1998, doi: 10.1002/(SICI)1096-9861(19980112)390:2<297::AID-CNE11>3.0.CO;2-V.

[193] L. J. Cauller and B. W. Connors, ‘Synaptic physiology of horizontal afferents to layer I in slices of rat SI neocortex’, J. Neurosci. Off. J. Soc. Neurosci., vol. 14, no. 2, pp. 751–762, Feb. 1994, doi: 10.1523/JNEUROSCI.14-02-00751.1994.

[194] R. W. Guillery and S. M. Sherman, ‘Thalamic relay functions and their role in corticocortical communication: generalizations from the visual system’, Neuron, vol. 33, no. 2, pp. 163–175, Jan. 2002, doi: 10.1016/s0896-6273(01)00582-7.

[195] N. Hansen, ‘The CMA Evolution Strategy: A Tutorial’, Mar. 10, 2023, arXiv: arXiv:1604.00772. doi: 10.48550/arXiv.1604.00772.

